# Synthetic lethality between TP53 and ENDOD1

**DOI:** 10.1101/2020.10.14.339309

**Authors:** Zizhi Tang, Ming Zeng, Xiaojun Wang, Chang Guo, Peng Yue, Xiaohu Zhang, Huiqiang Lou, Jun Chen, Dezhi Mu, Daochun Kong, Antony M. Carr, Cong Liu

## Abstract

The atypical nuclease ENDOD1 functions with cGAS-STING in innate immunity. Here we identify a previously uncharacterized ENDOD1 function in DNA repair. ENDOD1 is enriched in the nucleus following H_2_O_2_ treatment and *ENDOD1*^-/-^ cells show increased PARP chromatin-association. Loss of ENDOD1 function is synthetic lethal with homologous recombination defects, with affected cells accumulating DNA double strand breaks. Remarkably, we also uncover an additional synthetic lethality between ENDOD1 and p53. ENDOD1 depletion in *TP53* mutated tumour cells, or p53 depletion in *ENDOD1*^-/-^ cells, results in rapid single stranded DNA accumulation and cell death. Because *TP53* is mutated in ∼50% of tumours, ENDOD1 has potential as a wide-spectrum target for synthetic lethal treatments. To support this we demonstrate that systemic knockdown of mouse *EndoD1* is well tolerated and whole-animal siRNA against human *ENDOD1* restrains *TP53* mutated tumour progression in xenograft models. These data identify ENDOD1 as a potential cancer-specific target for SL drug discovery.

## Introduction

The development of PARP inhibitors (PARPi), such as Olaparib^1^ and Talazoparib^2^, to treat BRCA-deficient breast cancer^3,4^ opened up a new therapeutic strategy for cancer subtype-specific chemotherapy, synthetic lethality^5–7^. As our understanding of PARP and its inhibition has developed, the range of cancers considered for PARPi therapy has expanded to include homologous recombination deficient (HRD) tumours. Significant interest in potential new synthetic lethal (SL) targets has led to many drug development programs aiming to identify molecules that inhibit proteins found to be SL with specific genetic backgrounds common to defined tumour subtypes. An ideal SL target would be mutated or silenced in a wide spectrum of tumours.

During a characterisation of PARPi-induced DNA damage responses in the presence of hepatitis B virus oncoprotein HBx expression, which renders cells HRD by sequestering Cullin4-DDB1 and thus depleting CRL4^WDR70^ (ref^8^), we initially identified an increase in endonuclease domain-containing protein 1 (ENDOD1) peptides following six days of Olaparib treatment (Supplementary Figure 1a). This prompted us to characterise a potential role in DNA repair for ENDOD1 in more detail.

## Results

### ENDOD1 functions in DNA single strand break repair

ENDOD1 has previously been identified as a 500 amino acid protein that interacts with RNF26 to modulate the cGAS-STING innate immunity pathway^9^. ENDOD1 contains three C-terminal transmembrane motifs, a single endonuclease domain (residues 49-257) and an N-terminal signal peptide (residues 1-22) (Figure 1a). Upon immunoblotting, both endogenous and ectopically expressed ENDOD1 displayed multiple bands (Supplementary Figure 1b-c). This indicates the presence of multiple isoforms, potentially derived from post-translational processing and/or modification. In addition, upon expression of either C-or N-terminal Flag-tagged constructs, only the C-terminal, but not the N-terminal, construct could be detected with α-Flag. This suggests that the signal peptide is cleaved as predicted (Supplementary Figure 1c). In untreated RPE1 cells, indirect immunofluorescent staining for ENDOD1 was predominantly cytoplasmic, but following high-dose hydrogen peroxide (H_2_O_2_) treatment, we identified the formation of α-ENDOD1 reactive foci in nuclei that are absent in *ENDOD1* knockout RPE1 cells (*ENDOD1*^-/-^), (Figure 1b). This implies that certain forms of ENDOD1 can enter the nucleus and access damaged DNA.

While ENDOD1 was first identified as a cytoplasmic protein^9^, a recent mass spectrometry study identified ENDOD1 peptides in the nucleus^10^. In the context of our identification of damage-induced intra-nuclear ENDOD1 foci, this is consistent with an additional role for ENDOD1 in DNA repair. Indeed, *ENDOD1*^-/-^ cells showed moderate sensitivities to CPT, cisplatin and the G4 inhibitor Cx5461. No detectable sensitivity was observed upon IR or HU treatment (Supplementary Figure 1d). Unexpectedly, *ENDOD1*^-/-^ cells displayed obvious resistance to the single strand break (SSB)-inducing agent H_2_O_2_. A similar H_2_O_2_ resistance was observed for GES-1, a normal gastric epithelial line, following siRNA against *ENDOD1*.

**Figure 1.**
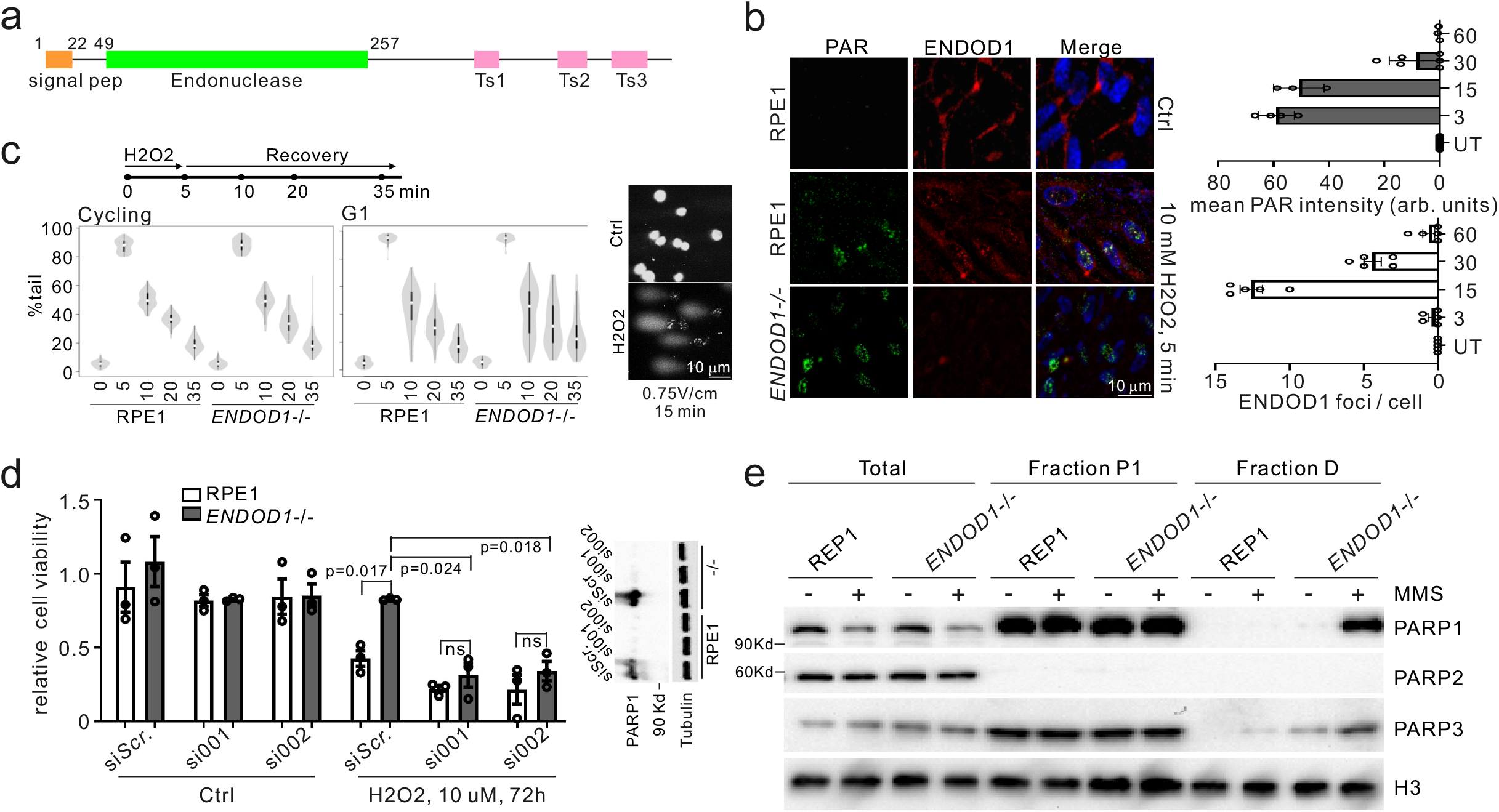
Characterization of ENDOD1 in DNA repair. **a**. Schematic of ENDOD1 protein showing the predicted signal peptide (residues 1-22), the endonuclease domain (residues 49-257) and three C-terminal transmembrane motifs (Ts). **b**. Left: Representative indirect immunofluorescence images for α-ENDOD1 (ABclonal) and α-PAR: untreated RPE1 cells (top row), RPE1 and *ENDOD1*^-/-^ cells after a 5 min treatment with 10 mM H_2_O_2_ (bottom two rows). For the merged panels nuclear DNA was counterstained with DAPI. Right: quantification for nuclear signals at the indicated time points after H_2_O_2_ treatment. arb. units: arbitrary units. n = 3 – 5 biologically independent experiments. **c**. Comet assay to assess repair efficiency. Left: quantification of tail moments from three repeats for proliferating RPE1 and *ENDOD1*^-/-^ cells at the indicated time points after H_2_O_2_ challenge (p = 0.197 for RPE 35 v *ENDOD1*^-/-^ 35, Kruskal test). Middle: tail moments from 150 cells from each of 3 biologically independent experiments for serum-starved G1 arrested RPE1 and *ENDOD1*^-/-^ cells at indicated time points after H_2_O_2_ challenge (p = 0.787 for RPE1 20 v *ENDOD1*^-/-^ 20; p = 0.0000159 for RPE1 35 v *ENDOD1*^-/-^ 35, Kruskal test). Right: representative images of alkaline comet assay. Percentage tail moment was calculated by dividing the pixel intensity of tails by that of heads. White dot: median. Thick whisker: third quartile. Thin whisker: upper/lower adjacent values (1.5x inter-quartile range). **d**. Relative viability of RPE1 or *ENDOD1*^-/-^ cells treated with two different si*PARP1* (si001 and si002) 48 hours before challenge, with or without continuous H_2_O_2_ treatment (10 μM). Assay: CCK8 colorimetry. Inset showing the knockdown efficiency of each siRNA. n = 3 biologically independent experiments. Significance test: two-tailed Student’s *t* test. p values: 0.0057, 0.047. ns: not significant. **e**. Whole cell extract (total), nuclear extracts (P1) and MNase-digested extract (fraction D) from RPE1 or *ENDOD1*^-/-^ cells probed for PARP1, PARP2 and PARP3. Cells were treated with 0.01% MMS for 30 minutes. Histone H3 serves as a control. Representative image of 3 independent experiments.

The kinetics of ENDOD1 foci induced by H_2_O_2_ treatment resembled that of poly(ADP-ribose) (PAR) foci, whose signals appeared prior to those of ENDOD1 (Figure 1b). The focal signals of ENDOD1, which clearly overlap with PAR-positive foci (Supplementary figure 2a), required PARP activity: siRNA knockdown of *PARP1/2*, or treatment with PARPi, diminished the H_2_O_2_-induced ENDOD1 foci (Supplementary Figure 2b). No effect was seen for siRNA knockdown of *PARP3*. Using single cell electrophoresis under denaturing conditions (alkaline comet assay) to assess the repair kinetics of gaps and nicks, we did not detect a major repair defect between proliferating *ENDOD1*^-/-^ and RPE1 control cells (Figure 1c). However, when cells were arrested in G1 phase by serum starvation before treatment, a modest but statistically significant defect was seen in repair kinetics. Consistent with this, despite the clearance of PAR foci in *ENDOD1*^-/-^ being higher in the early phase of repair, PAR foci loss over time was slower in *ENDOD1*^-/-^ serum starved (G1 arrested) cells when compared with control RPE1 cells. We also noted a modest increase of PAR foci in unperturbed serum starved (G1 arrested) *ENDOD1*^-/-^ cells (Supplementary Figure 2c, control untreated lanes). This may represent a low-level of PARP-responsive DNA lesions accumulating in G1 phase that are eventually eliminated before or when cells progress into S/G2. Taken together, these data are consistent with ENDOD1 influencing PARP-dependent SSB repair.

### ENDOD1 influences PARP chromatin association

Interestingly, *PARP1* knockdown with two different siRNAs significantly reversed the increased H_2_O_2_ resistance of *ENDOD1*^-/-^ cells (Figure 1d), whereas knockdown of either *PARP2* or *PARP3* (which plays a minor role in SSB repair^11^) did not affect the resistance (Supplementary Figure 2d). We therefore asked if PARPi could similarly reverse the relative H_2_O_2_ resistance of *ENDOD1*^-/-^ cells. Despite the fact that treating the parental RPE1 cells with a combination of PAPRi and H_2_O_2_ was toxic (Supplementary Figure 2e), the H_2_O_2_ resistance observed in *ENDOD1*^-/-^ cells was not reversed. These distinct outcomes of si*PARP1* and PARPi treatment implies that the protection against oxidative stress caused by *ENDOD1* deletion is attributable to the physical presence of PARP1 *per se*, rather than its enzymatic activity. Indeed, in *ENDOD1*^-/-^ cells 30 minutes after acute treatment with methyl methanesulfonate (MMS) we observed that PARP1 and PARP3 were enriched^12^ in the insoluble histone-containing nuclear fraction (Figure 1e). In contrast to *ENDOD1*^-/-^ cells, control RPE1 cells retained only residual PARP in this fraction. Taken together, we conclude ENDOD1 contributes to prevent excessive PARP association with damaged DNA. It is unclear how loss of ENDOD1 manifests as increased resistance to oxidative stress.

Previous work has identified HR factors, including BRCA1, BRCA2 and Fanconi Anaemia proteins, as being required for DNA repair following PARP inhibition and that compromising the HR pathway results in synthetic lethality with PARPi^3^. To examine how ENDOD1 interplays with homologous recombination factors and PARP we assayed cell survival following siRNA of either *WDR70, BRCA1, BRCA2, ARID1A, BLM, CTIP, CHK1, EXO1, MRE11* or *FANCA* in either *ENDOD1*^-/-^ or control RPE1 cells (Figure 2a). The profile of SL upon depletion of HR factors in *ENDOD1*^-/-^ cells mirrored that for PARP inhibition. Co-depleting *ENDOD1* and *BRCA1* in RPE1 cells using three independent *ENDOD1* siRNA’s showed similar synthetic lethality (Supplementary Figure 3a). In contrast, a range of HR-competent non-cancer cells (including RPE1 and GES-1) exhibited no proliferation defects upon siRNA ablation of *ENDOD1* (Figure 2b).

**Figure 2.**
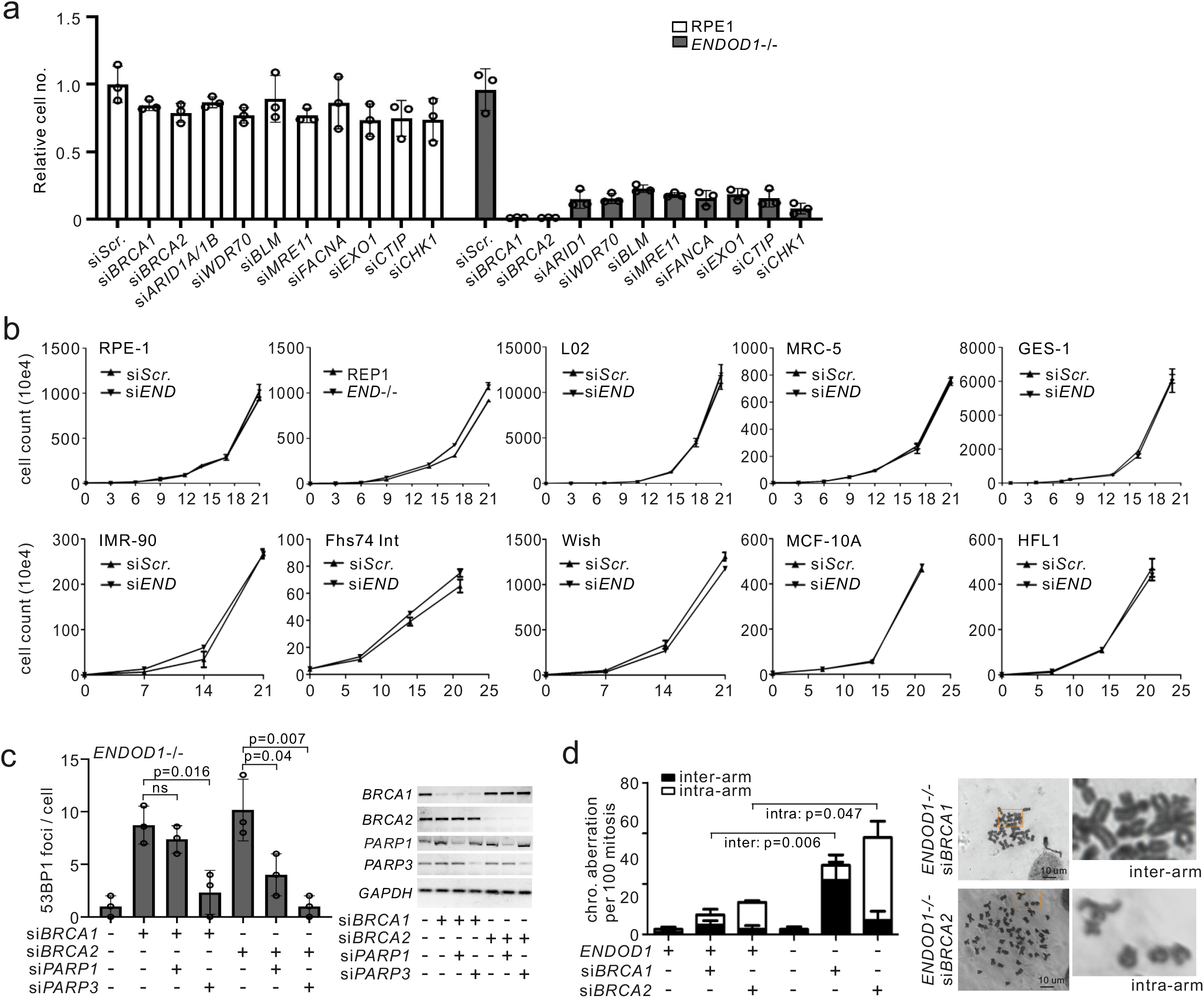
Depletion of *ENDOD1* causes SL with HRD. **a**. Relative viability of *ENDOD1*^-/-^ and control RPE1 cells subjected to the indicated siRNA treatments. siRNA was transfected three times with three-day intervals. Cell viability was determined by CCK8 colorimetric assay. n = 3 biologically independent experiments. **b**. Proliferation was quantified by haemocytometer cell counting over 3 weeks for the indicated immortalized non-cancerous cells with control siRNA (si*Scr*.) or si*ENDOD1* treatment every 3 days. Control experiments using RPE1 and *ENDOD1*^-/-^ are shown in the first two panels. **c**. Left: quantification of 53BP1 foci in *ENDOD1*^-/-^ cells 72 hours following treatment with the indicated siRNAs. n = 3 biologically independent experiments. Error bars: SEM. Significance test: two-tailed Student’s *t* test. ns: not significant. Right: semi-quantitative PCR showing knockdown efficiency. **d**. Ablation of *BRCA1* or *BRCA2* in *ENDOD1* null cells results in increased chromosome aberrations. Left: quantification of inter- and intra-chromosomal aberrations in RPE1 or *ENDOD1*^-/-^ cells treated with si*BRCA1* or si*BRCA2*. n = 3 biologically independent experiments. Error bars: SEM. Significance test: two-tailed Student’s *t* test. Right; representative images.

PARPi-induced HRD cell death is known to coincide with the generation of toxic DNA structures^13^. Consistent with this, *BRCA1* siRNA treatment of *ENDOD1*^-/-^ cells resulted in increased γH2AX and 53BP1 foci, common markers of DSBs (Figure 2c and Supplementary Figure 3b), and elevated chromosomal aberrations (Figure 2d), mimicking the consequences of PARPi treatment of HRD cells^13^. Similarly, depleting *BRCA2* in *ENDOD1*^-/-^ cells elevated 53BP1 foci formation (Figure 2c and Supplementary Figure 3c). ENDOD1 - HRD SL effects were dependent on the presence of PARPs (Figure 2c and Supplementary Figure 3c) but were not reversed by PARPi treatment (Supplementary Figure 3d), indicating that they are a result of “PARP trapping” and not PARP activity^13^. The cytotoxicity and DSB generation upon concomitant inhibition of HR and ENDOD1 could be reproduced in HRD breast cancer cell line MCF-7 (ref^14^) (Supplementary Figure 3e-f). Thus, we conclude that ablation of *ENDOD1* phenocopies PARPi in compromising the genomic integrity of HR-defective cells.

### ENDOD1 is SL with *TP53*

We next tested a panel of cell lines to establish the SL profile of si*ENDOD1* treatment in cancer cells (Figure 3a, Supplementary Figure 4 and Table S1). Unexpectedly, in addition to preventing proliferation of HRD cancer cells, si*ENDOD1* also inhibited the proliferation of multiple non-HRD cancer cells. Upon further analysis this SL correlated with *TP53* status (Figure 3b), a potentially important observation. The *TP53* mutations spanned the common mutation hotspots (i.e. R248, R273 and R280)^15^. Cell lines, including A549 and MDA-MB-361 that do not carry *TP53* or HR mutations were not sensitive to si*ENDOD1*. To rule out off-target effects we demonstrated that si*ENDOD1*-induced toxicity in the C33A cancer cells (*TP53-R273C*) could be reproduced with distinct siRNAs (Supplementary Figure 4b). We also validated the SL between p53 and ENDOD1 using our *ENDOD1*^-/-^ and RPE1 control cells. While siRNA control treated *ENDOD1*^-/-^ and si*TP53* treated RPE1 cells were viable, si*TP53* treated *ENDOD1*^-/-^ cells arrested in G1 within 60 hours and cell death became apparent at 5 days and was extensive after 7 days (Figure 3c). Concomitant treatment of RPE1 cells with si*ENDOD1* and si*TP53* also revealed SL (Supplementary Figure 4c). The cell death correlated with nuclear abnormalities and markers of apoptosis (Supplementary Figure 4d). Knockdown of *TP53* exacerbated the G1 cell cycle arrest that is already apparent in *ENDOD1*^-/-^ cells (Supplementary Figure 4e). Importantly, the toxicity of si*TP53* to *ENDOD1*^-/-^ cells can be rescued by ectopic expression of *ENDOD1* full length protein (Supplementary Figure 4f).

**Figure 3.**
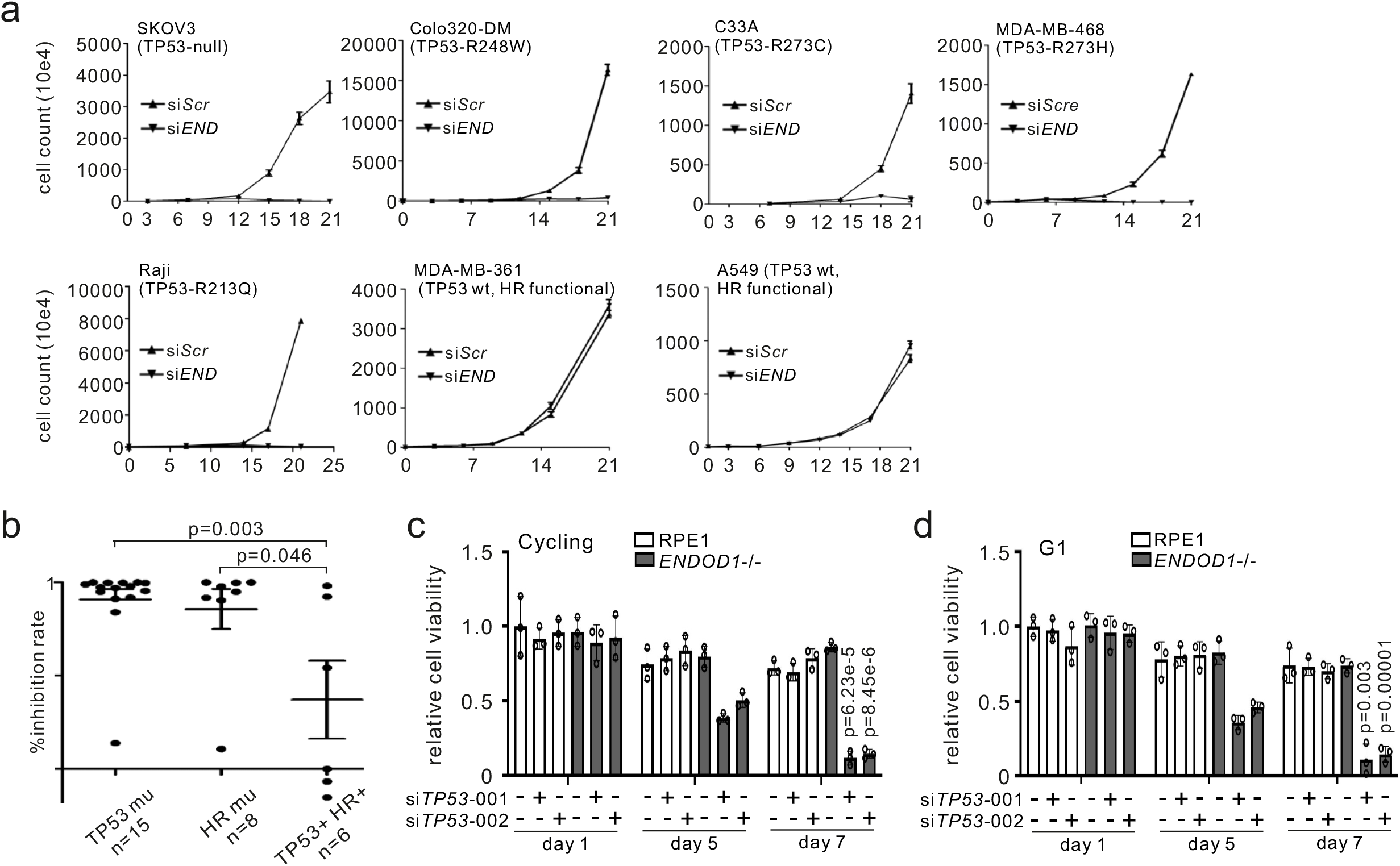
Concomitant loss of ENDOD1 and p53 causes SL. **a**. Proliferation curves for representative cancer cell lines following either si*ENDOD1* or control transfection. n = 3 biologically independent experiments. Error bars: SEM. At 3-day intervals cells were passaged, counted by haemocytometer and transfected. Reported *TP53* status is indicated. **b**. Correlation of *TP53*/HR mutations with cytotoxicity of si*ENDOD1* in terms of inhibition rate (%) for cancer cell lines (each point represents mean of n = 3 biologically independent experiments). Error bars; SEM. **c**. Relative viability determined by CCK8 colorimetry of *ENDOD1*^-/-^ and control RPE1 cells 5 and 7 days after treatment with two different si*TP53* (si001 and si002). n = 3 biologically independent experiments. Error bars: SEM **d**. Equivalent experiment as in **c**, using cells arrested in G1 by serum starvation. n = 3 biologically independent experiments. Error bars: SEM. All significance tests: two-tailed Student’s *t* test.

The killing effect of si*TP53* treatment of *ENDOD1*^-/-^ cells was also apparent in G1-arrested non-cycling cells (Figure 3d) that do not incorporate BrdU and lack bulk DNA synthesis (Supplementary Figure 4g), suggesting that the cytotoxicity is independent of DNA replication. The SL between *ENDOD1* and *TP53* was also recapitulated in coisogenic HCT116 cells (colon cancer): HCT116 cells expressing wildtype p53 remained viable, but those expressing an isoform of p53 (Δ40p53)^16^ did not proliferate (Supplementary Figure 4h). Taken together, these data indicate *ENDOD1* ablation, in addition to killing cells with HR defects due to chromatin-associated PARP, is synthetic lethal to cells with functional loss of *TP53* even in the absence of DNA replication.

### *ENDOD1 TP53* SL correlates with ssDNA formation

To characterise the SL interaction between ENDOD1 and p53 we used comet assays to asses DNA lesions occurring in G1 arrested *ENDOD1*^-/-^ and control RPE1 cells following si*TP53*. Alkaline comet assays revealed that si*TP53* induced a high level of DNA breaks in *ENDOD1*^-/-^ but not RPE1 cells (Figure 4a and Supplementary Figure 5a). Neutral comet assays did not show an increase in DNA damage in the same experiment. Thus, these breaks were likely in the form of SSBs. Consistent with this, analysis of phosphorylated RPA32 Serine 33 (pRPA32), a typical marker of single stranded DNA (ssDNA), showed that pRPA32 staining was elevated in the nuclei of either proliferating or G1 arrested *ENDOD1*^-/-^ si*TP53* cells, but was not elevated in the RPE1 cells treated with si*TP53* or *ENDOD1*^-/-^ cells co-transfected with control siRNA (Figure 4b and Supplementary Figure 5b). Non-denaturing α-BrdU staining showed evidence of ssDNA tracts in si*TP53* treated *ENDOD1*^-/-^ cells, but not control RPE1 cells (Figure 4c) and the α-BrdU signal was sensitive to S1 nuclease treatment (Figure 4d). The production of non-denatured α-BrdU staining upon si*TP53* treatment of *ENDOD1*^-/-^ cells was reproduced with a second si*TP53*, and could be eliminated by re-introducing *ENDOD1* upon lentiviral infection (Supplementary Figure 5c-d). Consistent with the production of ssDNA, S1 nuclease preferentially digested genomic DNA extracted from si*TP53* treated *ENDOD1*^-/-^ cells when compared to relevant controls (Figure 4e and Supplementary Figure 5e. In contrast to these indicators of ssDNA lesions, DSB surrogate markers, γH2AX and 53BP1 foci, were not significantly increased in G1-arrested *ENDOD1*^-/-^ cells 48-96 hours after si*TP53* transfection (Supplementary Figure 5f-g). We conclude that concomitant loss of ENDOD1 and p53 functions results in the generation of ssDNA and cell death.

**Figure 4.**
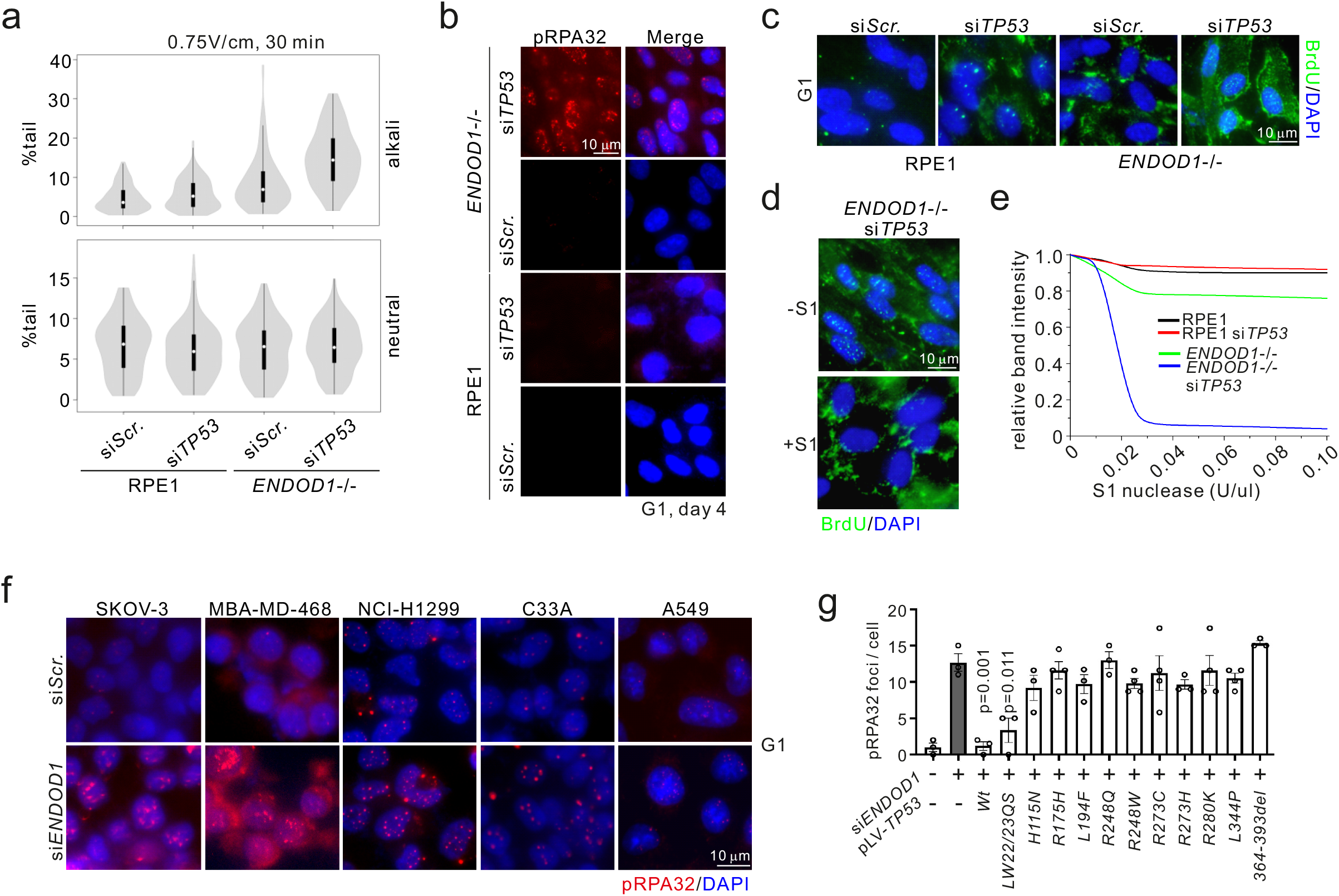
SL between *ENDOD1* and *TP53* correlates with ssDNA formation. **a**. Quantification of tail moments for neutral (bottom) and alkali (top) comet assays in G1-arrested *ENDOD1*^-/-^ and control RPE1 cells treated with si*TP53* or control siRNA. n = 150 cells from each of 3 biologically independent experiments. White dot: median. Thick whisker: third quartile. Thin whisker: upper/lower adjacent values (1.5x inter-quartile range). **b**. Immunofluorescent staining for pRPA32 in *ENDOD1*^-/-^ and control RPE1 G1 arrested cells 96 hours following si*TP53* or control treatment. Merged image is with DAPI staining. **c**. Equivalent experiment as in **b**, non-denatured staining for BrdU that was incorporated into cells before serum starvation. **d**. Non-denatured BrdU signals in *ENDOD1*^-/-^ si*TP53* cells with or without prior S1 nuclease digestion. Representative of 3 independent experiments. **e**. Quantification of agarose gel band intensity of undigested genomic DNA purified from the indicated cells after treatment with the specified units of S1 nuclease. Representative image of 3 independent experiments. A representative original gel for 0.02 U/μl is shown in Supplementary Figure 5e. **f**. pRPA32 staining in cancer cell lines that harbour *TP53* mutations 72 hours after treatment with control (si*Scr*.) or si*ENDOD1*. A549 is a *TP53* wild type control. **g**. Quantification of pRPA32 foci in si*ENDOD1* treated SKOV-3 cells complemented with the indicated *TP53* alleles. n = 3 - 4 biologically independent experiments. Error bars; SEM. Significance test: two-tailed Student’s *t* test.

### *TP53* hotspot mutations permit ssDNA production upon *ENDOD1* ablation

*TP53* mutations in human cancers exhibit diverse functional consequences including loss- and gain-of-function^17^ and almost invariably display abnormal DNA binding and transcriptional properties^18^. A recent study reported rapid PARP-dependent recruitment of p53 to sites of DNA damage that required the DNA binding and carboxyl terminal domains of p53^19^. We therefore assayed the production of pRPA32 foci upon si*ENDOD1* in a range of cell lines with defined *TP53* mutations. si*ENDOD1*-induced pRPA32 foci were increased in serum-starved (G1 phase) cancer lines harbouring a *TP53* null allele (SKOV-3 and NCI-H1299) in addition to cancer cells harbouring gain-of-function (GOF) mutants *R273C* (C33A) and *R273H* (MBA-MD-468). No increase in signal was apparent when the *TP53* competent line A549 was treated with si*ENDOD1* (Figure 4f).

To establish if this phenomenon in a broader range of *TP53* mutations we complemented a *TP53* null cell line, SKOV-3, with either wild-type or domain-specific and hotspot mutants and tested for pRPA32 foci upon si*ENDOD1* treatment (Figure 4g and Supplementary Figure 5 fecting the DNA binding domains (*R175H, L194F, R248Q, R248W, R273H, R273C, R280K*), the oligomerization domain (*L344P*), the putative nuclease activity (*H115N*)^20^ or the non-specific nucleic acid binding activity (C-terminal deletion 363-393)^21^, could not prevent ssDNA production. A transactivation domain mutation (LW22/23QS) showed an intermediate phenotype. Notably, the gain-of-function (GOF) mutants (i.e. *R273C, R273H* and *R280K*) in the DNA binding domain^22^ were, like the loss-of-function DNA binding domain mutants, not capable of preventing pRPA32 foci formation. This indicates that both decreased and increased p53 DNA binding causes cytotoxicity upon *ENDOD1* ablation. Collectively, these data show that *ENDOD1* inhibition is toxic to cells bearing *TP53* mutations that span the common hotspot sites.

### PARylation is required for ssDNA production

We next characterized the requirements for the generation of ssDNA. Dual treatment of *ENDOD1*^-/-^ cells with si*TP53* and either si*PARP1*/*2* or si*PARP3* significantly reduced pRPA32 foci formation compared to controls (Figure 5a). Ablation of either PARP1/2 or PARP3 function supressed pRPA32 foci when the *TP53* mutated cancer cell lines C33A, MBA-MD-468, SKOV-3 and NCI-H1299 were treated with si*ENDOD1* (Supplementary Figure 6a-b). We subsequently examined the chromatin association of PARP1 when p53 was depleted in *ENDOD1*^-/-^ cells. si*TP53* treatment of RPE1 control cells resulted in a modest increase in chromatin-associated PARP1. Untreated *ENDOD1*^-/-^ cells already displayed a modest level of chromatin associated PARP1. Treatment of *ENDOD1*^-/-^ cells with si*TP53* generated a pronounced increase in the chromatin association of both PARP1 and PARP3 (Figure 5b), suggesting that ssDNA formation and cell inviability are dependent on PARP association with chromatin.

**Figure 5.**
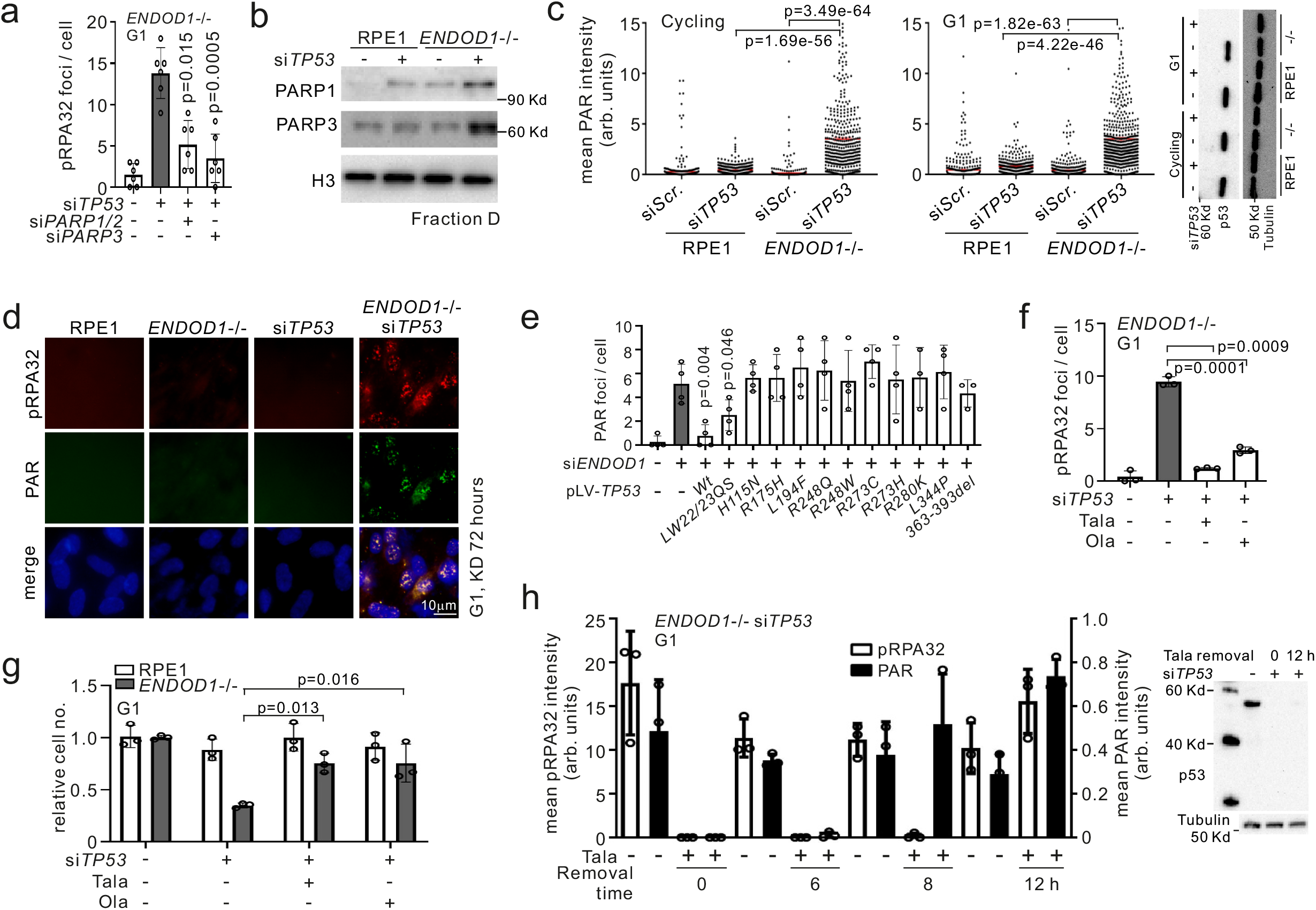
PARP activity is required for the SL between *ENDOD1* and *TP53*. **a**. pRPA32 foci in serum starved *ENDOD1*^-/-^ cells treated with the specified siRNAs. n = 6 biologically independent experiments. Error bars: SEM. **b**. Immunoblotting for PARP1 on chromatin (fraction D - see Figure 1e) in *ENDOD1*^-/-^ and control RPE1 cells following si*TP53* or siRNA control treatment. Representative image of 4 independent experiments. **c**. Quantification of nuclear PAR staining 72 hours following the indicated siRNA treatments of *ENDOD1*^-/-^ and control RPE1 cells (left: cycling, middle: G1 arrested, right: immunoblot showing knockdown efficiency for si*TP53*). arb. units: arbitrary units. n = 381-731 cells. Red bar: median. Whiskers: SEM. **d**. pRPA32 and PAR co-staining 72 hours following the indicated siRNA treatments of *ENDOD1*^-/-^ and control RPE1 cells. Representative image of 3 independent experiments. **e**. Quantification of PAR foci in SKOV-3 cells upon exogenous expression of the indicated *TP53* alleles. arb. units: arbitrary units. n = 3 – 4 biologically independent experiments. **f**. pRPA32 foci 3 days after serum starved *ENDOD1*^-/-^ cells were treated with si*TP53* and PARPi (10 nM Talazoparib, 100 nM Olaparib). n = 3 biologically independent experiments. Error bars: SEM. **g**. Relative proliferation 6 days after *ENDOD1*^-/-^ and control RPE1 cells were treated with si*TP53* and PARPi (2 nM Talazoparib, 20 nM Olaparib) n = 3 biologically independent experiments. Error bars: SEM. **h**. Talazoparib addition-removal assay in *ENDOD1*^-/-^ cells. 10 nM Talazoparib was added upon si*TP53* transfection and removed 72 hours later (time 0). Cells were fixed at the indicated time for α-pRPA32 or α-PAR staining. Immunoblot evaluation for the knockdown efficiency of *TP53* is shown on the right. arb. units: arbitrary units. n = 3 biologically independent experiments. Error bars; SEM. All significance tests: two-tailed Student’s *t* test.

Quantifying PAR staining after si*TP53* treatment of RPE1 cells showed that the ablation of p53 alone did not significantly affect PARylation, but si*TP53* treatment of *ENDOD1*^-/-^ cells resulted in a significant increase in nuclear PAR in both cycling and G1 arrested cells (Figure 5c). Co-staining for pRPA32 and PAR showed that pRPA32 signals overlapped with PAR (Figure 5d). This indicates that chromatin-associated PARP1 is active upon the concomitant loss of ENDOD1 and p53. Importantly, the specificity to ENDOD1 and p53 of these phenomena were validated in SKOV-3 (*TP53* null) cells: the emergence of pRPA32 and PAR foci upon si*ENDOD1* was significantly reduced following expression of siRNA-resistant *ENDOD1* (Supplementary Figure 6c) and, like pRPA32 foci formation (cf. Figure 4g), PAR foci were significantly reduced by the expression of wild type *TP53* (Figure 5e and Supplementary Figure 6d). Consequently, toxicity of si*ENDOD1* to SKOV-3 was reverted by *ENDOD1* or *TP53* expression (Supplementary Figure 6e). With the exception of *TP53*-*LW22/23QS* (which also significantly reduced pRPA32 foci formation), mutant alleles of *TP53* did not prevent PAR foci following si*ENDOD1* (Figure 5e). These data imply that p53 supresses PARylation and that, like the effect on pRPA32, this is independent of transactivation.

The DSBs induced by PARPi treatment of HRD cells are not dependent on PARP activity^13^. Consistent with this, as we showed above, PARPi treatment did not reduce the 53BP1 foci observed in *ENDOD1*^-/-^ si*BRCA1* cells (cf. Supplementary Figure 3d). Intriguingly, and in contrast to this, inhibition of PARP activity with either Talazoparib or Olaparib prevented the formation of pRPA32 foci in G1-arrested cells (Figure 5f) and significantly reduced cell killing upon si*TP53* treatment of *ENDOD1*^-/-^ cells (Figure 5g). The same is observed in multiple *TP53*-deficient cancer lines treated with si*ENDOD1* (Supplementary Figure 6f). When *ENDOD1*^-/-^ cells were treated with Talazoparib for the first 72 hours after si*TP53* treatment and then the Talazoparib was removed (time 0) the nuclear PAR signal became evident at 6 hours and was highly induced at 8 and 12 hours (Figure 5h). pRPA32 foci lagged behind, but reached the levels seen in cells not treated with Talazoparib by 12 hours. The Inhibition of parylation by either si*PARP1* or PARPi also strongly inhibited ssDNA formation, as evidenced by the reduction of the non-denatured BrdU signal (Supplementary Figure 6g). Interestingly, PARPi only attenuated, but did not eliminate, the S1-sensitivity of DNA isolated from si*TP53* treated *ENDOD1*^-/-^ cells (Supplementary Figure 6h). This may reflect that DNA lesions (i.e. SSBs) are present, but not processed into longer gaps, when cells are treated with PARPi. This suggests that long-tract ssDNA is only one form of DNA damage in si*TP53* treated *ENDOD1*^-/-^ cells. Thus, unlike the SL observed between PARPi - HRD and *ENDOD1* - HRD, the SL between *ENDOD1* - *TP53* requires PARP activity as well as its physical presence.

### XRCC1 is required for PARP activation and ssDNA formation

Using the same Talazoparib withdrawal protocol we next examined the requirement for single strand break repair in ssDNA generation in G1 arrested cells. Upon withdrawal of Talazoparib, concomitant knockdown of si*TP53* with *XRCC1*, the scaffold of the SSB machinery, eliminated pRPA32 foci in *ENDOD1*^-/-^ cells. si*TDP1* showed a modest reduction in foci and si*LIG3* did not have an impact (Figure 6a). Nucleotide excision repair (NER) factors (XPA and XPC) did not influence pRPA32/PAR (Supplementary Figure 7a). Consistent with this we observed that si*XRCC1* supressed the formation of PAR foci upon si*TP53* in *ENDOD1*^-/-^ cells (Figure 6b and Supplementary Figure 7b*ENDOD1*^-/-^ cells (Figure 6c). These results indicate that XRCC1 stimulates PAR catalysation by chromatin-associated PARP only when ENDOD1 and p53 are simultaneously absent. It also suggests that XRCC1-mediated break repair triggers ssDNA production.

**Figure 6.**
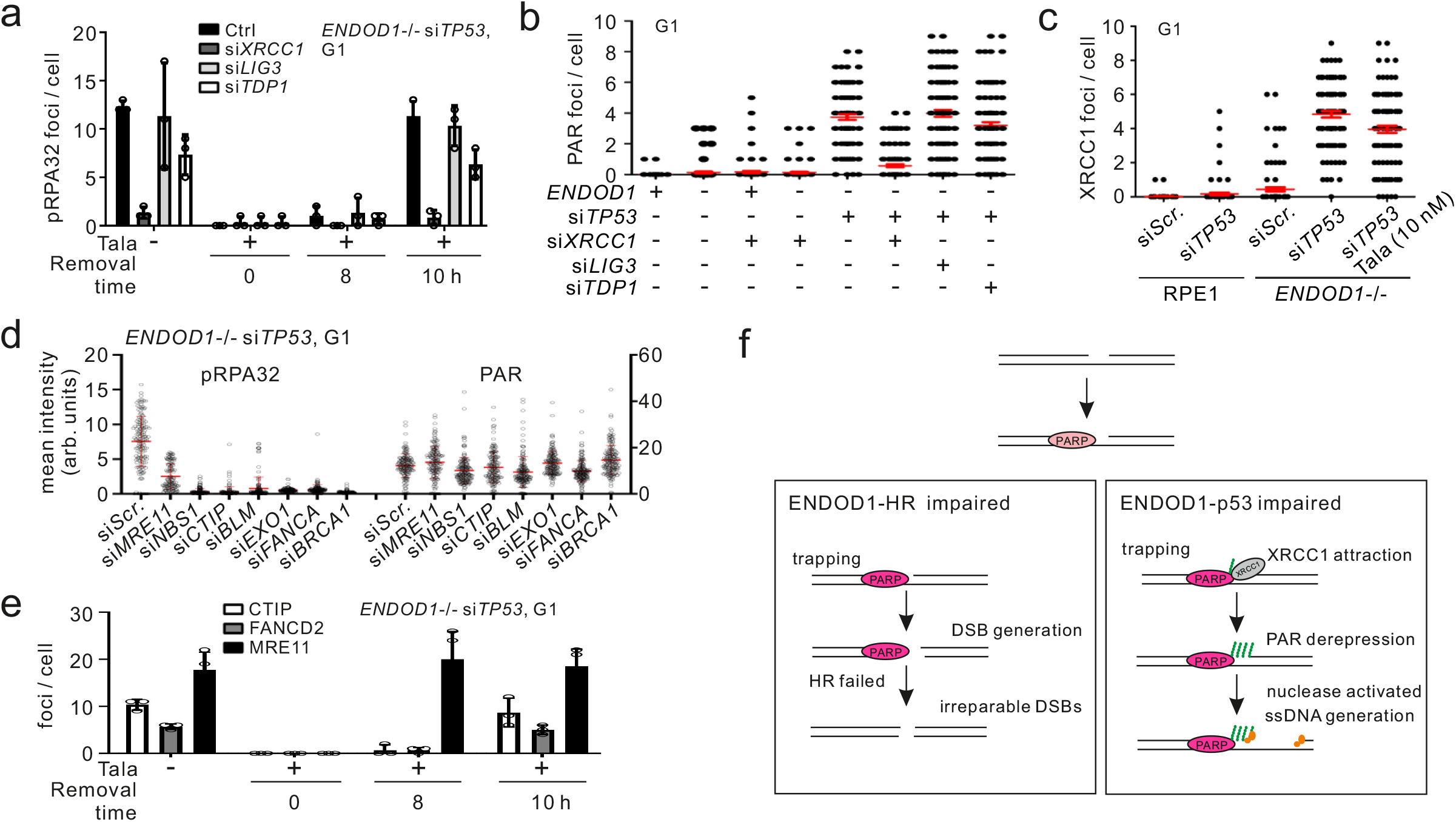
SSBR and HR machinery provides key signals for ssDNA production. **a**. Quantification of pRPA32 foci in a Talazoparib addition-removal assay with si*TP53* treated *ENDOD1*^-/-^ cells co-treated with the indicated SSBR siRNAs. 10 nM Talazoparib was added upon siRNA treatment and removed 72 hours later (time 0). n = 3 biologically independent samples. Error bars: SEM. **b**. Quantification of a-PAR foci in G1 serum starved *ENDOD1*^-/-^ and control RPE1 cells following treatment with the indicated siRNAs. n = (99-103) x 5 cells (each data point represents the average foci number of 5 cell counts). Red bar: median. Whiskers: SEM. **c**. Quantification of XRCC1 foci in serum-starved *ENDOD1*^-/-^ and control RPE1 cells following si*TP53* treatment with or without co-treatment with Talazoparib. n = (99-100) x 5 cells (each data point represents the average foci number of 5 cell counts). Red bar: median. Whiskers: SEM. **d**. pRPA32 and a-PAR foci in si*TP53* treated *ENDOD1*^-/-^ cells co-treated with and the indicated siRNAs targeting DNA end processing factors. n = 151-167 cells. **e**. Quantification of CTIP, FANCD2 and MRE11 foci in a Talazoparib addition-removal assay with si*TP53* treated *ENDOD1*^-/-^ cells. n = 3 biologically independent experiments. Error bars: SEM. 10 nM Talazoparib was added upon siRNA treatment and removed 72 hours later (time 0). **f**. Schematic model for how ENDOD1 protects genomic integrity together with HR (left) and p53 (right). See discussion for details.

### ssDNA production requires resection factors

To determine if the mechanism of ssDNA generation by PARP chromatin association in G1 phase involves the canonical DNA end resection enzymes that usually process DSBs in S/G2 phase of the cell cycle, we examined key resection factors in G1-arrested si*TP53* treated *ENDOD1*^-/-^ cells subjected to PARPi treatment and withdrawal. Knockdown of a range of resection factors, including MRE11, NBS, CTIP, BLM, EXO1, BRCA1 and FANCA suppressed pRPA32 foci formation, but not PAR formation, when G1 arrested *ENDOD1*^-/-^ cells were treated with si*TP53* (Figure 6d). Analysis for foci formation of resection factors identified an accumulation of MRE11, CTIP, BLM, FANCD2, NBS1 and BRCA1 foci (Supplementary Figure 7c). Like the formation of PAR foci (cf. Figure 5h), MRE11 foci formation in si*TP53* treated *ENDOD1*^-/-^ cells following Talazoparib removal was rapid, with foci visible at 8 hours (Figure 6e), whereas the formation of CTIP and FANCD2 foci only became apparent at 10 hours. This suggests that MRE11 as a key nuclease for generating the ssDNA. Interestingly, unlike α-NBS1 and α-BRCA1 antibodies, α-phospho-NBS1 and α-phospho-BRCA1 did not reveal foci (Supplementary Figure 7c), despite being robust markers of resected DSB ends in S/G2^23,24^. These data suggest that inappropriate activity of resection factors that would usually process DSBs in S/G2 phase are producing ssDNA tracts, even in G1 cells, when both ENDOD1 and p53 functions are impaired (Figure 6f). However, ssDNA production in the absence of p53 and ENDOD1 is mechanistically distinct from canonical DSB resection.

The impact of XRCC1 is dominant over the resection machinery as ablation of *XRCC1* significantly reduced levels of MRE11 staining (Supplementary Figure 7d), indicating that XRCC1 acts upstream of the resection machinery. Combining the results in Figure 6c, we propose that the ssDNA production in *ENDOD1* - *TP53* double mutant occurs in a stepwise manner: SSB-bound XRCC1 activates PARP1 that subsequently attracts the resection machinery to initiate long-tract ssDNA processing. Inhibition of PARP1 suppresses productive ssDNA generation but leaves SSBs unrepaired (cf. Supplementary Figure 6h).

### ENDOD1 is a potential drug target

*ENDOD1* is SL both with HRD and with *TP53* mutation, suggesting a wide range of potential target tumours. A key issue with drugs that target specific proteins is systemic tolerance. To address this, C57/B6 mice were injected twice weekly with siRNA for the murine homolog of *ENDOD1* (sim*Endod1*) or a disease causing control, sim*Wdr70*. Efficacy of whole animal gene silencing was assessed by semi-quantitative PCR to be ∼75% (Supplementary Figure 8a). After 60 days, sim*Wdr70* treated mice lost weight and were euthanised. 90 days into sim*Endod1* treatment littermates retained normal weight (Figure 7a). Histological examination revealed minimal pathological alterations in sim*Endod1* treated animals (Figure 7b), whereas sim*Wdr70* treated littermates displayed increased cell debris, fibrosis and disorganized tissue structures in the lung, intestine and liver. Carditis and event heart failure can also complicate cancer treatment with high-dose chemotherapeutic regimens^25^. Hematoxylin and eosin staining of heart tissues from sim*Endod1* treated animal showed no myopericarditis that manifested with lymphocyte infiltration (Figure 7b) and echocardiography showed normal cardiac function without hypertrophy in sim*Endod1* treated animals, with no changes to anatomic dimension or systolic and diastolic function (Supplementary Figure 8b).

**Figure 7.**
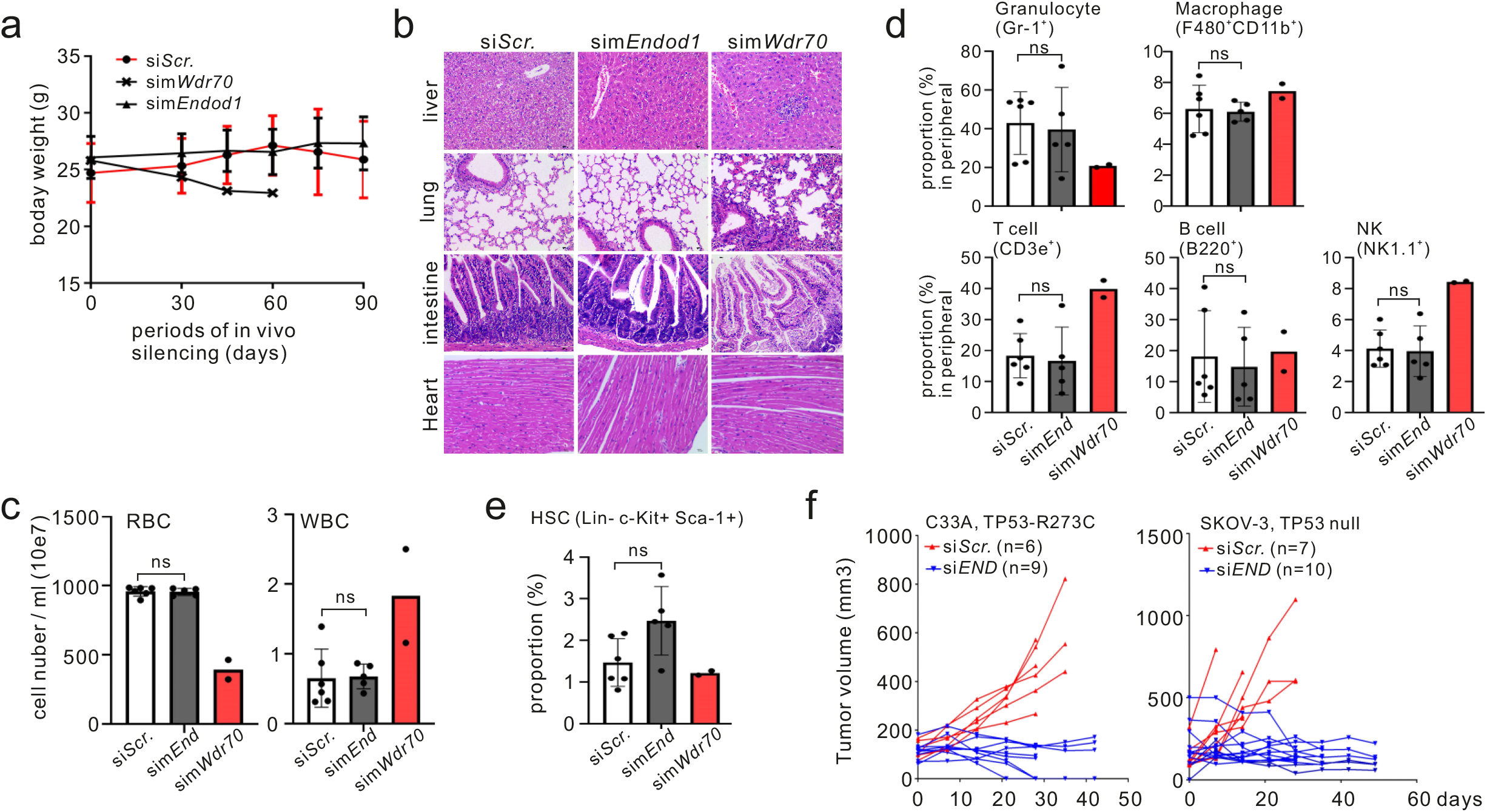
m*Endod1* systemic knockdown is well tolerated. **a**. Body weight tracked through 90 days for the indicated *in vivo* knockdown groups (two injections per week). sim*Wdr70* mice were sacrificed at 60 days due to severe disease. **b**. Representative images of hematoxylin and eosin staining of paraffin-embedded sections from the indicated tissues. **c**. Haemocytometer counts for peripheral blood cells at the endpoint of the experiment for each knockdown group. Control 90 days (n = 6 animals), sim*Endod1* 90 days (n = 5 animals), sim*Wdr70* 60 days (n = 2 animals). Significance test: two-tailed Student’s *t* test. ns: not significant. **d**. FACS analysis for peripheral myeloid and lymphoid cells when experiments terminated. n = 6, 5 and 2 animals for simScr, simEnd and simWdr70 respectively. Error bars: SEM. Significance test: two-tailed Student’s *t* test. ns: not significant. Cell surface markers used are shown in parentheses. **e**. Equivalent FACS analysis as above for bone marrow HSC. **f**. Anti-tumour treatment using whole animal *in vivo* knockdown of *ENDOD1* for p53-deficient (SKOV-3 and C33A) xenograft models in nude mice. Volumes of individual tumours were measured. *x*-axis: treatment days. Numbers of animals (n) are indicated.

A common side effect of antineoplastic drugs is myelosuppression and cytopenia. In the peripheral blood of sim*Endod1* mice, no apparent cytopenia was observed (Figure 7c) and populations of T cells (CD3e^+^), NK (NK1.1^+^) cells, granulocyte (Gr-1^+^) and macrophage (F480^+^CD11b^+^) were conserved (Figure 7d). The lack of peripheral myelosuppression in sim*Endod1* is supported by the preservation of hematopoietic stem cells (HSC) in bone marrow (Lineage^-^ Sca1^+^ c-Kit^+^ cells: LSK) (Figure 7e). Consistent with this, the multilineage potential of bone marrow was generally sustained for T/NK cells and myeloid compartments (granulocyte and macrophage) (Supplementary Figure 8c). We conclude that the suppression of m*Endod1* function does not cause short-term myelotoxicity (<3 months).

To establish the therapeutic potential of ENDOD1 xenograft tumour models were established from the *TP53* mutated SKOV-3 and C33A cancer cells and a *TP53* wildtype control cancer line, MDB-MA-361. Progression of SKOV-3 and C33A tumours was effectively curbed upon *in vivo* si*ENDOD1* treatment when compared to the si*Scr*. control group (Figure 7f). In contrast, si*ENDOD1* was ineffective in restraining disease progression of MDB-MA-361 Tumours (Supplementary Figure 8d). Taken together, these data suggest that exploiting *TP53* -*ENDOD1* SL may yield therapeutic advantage, while minimizing unwanted side-effects seen in conventional chemotherapy.

## Discussion

Previous reports identified ENDOD1 as an RNF26-interacting protein that modulates cGAS-STING-dependent innate immune signalling through an as yet uncharacterised mechanism^9^. Our identification of ENDOD1 during experiments that initially aimed to identify PARPi-dependent responses in a specific HRD cell line (HBx-induced CRL4^WDR70^ defect) led us to characterise ENDOD1 in the context of DNA repair. We show that ENDOD1 loss results in increased PARP chromatin association and, as with PARPi treatment, this manifests in SL with HRD. This SL resembled that of PARPi treated HRD cells, involving the accumulation of lethal DNA structures during replication that require resolution by HR-dependent pathways, and relying on the physical presence of PARPs rather than their activity (Figure 6f, left). This suggests that targeting ENDOD1 in HRD cancers may provide an alternative to PARPi therapy.

Surprisingly, we identified a second distinct SL interaction with mut-p53 that encompasses the major cancer-specific hotspots. The incompatibility of *ENDOD1* ablation and *TP53* mutations was also linked to PARPs, but followed a distinct pattern: SL was evident in G1-arrested serum starved cells, correlated with the formation of tracts of ssDNA and was dependent on both the physical presence of PARPs and their activity (Figure 6f, right). Mechanistically we found that, in the absence of both ENDOD1 and p53, intrinsic PARP1 chromatin association was elevated and that XRCC1 was required to stimulate the catalytic activity of chromatin-associated PARPs in order to generate ssDNA and cell death. ssDNA generation was also dependent on the activity of the resection machinery that is usually inactive in G1. These observations define a new PARP-dependent DNA damaging activity that is distinct from those engendered by PARPi. What structures are initiating PARP recruitment and activation, how this is prevented by ENDOD1 and p53, plus how the resection machinery is activated inappropriately remain important future questions.

*TP53* mutation occurs in the majority of human tumours and is associated with therapy-refractory malignancy. Previous efforts to exploit p53 in cancer treatment have largely focused on restoring the pro-apoptotic and cell cycle arrest potential of mutant versions^26^. Our work opens up the possibility that a wide range of *TP53* mutations, including common gain-of-function alleles, could be exploited using a SL approach to induce toxic DNA lesions specifically in cancer cells, but not surrounding tissue. To establish if such an approach is feasible, we treated mice with sim*Endod1* to ascertain if the *in vivo* knockdown of m*Endod1* could be tolerated. In contrast to whole animal ablation of m*Wdr70*, which acted as a disease-causing control, whole animal ablation of m*Endod1* was well tolerated, with minimal evidence of relevant tissue damage and acceptable levels of myelosuppression. We then established three xenograft models and demonstrated that whole animal treatment with human si*ENDOD1* resulted in profound disease control for two *TP53* mutated cancers when compared to mock treated animals. The third xenograft model of a *TP53* wildtype cancer did not respond to si*ENDOD1*. Our work opens up the possibility that a wide range of p53 mutations could be exploited using a SL approach to induce toxic DNA lesions specifically in cancer cells, but not in surrounding tissue. In summary, we identify ENDOD1 as a potential wide-spectrum and cancer-specific target for SL drug discovery.

## Methods

Information for cell lines, siRNA, primers and antibodies used in this study are listed in Table S1-4.

### Cell culture

The *ENDOD1*^-/-^ clone was obtained by targeting exon 1 with gRNA (5’-CAGCCTCTTCGCCCTGGCTGG-3’) in RPE1 cells (Supplementary Figure 9c). The resulting insertion (C→CG) creates a frameshift in the ORF at amino acid position Arg4. All human cell lines were cultured in complete media supplemented with 10 or 20% foetal bovine serum (FBS) according to ATCC protocols. Where indicated, G1 arrest was induced by serum starvation for 8-11 days in complete media: RPE1 and *ENDOD1*^-/-^, 1% FBS; GES-1, SKOV-3, NCI-H1299, MBA-MD-468 and C33A, 0.5% FBS. Representative FACS assays for cycling and serum starved cells are shown in Supplementary Figure 9a. All cell lines tested negative for mycoplasma contamination and were authenticated by providers (National Collection of Authenticated Cell Cultures, Shanghai and iCell Bioscience Inc, Shanghai). Double stranded siRNA’s were obtained from Ribobio, Guangzhou, China. Plasmid and siRNA transfections were performed using Lipofectamine 3000 (Invitrogen) or FuGENEHD transfection Reagent (Roche, E231A). 1 ug plasmid or 50 nM siRNA were applied per 10^6^ cells unless otherwise stated. Efficiencies of gene silencing for frequently used cell lines in this study are presented in Supplementary Figure 10a-d. Targeting sequence of siRNA are listed in Supplementary Table S2.

### Plasmids

For cloning of Flag-tagged *ENDOD1* and *TP53*, PCR fragments were inserted into the *Eco*RI site of pLVX-Flag -IRES-ZsGreen1 plasmid, using the In-Fusion cloning kit (Clontech, 639650). Point mutants for *ENDOD1* and *TP53* were converted from parental pLVX-Flag -IRES-ZsGreen1 plasmids using QuikChange Lightning Site-Directed Mutagenesis Kits (Stratagene, 200519). Truncations of *ENDOD1* and mutations of *TP53* were obtained by fusing different PCR fragments using In-Fusion cloning kit (Clontech, 639650). Primers used in this study were listed in Supplementary Table S3.

### Chemicals and genotoxic treatments

To induce DNA damage, cells were treated with the indicated concentrations of: CPT (Selleck, S2423); HU (Selleck, S1896); Cisplatin (Supertrack Bio-pharmaceutical, 131102); CX5461 (Selleck, S2684) and H_2_O_2_ for the times stated for each experiment. Suppression of PARP enzymatic activities was achieved by adding pre-determined concentrations of Olaparib (Selleck, S1060) or Talaparib (Selleck, S7048) as indicated for individual experiments.

### LC-MS/MS for PARPi induced proteomic changes

HBV-integrated T43 cells were continuously treated with 100 nM Olaparib for 2, 4 and 6 days. Whole cell samples were ground in liquid nitrogen and lysed in 8 M urea supplemented with 1% protease inhibitor cocktail (Calbiochem). Lysates were centrifuged at 12,000 x g and the supernatant was quantified by BCA assays (Beyotime). For mass spectrometry: in brief, protein solutions were reduced with dithiothreitol (5 mM) and alkylated with iodoace-tamide (11 mM). Samples were diluted by Triethyl ammonium bicarbonate to reduce the final concentration of urea to less than 2M. After two rounds of trypsin (Promega) digestion, peptides were desalted using Strata X C18 SPE columns (Phenomenex) and reconstituted in 0.5 M TEAB. Peptide solutions were dissolved in UHPLC buffer A (0.1% (v/v) formic acid in water) and loaded for LC separation on a NanoElute high-performance liquid chromatography (UHPLC) system (Bruker Daltonics), using a 90 min LC gradient at 300 nL/min. MS data were collected using a tims-TOF Pro mass spectrometer (Bruker Daltonics), and processed using the Maxquant search engine (v.1.6.6.0). Mass spectra data were searched against SwissProt Human database concatenated with reverse decoy database.

### Cell proliferation and viability

Cells treated with siRNA or chemicals were incubated in complete media. Proliferation curves were determined by counting cell number by haemocytometer every 3-5 days upon passage, dependent of the growth rate of individual cell lines. Inhibition rate (%) for each cell line was calculated as (1-siRNA/Control) x 100%. siRNA and control represent the number of remaining cells for the specific siRNA and si*Scramble* (si*Scr*.) control at the endpoint of the experiment. Relative cell survival: the ratio calculated by dividing the number of cells in the treatment group with untreated control at the endpoint of the experiment. For Giemsa staining, remaining cells were fixed with prechilled methanol for 10 min and stained with Giemsa solution (Baso, BA-4122). Cells subjected to CCK8 viability assay were replaced with fresh medium containing 10% CCK8 and incubated for 2-4 h. The colorigenic supernatant was carefully aspirated and transferred to new 96-well plates. Absorbance values at 450 nm was detected by Multifunctional enzyme marker. Relative cell survival was calculated according to OD values. A medium blank control was set for each plate and three replicates were included for statistical analysis.

### Measurement for PARP-DNA complex

The method for assessing tight DNA/chromatin association of PARPs is described elsewhere^12^. Briefly, cells were treated for the indicated time with or without drugs and trypsinizied. Pellets were then extracted with different stringency. 3×10^6^ cells were treated with 100 μl of hypotonic buffer (100 mM MES-NaOH, pH 6.4, 1 mM EDTA and 0.5 mM MgCl_2,_ 0.05% TritonX-100) supplemented with protease inhibitors (Complete Mini, Roche), layered gently onto 100 μl of hypotonic buffer containing 30% sucrose and centrifuged at 15,000 g at 4°C for 10 min. The P1 fraction was obtained by dissolving pellets in 100 μl of buffer A (50 mM HEPES-NaOH, pH 7.5, 100 mM KCl, 2.5 mM MgCl_2_, 0.05% TritonX-100 and protease inhibitors). 50 μl of P1 was centrifuged at 15,000 g at 4°C for 10 min. The supernatant was preserved (fraction A) and pellets dissolved in 50 μl buffer B (50 mM HEPES-NaOH, pH 7.5, 250 mM KCl, 2.5 mM MgCl_2_, 0.05% TritonX-100 and protease inhibitors), centrifuged at 15,000 g at 4°C for 10 min. The supernatant was preserved (fraction B) and the pellet redissolved in 50 μl buffer C (50 mM HEPES-NaOH, ph 7.5, 500 mM KCl, 2.5 mM MgCl_2_, 0.1% TritonX-100 and protease inhibitors), centrifuged at 15,000 g at 4°C for 10 min. The supernatant was reserved (fraction C). The pellet was dissolved and digested in 50 ml buffer A with 2 mM CaCl_2_ and 4,000 units of micrococcal nuclease (M0247S, NEB) at RT for 20 min. The supernatant was collected (fraction D) after centrifugation at 15,000 g at 4°C for 10 min.

### Immunofluorescent staining

Briefly, cells were grown on coverslips and fixed with Carnoy’s fluid (methanol:glacial acid:3:1) or 4% paraformaldehyde (PFA) and permeabilize with 0.3% TritonX-100 followed by blocking in PBS with 3% BSA, 3% donkey serum and 0.2% Triton X-100. Primary antibodies were diluted with antibody buffer (0.1% Triton 5% BSA in PBS) and incubated for 2 hours at ambient temperature. Primary antibodies were detected with anti-rabbit-Cy3 or anti-mouse-FITC. Fluorescent images were acquired using an Olympas (BX51) or LEICA DM4 B and images were processed analysed using Image-Pro Plus software. ENDOD1 Antibody used for immunofluoscent staining experiment is ABclonal (A16502). In general data from 200 - 500 cells were quantified per sample in each independent experiment for statistical analysis of imaging assays. For non-denatured BrdU staining of ssDNA, proliferative cells on coverslips were pre-labelled with 40 ug/ml BrdU for 48 hours before serum starvation, followed by fixation with methanol-acetic acid buffer (3:1) for 15 min. Coverslips were sequentially incubated in blocking buffer (0.3% Triton, 5% donkey serum in PBS) for 15 min and BrdU antibody (1:100) added and incubated overnight at 4°C. Microscopic visualization and image capture were performed as described above.

TUNEL Cell death assays were performed following the instructions of DeadEnd™ Fluorometric TUNEL System kit (Promega, G3250). Briefly, cells were fixed in freshly prepared 4% methanol-free formaldehyde PBS solution for 25 min at 4°C, followed by washing with PBS and 0.3% Triton X -100/PBS. After equilibration, cells were reacted with rTdT solution at 37°C for 60 minutes and the reaction was terminated by adding 2 x SSC for 15 min at room temperature. Samples were re-stained with propidium iodide (1 μg/ml in PBS) in the dark and fluorescent images captured.

### Flow cytometry and Fluorescent-Activated Cell Sorting (FACS)

Cell cycle analysis was performed using 3 × 10^4^ trypsin-dissociated cells. After rinsing in PBS two times cells were fixated with 75% ethanol overnight and stained with PBS-PI (50 μg/ml) for 20 min before cytometry using a BD FACSCalibur. Modfit software was used for data processing. For immunotyping of peripheral blood cells, erythrocytes were removed by treatment with chilled RBC buffer (15 mM NH_4_Cl, 1 mM KHCO_3_, 0.1 mM EDTA, pH 7.1-7.4) for 10 min to cause lysis. The product was centrifuged for 5 min at 400 x g and cells (1-2 × 10^6^ cells/sample) were resuspended in 100 μl PBS, followed by incubation with the appropriate dilution of fluorescent antibody conjugates (listed in Supplementary Table S4) for 30 min at room temperature or 45 min on ice. For bone marrow analysis, a single cell suspension was obtained by flushing bone marrow cells with PBS containing 2% FBS, followed by incubating with marker antibody as above. Cells were considered as live cells after FSC/SSC gating and then used in fluorescence histograms. Labeled cells were analyzed on Beckman Cytoflex S and Flow Jo V10 software were used for data analysis. Sorted cells were defined as the following: T cells (CD3e+), NK cells (NK1.1+), granulocyte (Gr-1+), macrophage (F480+ and CD11b+), hematopoietic stem cells (Lin-Sca1+ and c-Kit+). The border between negative and positive was determined by an isotype-matched control antibody. Gating strategy is exemplified in Supplementary Figure 9b.

### Preparation of metaphase chromosomal spread

Cells were plated in a 60-mm dish and arrested in mitosis by 2-hour treatment with colcemid (final concentration; 200 ng/ml). Cells were trysinized and pre-warmed 0.075 M KCl added and incubated for 20 min at 37°C. Four drops of freshly prepared fixative (3:1 solution of methanol:acetic acid) was added and cells were pelleted, resuspended in 5 ml fixative and incubated for 20 min at 4°C. After repeating the fixation two times, pellets were resuspended in 0.5 ml fixative solution. Two or three drops of cell suspension were precipitated onto a pre-chilled microscope slide from a height of 18 inches. Slides were thoroughly air-dried and stained using Giemsa. The mitotic chromosomes were observed and evaluated using an Olympus fluorescence microscope (BX51) at 1000X magnification.

### Comet assay

Cells were treated with 10 mM H_2_O_2_ on ice and subsequently transferred to drug-free complete media for the indicated recovery periods. Cells were then trypsinizied and resuspended in PBS and mixed with an equal volume of 1.6% low-gelling-temperature agarose (Thermo, 16520100) maintained at 42°C. The mixture was immediately spread onto a frosted glass slide (Fisher) pre-coated with 0.8% agarose (Thermo, 16500500) and air dried over-night at RT to set. For alkaline electrophoresis, slides were immersed in pre-chilled lysis buffer (2.5 M NaCl, 10 mM Tris-HCl (pH10), 100 mM EDTA, 1% Triton X-100, 1% DMSO) for 1 h, washed twice with pre-chilled distilled water for 10 min and placed for 20 min in pre-chilled alkaline electrophoresis buffer (300 mM NaOH, 1 mM EDTA). Electrophoresis was then conducted at 0.75 V/cm for 15 min (0.75 V/cm for 30 min for detecting endogenous DNA breaks). Slides were and subsequently neutralized 3 times in 400 mM Tris-HCl pH7.0 for 5 min. For the neutral Comet Assay, electrophoresis solution (300 mM sodium acetate, 100 mM Tris-HCl pH 8.3) was used. DNA was fixed with absolute ethanol for 20min then stained with EtBr (2 ug/ml) for 5 min and washed twice in ddH_2_O. The percentage of tail moment was calculated by dividing the intensity of tails by that of heads as measured with Image Pro Plus. Data presented in violin plots are 150 cells per data point from 3 independent biological repeats. The three repeats showed the same trend.

### S1 nuclease digestion

Genomic DNA was extracted from 10^6^ cells using High Pure PCR Template Preparation Kit (Roche, 11796828001). 150 ng of purified DNA was digested in Reaction Buffer (200 mM sodium acetate [pH 4.5], 1.5 M NaCl and 10 mM ZnSO4) with different dilutions of S1 Nuclease for 10 mins at room temperature and inactivated by heating at 70°C for 5 min in the presence of EDTA. Digested DNA was resolved by 0.8% agarose electrophoresis. Images were captured by Chemidoc XRS(Bio-Rad) and bands quantified by ImageJ.

### Animal, biopsies and histochemical staining

The animal experiments in this study were registered and approved by the Medical Ethical Committee of the West China Second University Hospital of Sichuan University on 8th June, 2018 (approval Reference number, Medical Research 2018 (015). Experiments were carried out in accordance with the approved guidelines. All mice were housed in standard SPF condition throughout the experiments at a maximum of 5 per cage with 12 hour light / dark cycles at 23°C and 55% humidity. C57BL/6JGpt mice were purchased from Gempharmatech co., Ltd. Sexually mature mice (C57/B6) were randomly divided into control siRNA treatment, sim*Endod1* and sim*Wdr70*. In vivo knockdown was performed by injecting specific double strand siRNA (10 mg/kg) and 100 μl RNA transfection reagent (Biotool, B45215). Injection started from 10 week after birth and occurred twice per weekly via alternative tail vein or intraperitoneal route. Body weight and the health condition of each mouse was examined weekly. Dissected tissues were embedded in paraffin and sliced at 8 microns for histochemical staining and images were captured by BX51 (Olympus). 5 slices of biopsied tissues were examined for HE staining to avoid individual variation. Echocardiography was performed on a VisualSonics Vevo 2100 and output analysed with Vevostrain software, and operated by a person blinded to the treatment group. Animals were handled awake and held in a standard handgrip.

### Xenograft model in nude mice

Female athymic nude immunodeficient mice (BALB/cGpt-Foxn1nu/Gpt, purchased from Gempharmatech co., Ltd) of 4-5 weeks of age were used for xenograft implants. Mice were subcutaneously inoculated in both sides of armpits or hind flanks (2-4×10^6^ cells *per* site). Animals were randomized using random number table into control and treatment groups when xenografts had reached an average volume of approximate 100 mm^3^. For treatment, animal was administered with 10 mg/kg *ENDOD1* siRNA twice a week with 100 μl RNA transfection reagent. A parallel group of mice was administered with control siRNA. Inoculated and siRNA-administered mice were observed each day. Tumour Size was measured by Vernier calliper and tumor volume calculated by the three-dimensional measurement (length×width×width/2) until termination of the experiments. Experiments were performed blind.

### Statistics

All histograms are presented as means ± SEM. For quantitative analysis including immunoblotting, image analysis and repair analysis at least three independent biological repeats were carried out. Analysis for significant difference between two groups was performed either by Two-tailed Student’s *t* tests (2 sided) using GraphPad Prism 6 or SPSS 16.0. or by non-parametric Kruskal tests using Python scipy.stats package. End-point values of cell survival assays were used for statistical analysis. The level of significance was set as p<0.05.

## Data Avaliability

The data that support the findings of this study are available from the corresponding authors upon reasonable request.

## Acknowledgements

This work is supported by NSF China (31771580, 82073027, 81702795, 81630038 and 81971433), Sichuan University (2020SCUNL204, SCU2019C4198), National Key Research and Development Program of China (2019YFE0198400, 2018YFC1002801), Department of Science and Technology of Sichuan Province (2019JDTD0020, 2020YFS0131, 2020YFH0019), and Wellcome Trust Award 110047/Z/15/Z. We thank Owen Wells, Penny Jeggo, Keith Caldecott, Dacai Liu and Jichen Hu for critical comments. We also thank PTM Biolab (Hangzhou) for mass spectrometry analysis.

## Author contribution

C.L., A.M.C and Z.T. co-discovered a repair role for ENDOD1. C.L. and Z.T. established SL between ENDOD1, TP53 and HR defects, and formulated the SL model with A.M.C.. M.Z. and X.W. performed biochemical and cell biology assays. P.Y. analysed heart function. C.G. and X.Z. assisted immunoblotting and cytometry. J.C., D.M., H.L. and D.K advised on p53, animal experiments and DNA repair. C.L. and A.M.C. supervised the overall project and wrote the manuscript.

