## Supplementary data for "Synthetic lethality between TP53 and ENDOD1"

**Supplementary Figures**

**Supplementary Figure 1.**

**a**. Sequences of ENDOD1 peptides captured by LC-MS/MS from T43 cells (expressing HBx from HBV) following 6 days treatment with Olaparib. **b.** Multiple isoforms of ENDOD1 are detected by immunoblotting with Abcam $\alpha$-ENDOD1 in RPE1 cells. Left panel: Control siRNA (si*Scr.*) and si*ENDOD1* treated RPE1 cells. The main full-length product is estimated to have a MW of 55 Kd and is labelled FL/a. Bands b-d appear to be truncated species. Asterisks: non-specific bands. Right panel: immunoblotting with Abcam $\alpha$-ENDOD1 of RPE1 control cells, CRISPR/Cas9 edited RPE1 *ENDOD1*$^{-/-}$ cells and RPE1 cells expressing a full-length C-terminally flag tagged construct (FL-Flag). Representative image of 3 biological repeats. **c**. Left panel: immunoblotting with Abclone $\alpha$-ENDOD1 of exogenously expressed Flag-tagged ENDOD1 in RPE1 cells. Right panel: immunoblotting with $\alpha$-Flag. The full-length construct Flag-tagged at the N-terminus (Flag-FL) is not detected by the $\alpha$-Flag antibody, whereas the full-length construct Flag-tagged at the C-terminus (FL-Flag) is recognised, as are N-terminally tagged constructs missing the first 21 amino acids. All constructs were recognized by $\alpha$-ENDOD1, indicating cleavage of the predicted signal peptide. Representative image of 3 biological repeats. **d**. Survival analysis of the indicated cells subjected to the stated genotoxic treatments. *ENDOD1*$^{-/-}$ cells showed modest sensitivity to CPT, Cisplatin and the G4 stabilising agent Cx5461, but not to HU or IR. Unexpectedly, *ENDOD1*$^{-/-}$ cells showed increased resistance to $H_2O_2$, even when treated during serum starvation (G1 arrest). The increased resistance to $H_2O_2$ was recapitulated in GES-1 cells (normal gastric cell line) upon si*ENDOD1* treatment when compared to control (si*Scr.*). n = 3 biologically independent samples. Error bars: standard error of the mean.

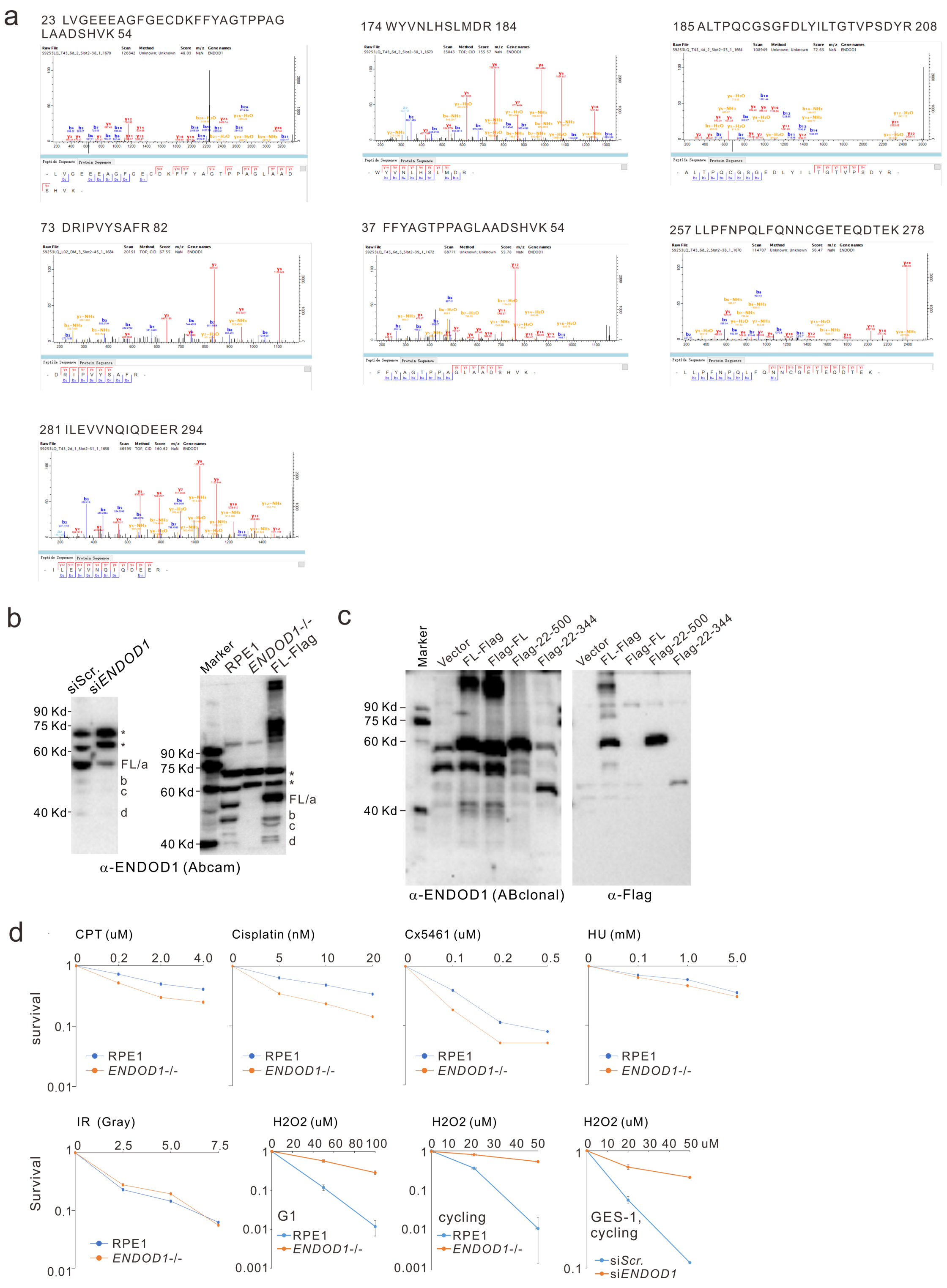
a
23 LVGEEEAGFGECDKFFYAGTPPAG LAADSHVK 54
174 WYVNLHSLMDR 184
185 ALTPQCGSGFDLYILTGTVPSDYR 208
73 DRIPVYSAFR 82
37 FFYAGTPPAGLAADSHVK 54
257 LLPFNPQLFQNNCGETEQDTEK 278
281 ILEVVNQIQDEER 294
b
siScr.
siENDOD1
90 Kd
75 Kd
60 Kd
40 Kd
*
*
FL/a
b
c
d
Marker
RPE1
ENDOD1-/-
FL-Flag
90 Kd
75 Kd
60 Kd
40 Kd
*
*
FL/a
b
c
d
α-ENDOD1 (Abcam)
c
Marker
Vector
FL-Flag
Flag-FL
Flag-22-500
Flag-22-344
Vector
FL-Flag
Flag-FL
Flag-22-500
Flag-22-344
90 Kd
75 Kd
60 Kd
40 Kd
α-ENDOD1 (ABclonal)
α-Flag
d
CPT (uM)
0 0.2 2.0 4.0
Cisplatin (nM)
0 5 10 20
Cx5461 (uM)
0 0.1 0.2 0.5
HU (mM)
0 0.1 1.0 5.0
survival
1
0.1
0.01
RPE1
ENDOD1-/-
IR (Gray)
0 2.5 5.0 7.5
H2O2 (uM)
0 20 40 60 80 100
H2O2 (uM)
0 10 20 30 40 50
H2O2 (uM)
0 10 20 30 40 50 uM
Survival
1
0.1
0.01
0.001
G1
cycling
GES-1, cycling
RPE1
ENDOD1-/-
siScr.
siENDOD1

**Supplementary Figure 2.**

**a.** Enlarged image as in Figure 1b showing overlapping $H_2O_2$-induced foci of PAR and ENDOD1 in RPE1 cells. Representative image from 3 biological repeats. **b.** ENDOD1 foci in RPE1 cells with or without pretreatment with indicate siRNA or the PARPi Talazoparib. **c.** Time course for PAR foci formation in cycling or serum starved (G1 arrested) *ENDOD1*$^{-/-}$ and control RPE1 cells. n = 5 biologically independent samples. Error bars: standard error of the mean. **d**. Relative proliferation of *ENDOD1*$^{-/-}$ and control RPE1 cells with or without continuous $H_2O_2$ treatment (10 μM). Cells were treated with the indicated siRNAs 48 hours before challenge. n = 3 biologically independent samples. Error bars: standard error of the mean. **e**. Relative proliferation of *ENDOD1*$^{-/-}$ and control RPE1 cells with or without continuous $H_2O_2$ treatment (10 ∝M) and co-treatment with Olaparib (200 uM) or Talaparib (20 nM) for 72 hours. n = 3 biologically independent samples. Error bars: standard error of the mean. All significance tests: two-tailed Student's *t* test. n.s. not significant.

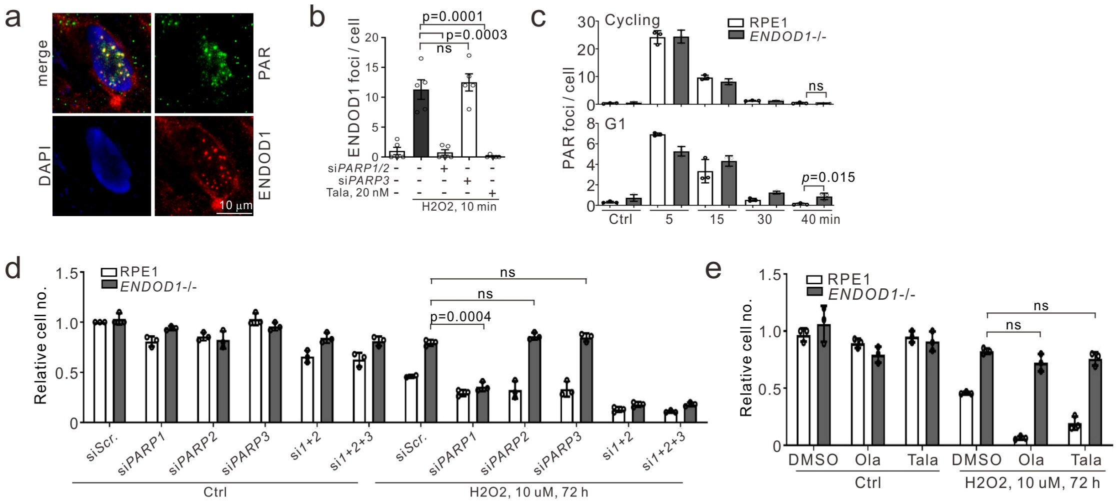

**Supplementary Figure 3.**

**a**. Top: SL between si*ENDOD1* and si*BRCA1* validated with three additional si*ENDOD1* sequences (si002-004). Representative image from 3 biological repeats. Bottom: knockdown efficacies of each si*ENDOD1* is shown by immunoblotting with α-ENDOD1 (Abcam). **b**. Quantification of 53BP1 and γH2AX foci in *ENDOD1*$^{-/-}$ and control RPE1 cells following either control (si*Scr.*) or si*BRCA1* treatment. n = 3 biologically independent samples. Inset: semi-quantitative PCR showing knockdown efficiencies of *BRCA1*. **c.** Representative images for experiments described in Figure 2c. **d**. Left: relative survival for si*BRCA1* treated *ENDOD1*$^{-/-}$ cells co-treated with either Olaparib (20 nM) or Talazoparib (2 nM) n = 3 biologically independent samples. Right: quantification of 53BP1 foci in si*BRCA1* or control (si*Scr.*) treated *ENDOD1*$^{-/-}$ cells co-treated with eiher Olaparib (100 nM) or Talaparib (10 nM). n = 3 biologically independent samples **e**. SL with ENDOD1 and HRD recapitulated in an HRD cancer cell line, MCF-7. Cell proliferation was followed by Haemocytometer counting of cells treated at 3 day intervals with control (si*Scr.*) or si*ENDOD1*. n = 3 biologically independent samples Knockdown efficiency is shown by immunoblotting with a-ENDOD1 (Abcam). **f.** 53BP1 foci formation (left) and quantification (right) in MCF-7 cells treated with the indicated siRNAs. n = (51-59) x 5 cells (each data point represents the average foci number of 5 cells). All error bars; standard error of the mean. All significance tests: two-tailed Student's *t* test. n.s. not significant.

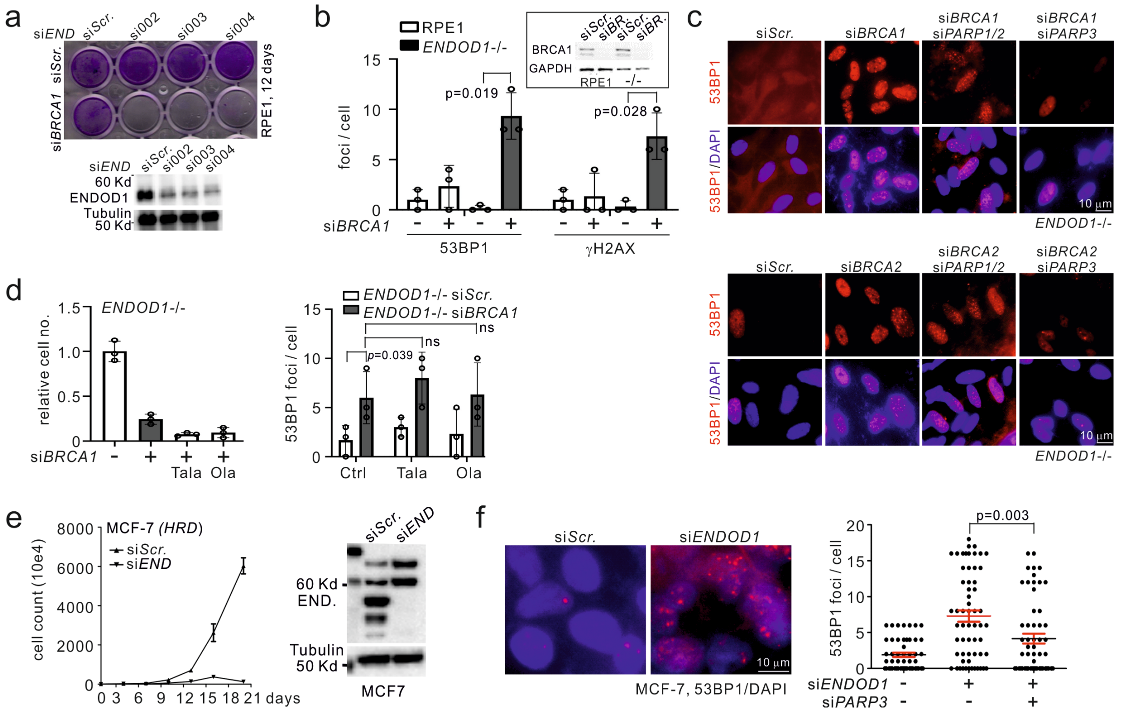

**Supplementary Figure 4.**

**a**. The effect of si*ENDOD1* treatment on *TP53* mutated cancer cells (extending the data shown in Figure 3a). Proliferation curves following either si*ENDOD1* or control (si*Scr.*) transfection. At 3-day intervals cells were passaged, counted by haemocytometer and transfected. n = 3 biologically independent samples. Reported *TP53* status is indicated for each cell line[1] (see also: https://www.lgcstandards-atcc.org). Cell numbers for NCI-H1299, HL60 and MDA-MB-231 were counted only once, 10 days after the initial transfection. The cervical cancer line (HeLa) expresses HPV-E6 protein that functionally inactivates p53[2]. **b.** Relative cell viability as measured by CCK8 colorimetry at the indicated time points for C33A cells treated with two independent siRNAs against *ENDOD1*. n = 3 biologically independent samples. Significance test: two-tailed Student's *t* test. Right: immunoblotting evaluating knockdown efficacies. **c**. Crystal blue staining of *ENDOD1*$^{-/-}$ and control RPE1 cells following treatment with the indicated siRNAs. Cells were transfected at 3-day intervals and stained on day 10 unless otherwise indicated. **d**. *TP53* knockdown in *ENDOD1* null cells results in cell death by apoptosis. Left/middle: cell death assayed by TUNEL for the indicated cells 6 days after control (si*Scr.*) or si*TP53* transfection. Right: abnormal nuclear morphology revealed by Giemsa staining. Representative images from 1 of three independent experiments. **e**. *TP53* knockdown in *ENDOD1* null cells results in G1 arrest. FACS analysis for *ENDOD1*$^{-/-}$ and control RPE1 cells following the indicated siRNA treatment. Representative of three independent experiments. **f.** *ENDOD1*$^{-/-}$ cells treated with si*TP53* and concomitantly infected with control (pLV) or pLV-*ENDOD1* (END) lentivirus. Cell viability were determined by CCK8 colorimetry 7 days after transfection. n = 3 biologically independent samples. Right: detection of indicated proteins by immunoblotting. **g.** Denatured BrdU staining after 60 min chase, to show DNA synthesis of RPE1 and *ENDOD1*$^{-/-}$ cells with or without serum starvation. Representative of three independent experiments. **h.** Left: immunoblot showing knockdown of *ENDOD1* with two different siRNAs targeting *ENDOD1* (si001 and si002) in two congenic HCT116 cell lines carrying wild type (Wt) or mutant (mu) form of p53 (Δ40p53). Right cell viability counted by haemocytometer at the indicated time points (right). n = 3 biologically independent samples. All error bars: standard error of the mean. All significance tests: two-tailed Student's *t* test.

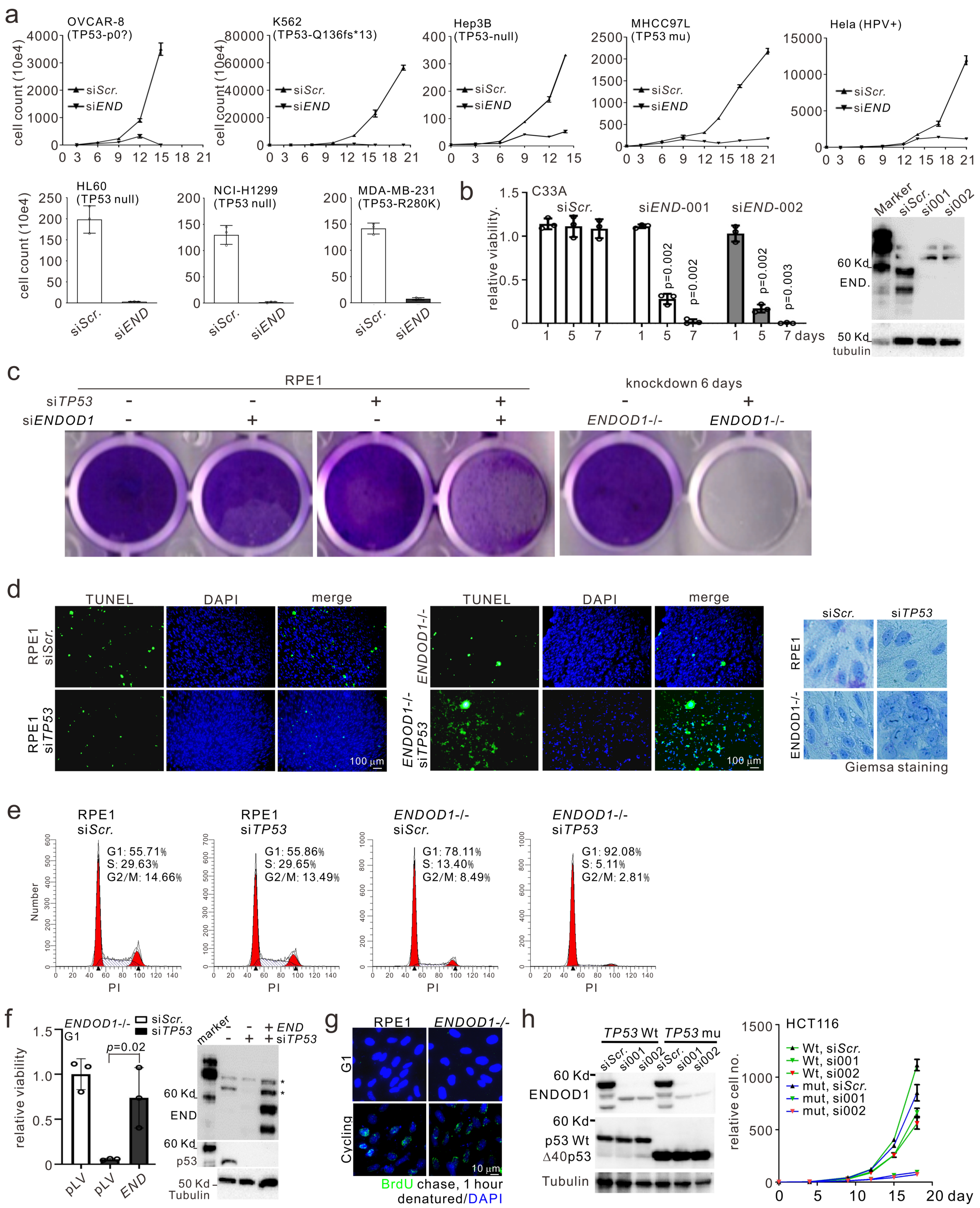
a
OVCAR-8 (TP53-p0?)
K562 (TP53-Q136fs*13)
Hep3B (TP53-null)
MHCC97L (TP53 mu)
Hela (HPV+)
HL60 (TP53 null)
NCI-H1299 (TP53 null)
MDA-MB-231 (TP53-R280K)
cell count (10e4)
siScr.
siEND
b
C33A
relative viability
siEND-001
siEND-002
7 days
c
RPE1
knockdown 6 days
siTP53
siENDOD1
ENDOD1-/-
d
TUNEL
DAPI
merge
Giemsa staining
e
PI
f
relative viability
g
BrdU chase, 1 hour denatured/DAPI
h
TP53 Wt
TP53 mu
ENDOD1
Tubulin
HCT116
relative cell no.

**Supplementary Figure 5.**

**a.** Representative images of alkaline (left) or neutral (right) comet assays for one of three biological repeats of the serum starved (G1 arrested) *ENDOD1*$^{-/-}$ and control RPE1 cells with or without prior si*TP53* transfection as described in Figure 4a. **b**. Quantifications for pRPA32 foci in *ENDOD1*$^{-/-}$ and control RPE1 cells in serum-starved (G1 arrested) or cycling cells, 72 hours after si*TP53* treatment. n = 100 x 5 cells (each data point represents the average foci number of 5 cells). **c.** Detection of ssDNA in ENDOD1$^{-/-}$ cells (G1 phase) as in Figure 4c reproduced with a second *TP53* siRNA (si002). Representative images from 1 of three independent experiments. **d**, Quantification of ssDNA upon depletion of TP53 with or without re-introducing ENDOD1 via lentiviral infection (control, pLV; ENDOD1 expression, END). n = 3 biologically independent samples. arb.units: arbitrary units. **e**. 150 ng purified genomic DNA from indicated cells digested by S1 nuclease (0.02 U/$\mu$l) for 10 min. DNA was resolved by 1% agarose electrophoresis. C = control (no nuclease). See Figure 4e. Representative images from 1 of three independent experiments. **f**. Quantification of 53BP1 and $\gamma$H2AX foci in serum starved (G1 arrested) *ENDOD1*$^{-/-}$ and control RPE1 cells 72 hours after control or si*TP53* treatment. n = 3 biologically independent samples. **g**. Quantifications of pRPA32 and 53BP1 foci in serum starved *ENDOD1*$^{-/-}$ and control RPE1 cells 0, 48, 72 and 96 hours after control (si*Scr.*) or si*TP53* treatment. n = 3 biologically independent samples. **h**. $\alpha$-p53 immunoblot analysis upon ectopic expression of *TP53* wild type and individual alleles in SKOV-3 cells analysed in Figure 4g and Figure 5e. Representative of three independent experiments. All error bars; standard error of the mean. All significance tests: two-tailed Student's *t* test.

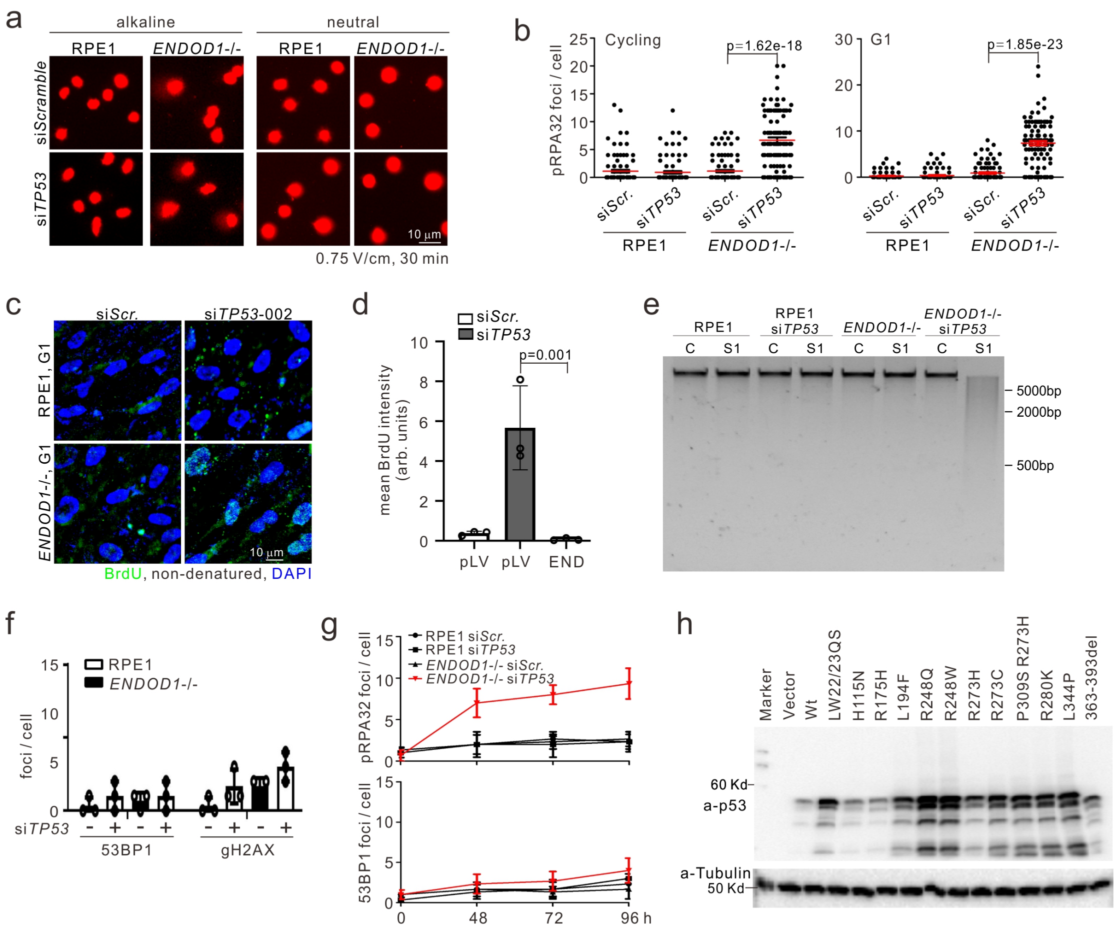

a
alkaline
neutral
RPE1
ENDOD1-/-
RPE1
ENDOD1-/-
siScramble
siTP53
10 µm
0.75 V/cm, 30 min
b
pRPA32 foci / cell
Cycling
p=1.62e-18
G1
p=1.85e-23
siScr.
siTP53
siScr.
siTP53
RPE1
ENDOD1-/-
c
siScr.
siTP53-002
RPE1, G1
ENDOD1-/-, G1
10 µm
BrdU, non-denatured, DAPI
d
siScr.
siTP53
p=0.001
mean BrdU intensity (arb. units)
pLV
pLV
END
e
RPE1
RPE1 siTP53
ENDOD1-/-
ENDOD1-/- siTP53
C
S1
C
S1
C
S1
C
S1
5000bp
2000bp
500bp
f
RPE1
ENDOD1-/-
foci / cell
siTP53
53BP1
gH2AX
g
pRPA32 foci / cell
RPE1 siScr.
RPE1 siTP53
ENDOD1-/- siScr.
ENDOD1-/- siTP53
53BP1 foci / cell
0
48
72
96 h
h
Marker
Vector
Wt
LW22/23QS
H115N
R175H
L194F
R248Q
R248W
R273H
R273C
P309S R273H
R280K
L344P
363-393del
60 Kd
a-p53
a-Tubulin
50 Kd

**Supplementary Figure 6.**

**a-b**. Representative images (a) and quantifications (b) of pRPA32 staining for the indicated cancer cells treated with control or si*ENDOD1* and co-treated with siRNAs for the indicated PARPs. n = 3 biologically independent samples. **c-d**. Quantification and images of pRPA32 and PAR foci in serum starved (G1 arrested) SKOV-3 (*TP53* null) cells in response to si*ENDOD1* treatment. The signals can be complemented with either siRNA-resistant *ENDOD1* expression (END). (c. n = 5 biologically independent samples. arb.units: arbitrary units.) or by wild type *TP53* expression (oe*TP53*) (d. Representative images from 1 of three independent experiments). **e**. Viability for SKOV-3 cells treated with si*ENDOD1*, complemented or not (pLV) by lentivirus encoding *ENDOD1* (siRNA resistant: END) or wild type *TP53*. n = 3 biologically independent samples. **f.** Quantifications of pRPA32 foci in the indicated serum-starved (G1 arrested) cancer lines following si*ENDOD1* and co-treated, or not, with PARPi's. n = 5 biologically independent samples **g.** Detection of ssDNA by non-denatured IF for BrdU in si*TP53* treated *ENDOD1*$^{-/-}$ (G1 arrested) cells co-treated, or not, with either si*PARP1* or PARPi. n = 3 biologically independent samples. arb.units: arbitrary units. **h.** S1 nuclease-sensitivity assay for genomic DNA extracted from si*TP53* treated seum starved (G1-arrested) *ENDOD1*$^{-/-}$ cells co-treated, or not, with PARPi or si*PARP1*. Representative images from 1 of three independent experiments. All error bars: standard error of the mean. All significance tests: two-tailed Student's *t* test.

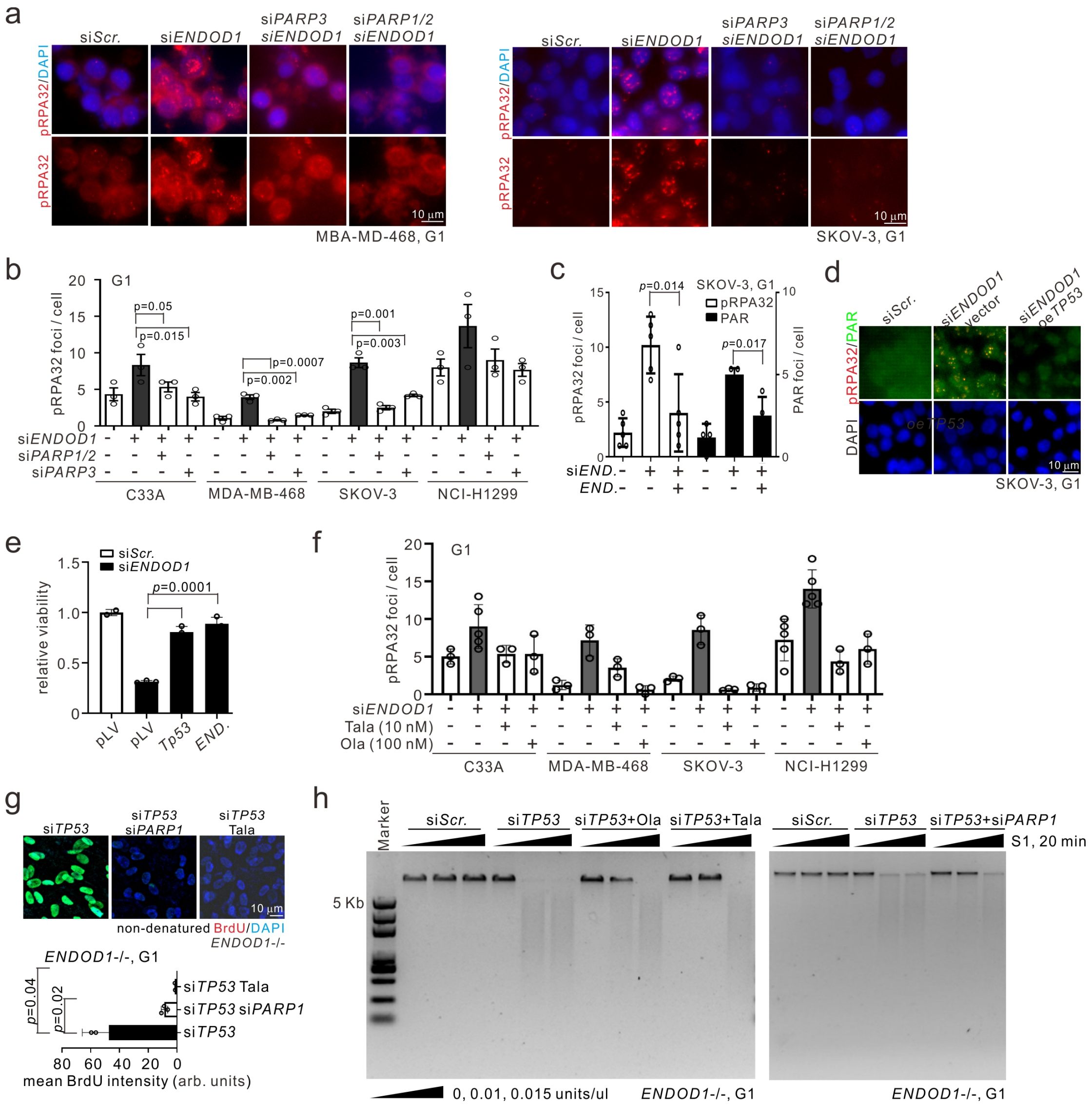
a
siScr.
siENDOD1
siPARP3 siENDOD1
siPARP1/2 siENDOD1
pRPA32/DAPI
pRPA32
10 μm
MBA-MD-468, G1
SKOV-3, G1
b
G1
pRPA32 foci / cell
p=0.05
p=0.015
p=0.0007
p=0.002
p=0.001
p=0.003
siENDOD1
siPARP1/2
siPARP3
C33A
MDA-MB-468
SKOV-3
NCI-H1299
c
p=0.014
SKOV-3, G1
pRPA32
PAR
p=0.017
PAR foci / cell
siEND.
END.
d
siScr.
siENDOD1 vector
siENDOD1 oeTP53
pRPA32/PAR
DAPI
SKOV-3, G1
e
siScr.
siENDOD1
p=0.0001
relative viability
pLV
pLV
Tp53
END.
f
G1
pRPA32 foci / cell
siENDOD1
Tala (10 nM)
Ola (100 nM)
C33A
MDA-MB-468
SKOV-3
NCI-H1299
g
siTP53
siTP53 siPARP1
siTP53 Tala
10 μm
non-denatured BrdU/DAPI
ENDOD1-/-
ENDOD1-/-, G1
p=0.04
p=0.02
siTP53 Tala
siTP53 siPARP1
siTP53
mean BrdU intensity (arb. units)
h
Marker
siScr.
siTP53
siTP53+Ola
siTP53+Tala
siScr.
siTP53
siTP53+siPARP1
S1, 20 min
5 Kb
0, 0.01, 0.015 units/ul
ENDOD1-/-, G1
ENDOD1-/-, G1

**Supplementary Figure 7.**

**a**. Quantification of pRPA32 foci formation in si*TP53* treated *ENDOD1*$^{-/-}$ cells (G1-arrested) with co-treatment using control (si*Scr.*) or siRNA against the indicated gene. n = 3 biologically independent samples. **b**. PAR and pRPA32 foci in si*TP53* treated *ENDOD1*$^{-/-}$ cells co-treated with either control (si*Scr.*) or si*XRCC1*. Representative of three independent experiments. **c**. Representative images (left) and quantification (right) of the indicated resection factor foci in serum-starved and si*TP53* treated *ENDOD1*$^{-/-}$ cells. n = 3 biologically independent samples. Note foci are shown separately for phospho- and non-phospho-BRCA1 and NBS1. **d**. Quantification for PAR and MRE11 foci in si*TP53* treated *ENDOD1*$^{-/-}$ cells co-treated with either control (s*iScr.*) or si*XRCC1*. n = 3 biologically independent samples. All error bars: standard error of the mean. Significance test: two-tailed Student's *t* test.

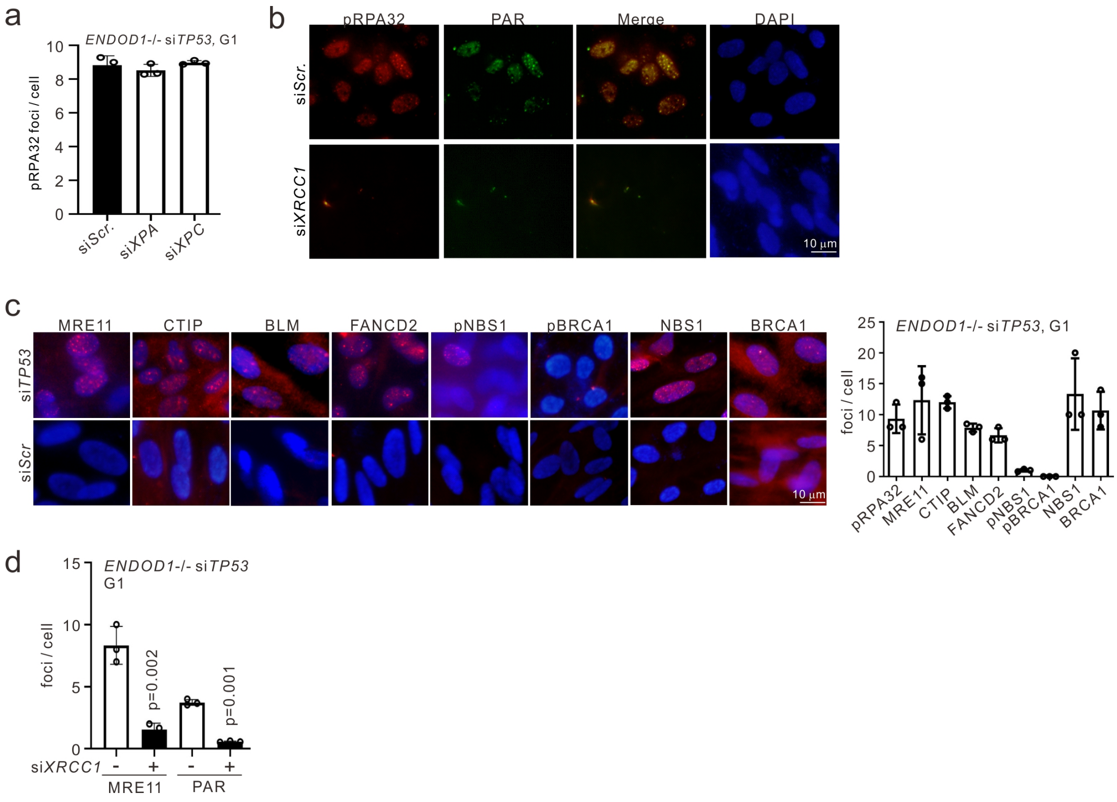

**Supplementary Figure 8.**

**a.** Semi-quantitative PCR for m*Endod1* or m*Wdr70* in the indicated tissues from individual animals subjected to 3 or 2 month *in vivo* knockdown, respectively. representative individual animals shown: n = 6 and 5 for siScr and simEnd, respectively. **b.** Echocardiographic parameters charactering heart function after two-month knockdown of the three treatment groups. Indicators of ventricular wall thickness (millimeter (mm), left panel) and cardiac systolic function (right panel) are shown. LVIDd, left ventricular internal diameter at end diastole. LVPWd: left ventricular posterior wall end diastole. LVPWs: left ventricular posterior wall end systole. EF: ejection fraction. FS: fraction shortening. n = 6,5 and 2 for siScr, simEnd and simWdr70 respectively. **c**. Myeloid and lymphoid populations in bone marrow assessed by FACS sorting. n = 6,5 and 2 for siScr, simEnd and simWdr70 respectively. The antibody conjugates are given in parentheses. **d.** Tumour volume measured for MDA-MB-361 xenografts treated, or not, with si*ENDOD1*. All error bars: standard error of the mean. All significance tests: two-tailed Student's *t* test. n.s. not significant.

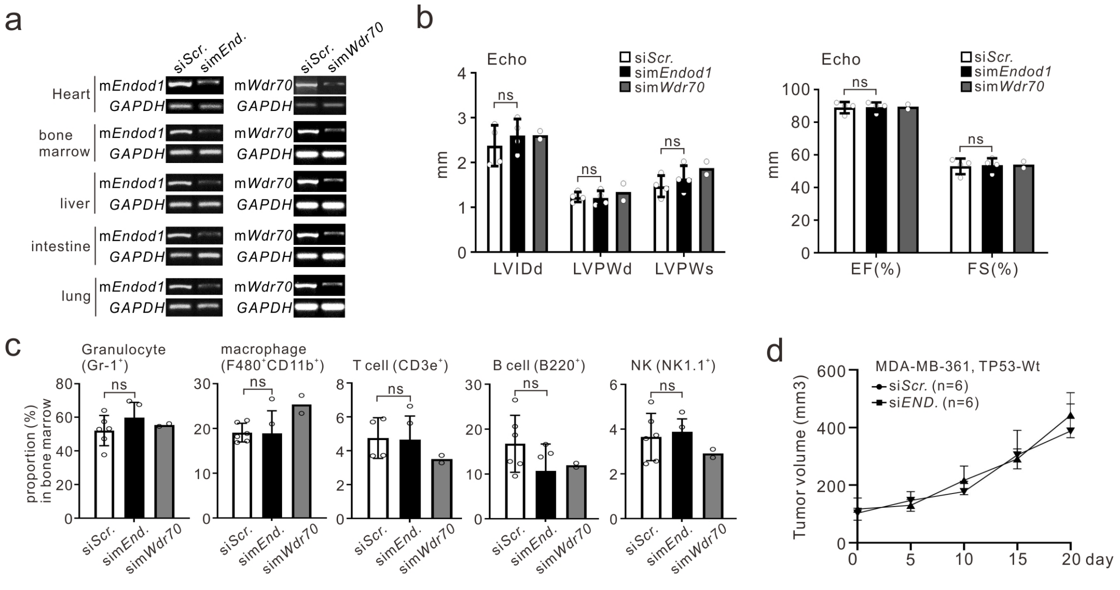

**Supplementary Figure 9.**

**a**. FACS analysis of the indicated cycling and serum starved cells (see Supplementary Methods). **b**. The gating FACS gating is exemplified by the strategy used to identify hematopoietic stem cell (HSCs) progenitors is shown. The areas delineated by the solid black lines in the scatter plots indicate the population of cells gated. Dashed arrows point to further gating of these cells. Lin-C-kit+Sca-1+ cell population was defined as HSCs. Lineage markers (Lin) included CD3e, B220, CD11b, Gr-1 and Ter-119. SSC, side light scatter; FSC, forward light scatter. **c**. Schematic for CRISPR targeting Exon1 of the *ENDOD1* locus in RPE1 cells. gRNA sequence and the single insertion of a guanine base are shown.

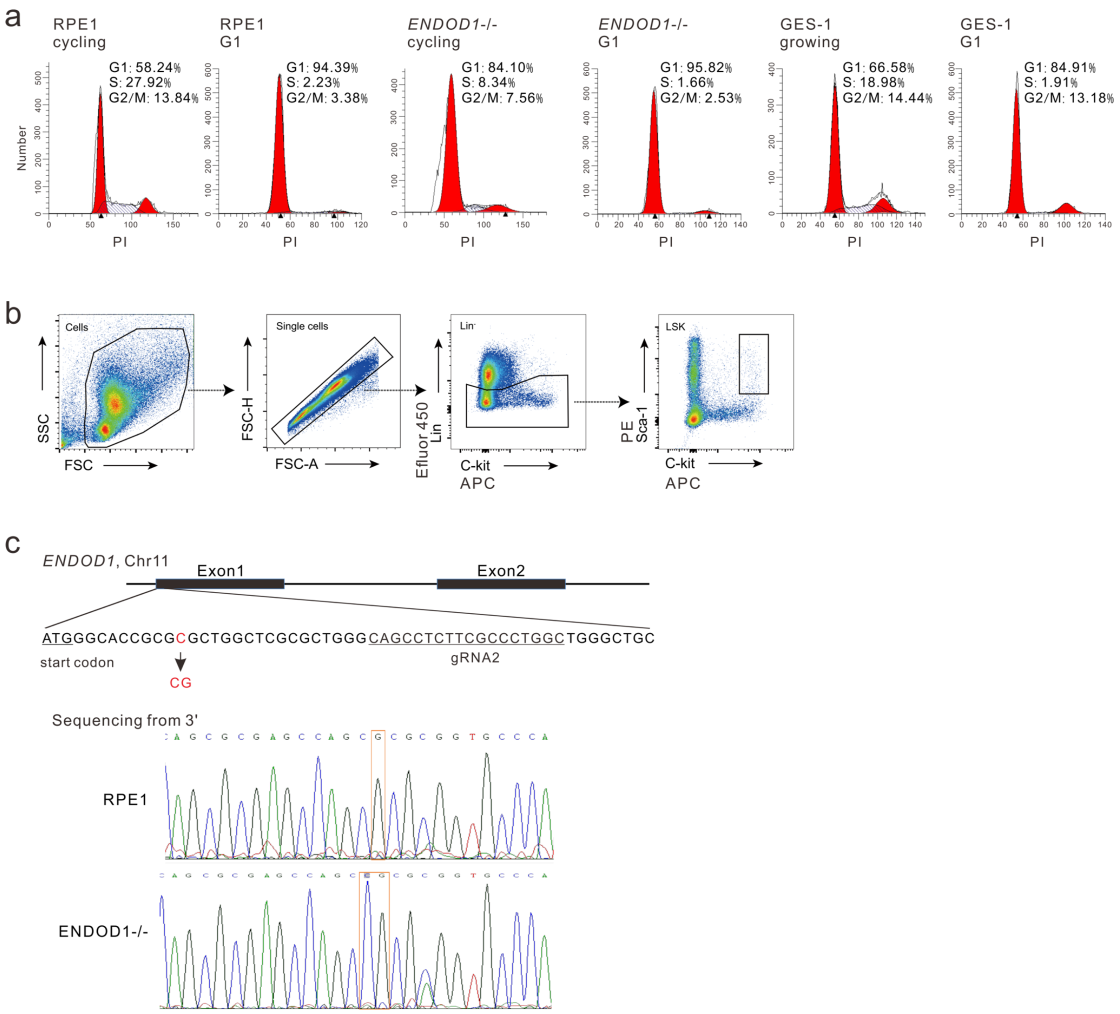

**Supplementary Figure 10.**

**a.** Immunoblotting validating siRNA knockdown efficiencies for si*PARP1-3.* Representative of three independent experiments. **b**. Immunoblotting validating siRNA knockdown of *TP53* in RPE1 and *ENDOD1*$^{-/-}$ and the impact of *TP53* silencing on transactivation assessed by p21 levels. SKOV-3 serves as *TP53* null control. Representative of three independent experiments. **c.** Semi-quantitative PCR and immunoblotting showing knockdown efficiencies for indicated genes. Representative of 2 – 3 independent experiments. **d**. Immunoblotting validating siRNA knockdown efficiencies in different cell lines for si*ENDOD1.* Black arrows indicate the full length of ENDOD1. Representative of 2 – 3 independent experiments.

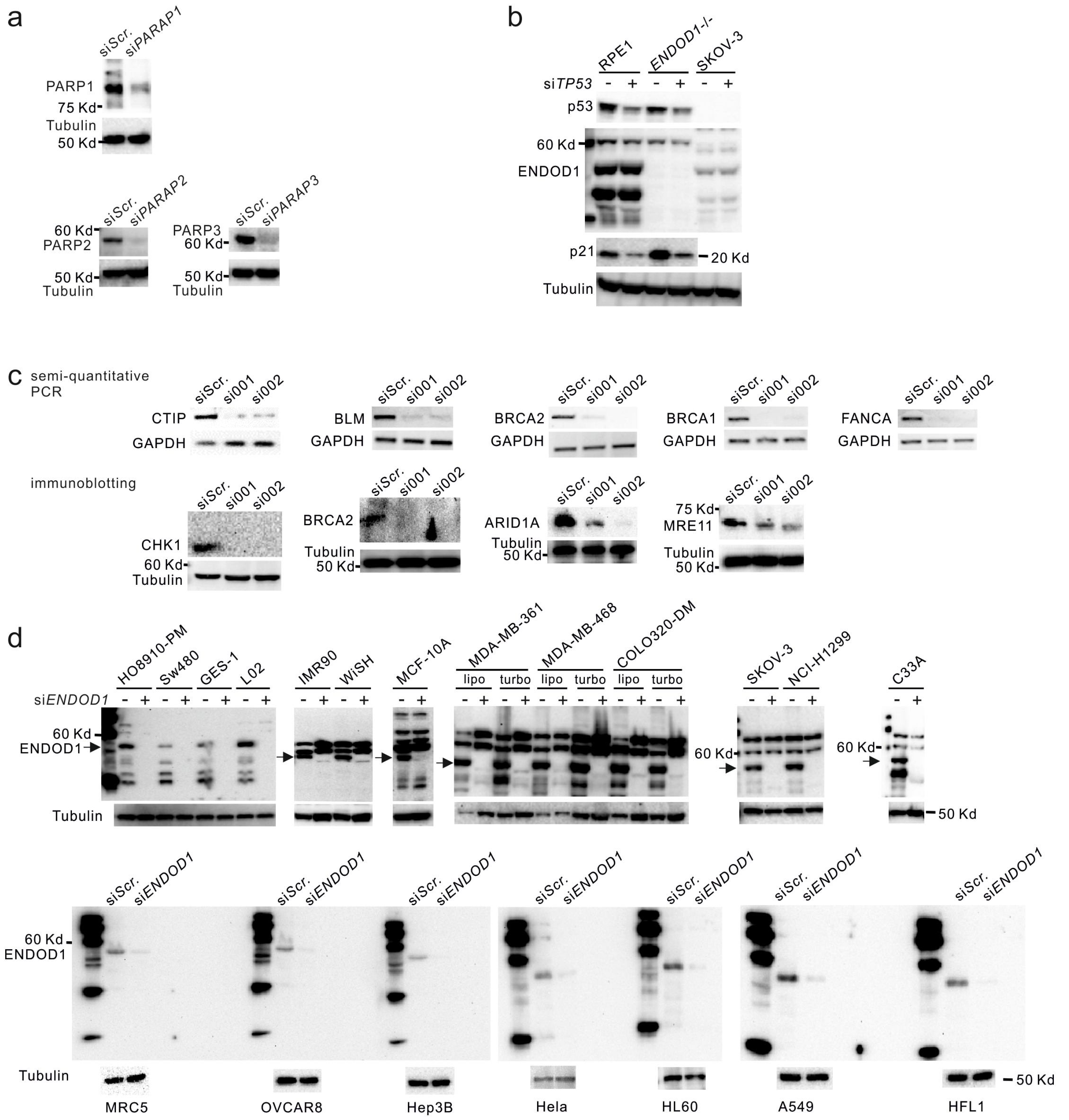

**Supplementary Figure 11**

Uncropped blots relating to the indicated figures.

Fig 1d

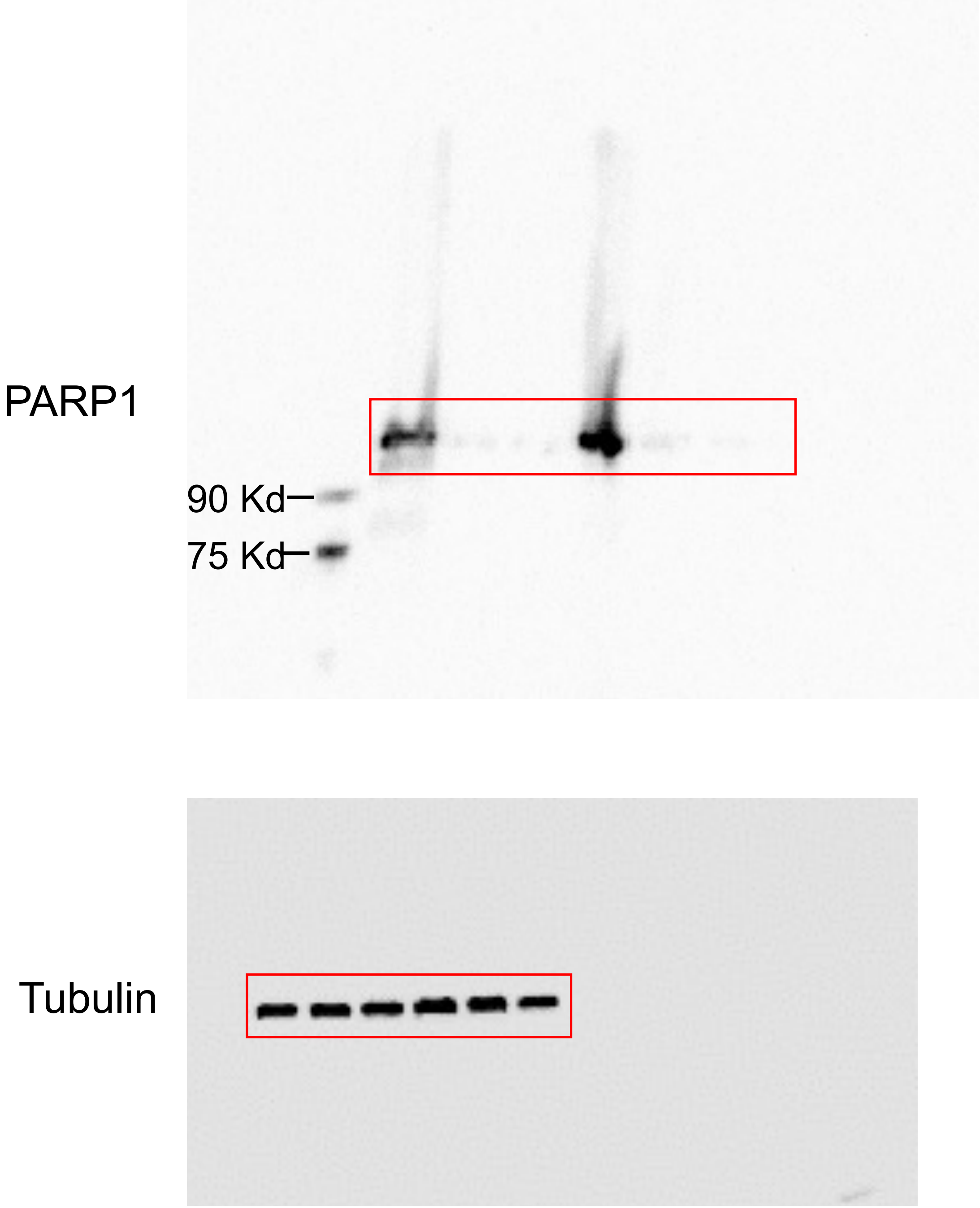


Protein size was estimated by prestained Precision Plus Protein Standards(Bio-Rad, 161-0374) of 50 Kd.

Fig 1e

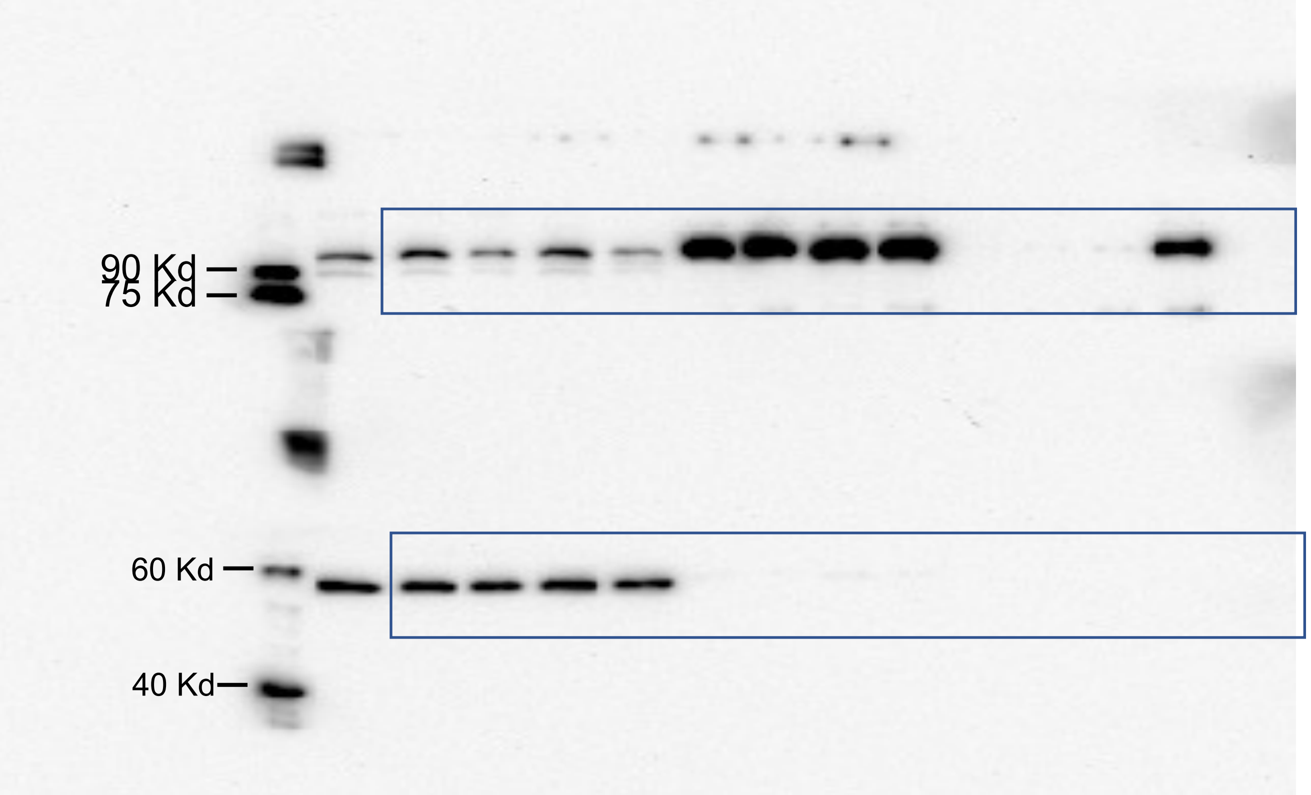


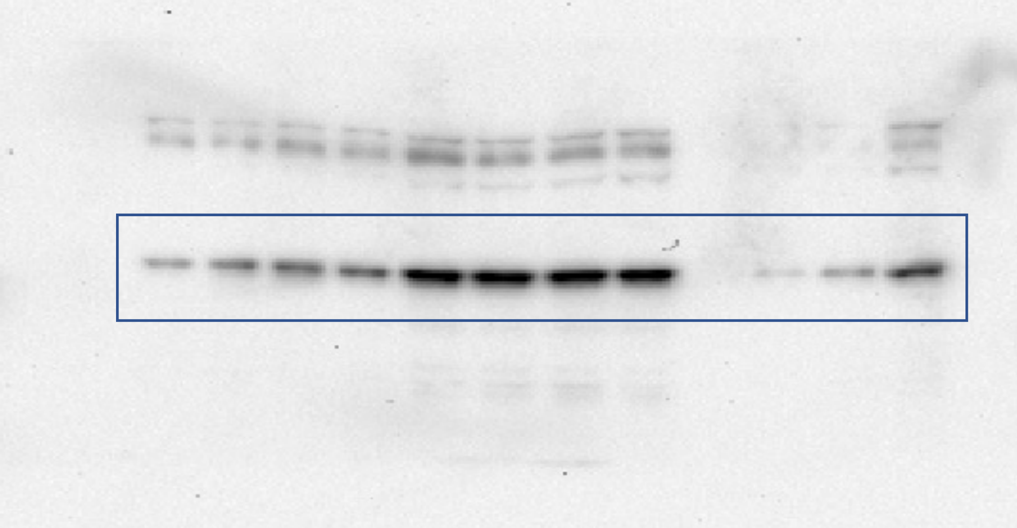


Protein size was estimated by prestained Precision Plus Protein Standards(Bio-Rad, 161-0374) of 50 Kd.

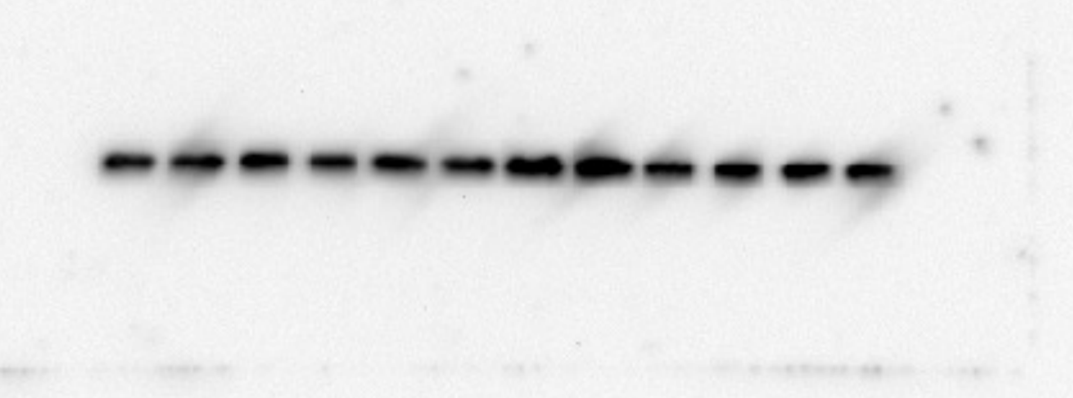


The ECL marker was cut off in this blot. Protein size was estimated by prestained Precision Plus Protein Standards(Bio-Rad, 161-0374) of 15 Kd.

Fig 2c

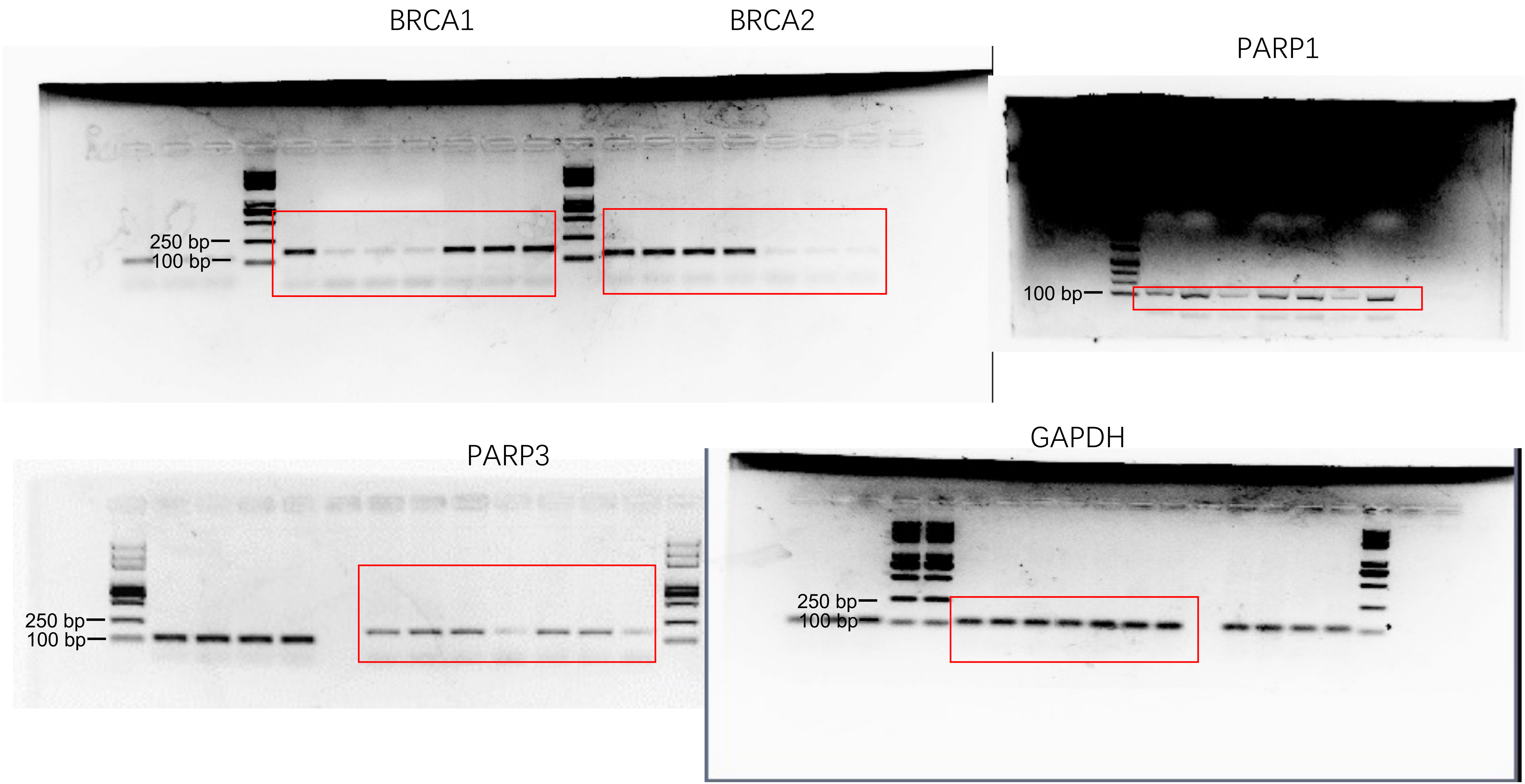
BRCA1
BRCA2
PARP1
250 bp
100 bp
100 bp
PARP3
GAPDH
250 bp
100 bp
250 bp
100 bp

Fig 5b

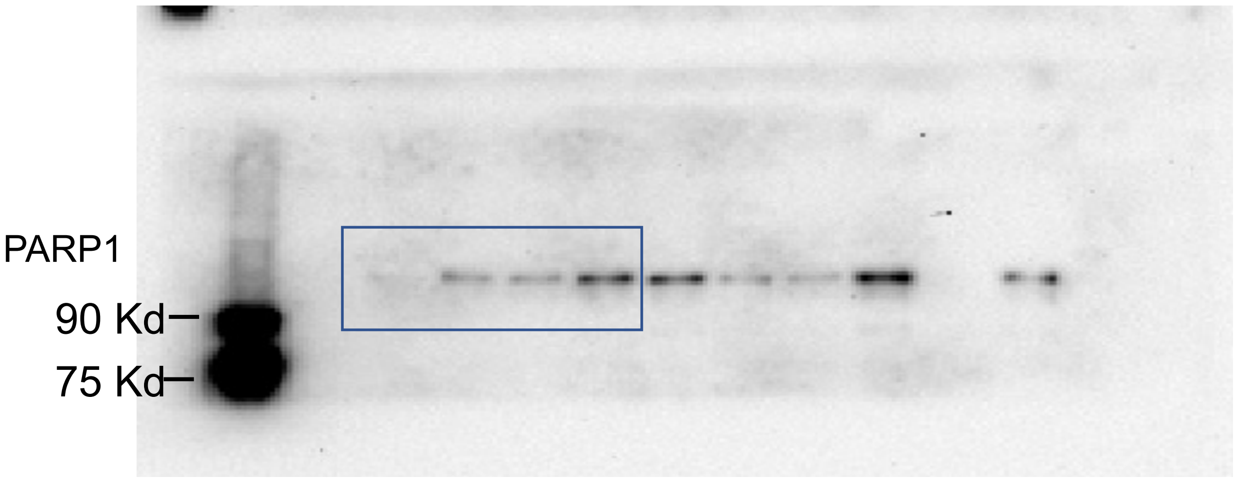


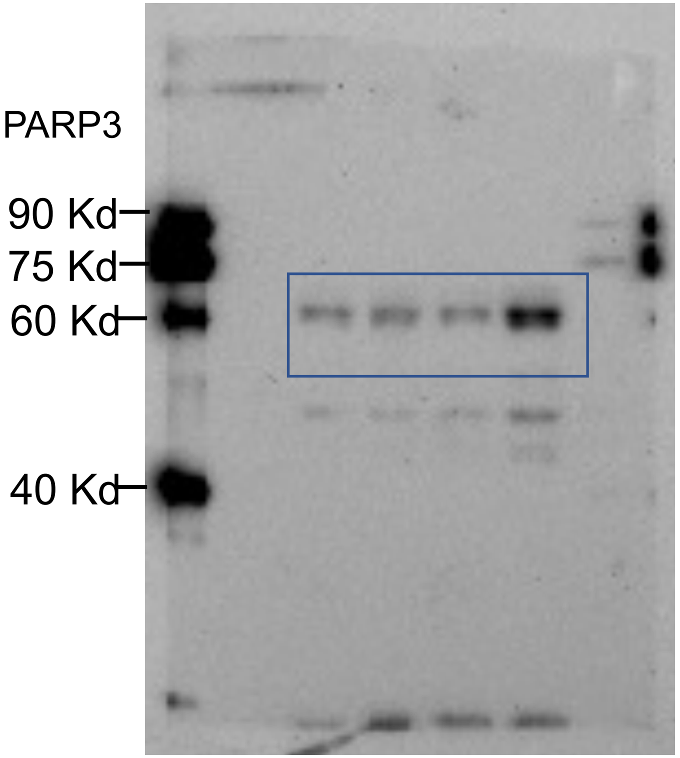


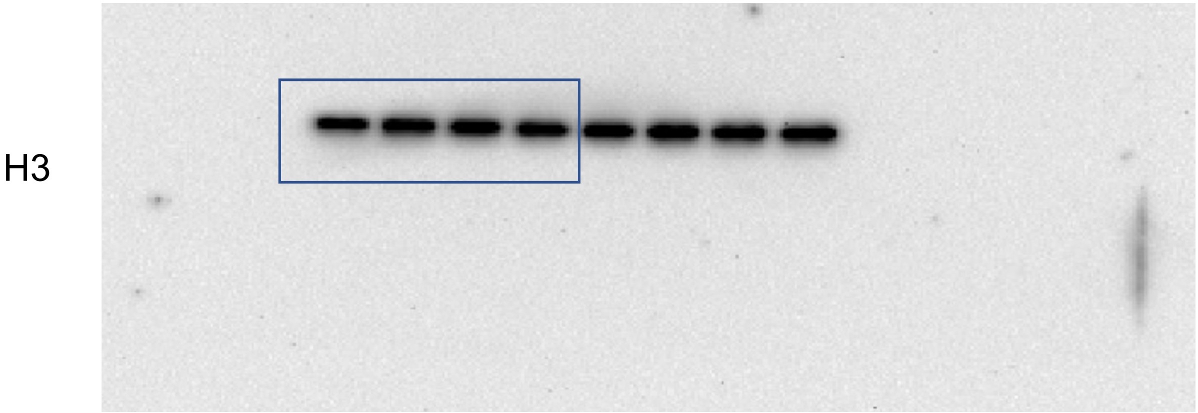


The ECL marker was cut off in this blot. Protein size was estimated by prestained Precision Plus Protein Standards(Bio-Rad, 161-0374) of 15 Kd.

Fig 5c

p53

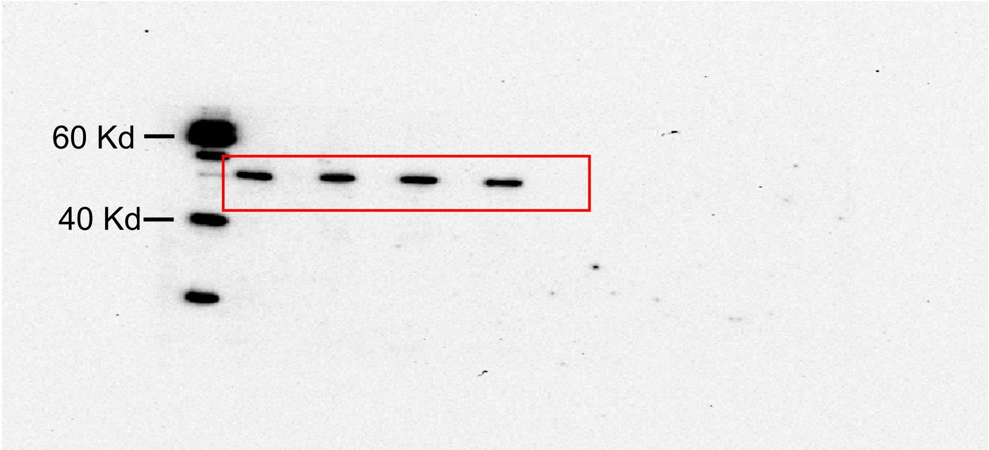


Tubulin

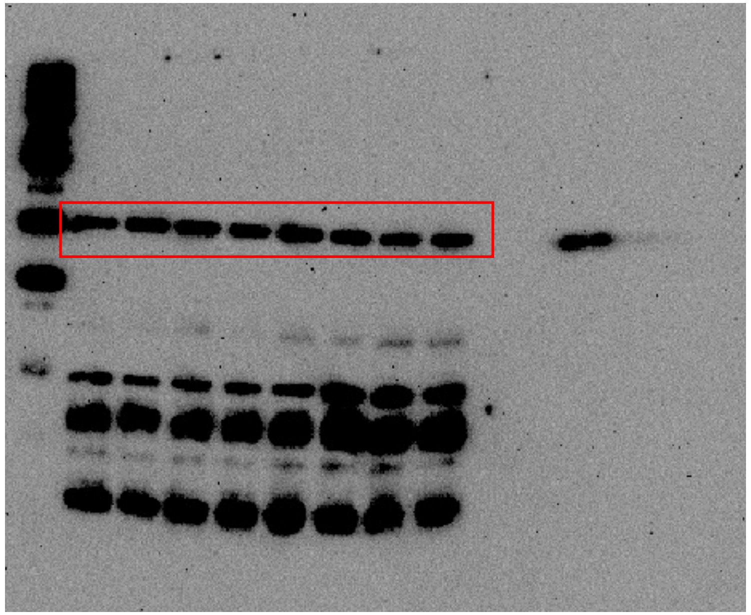

Fig 5h

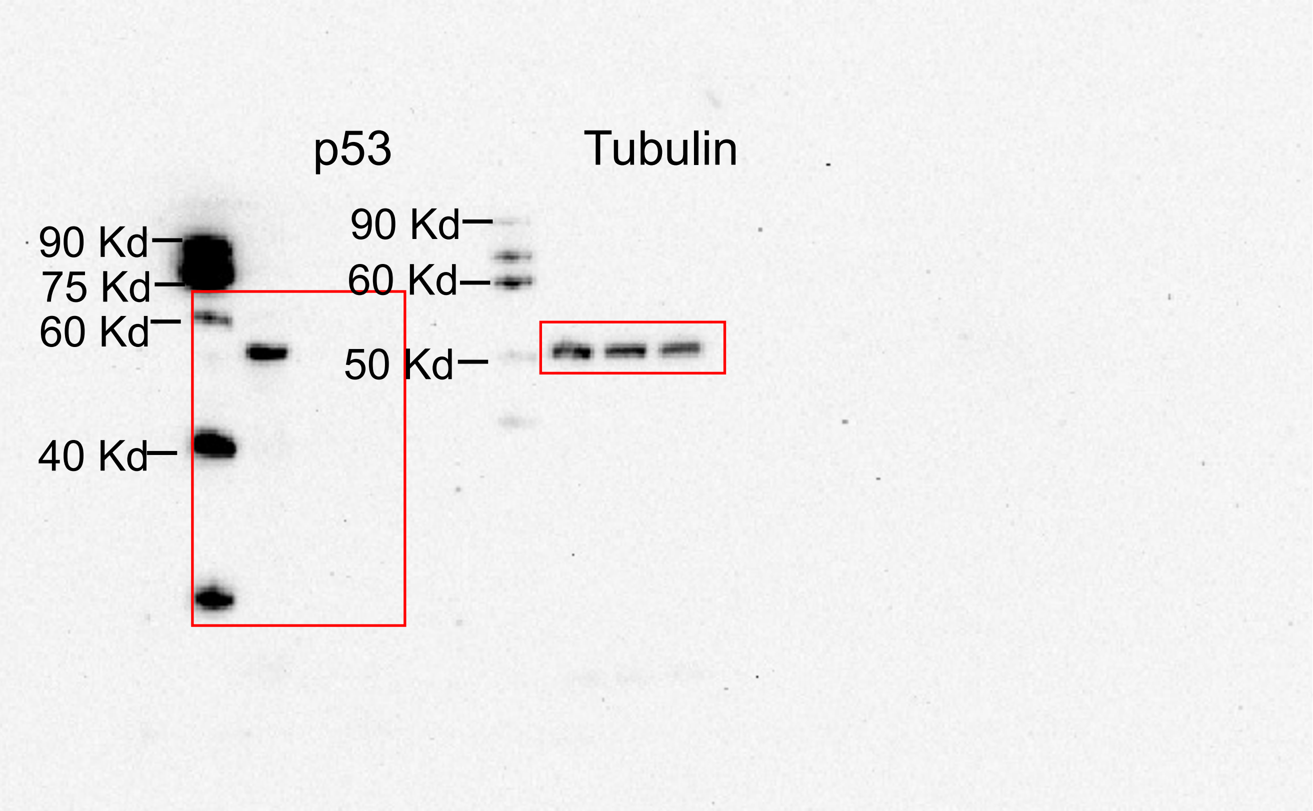

Fig S1b

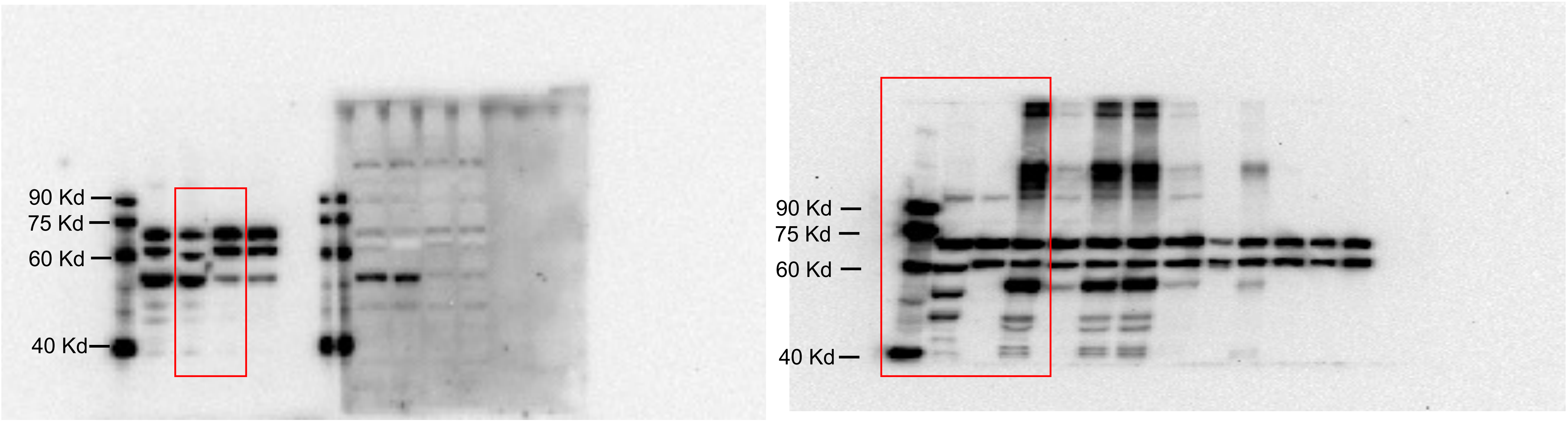


ENDOD1

ENDOD1

Fig S1c

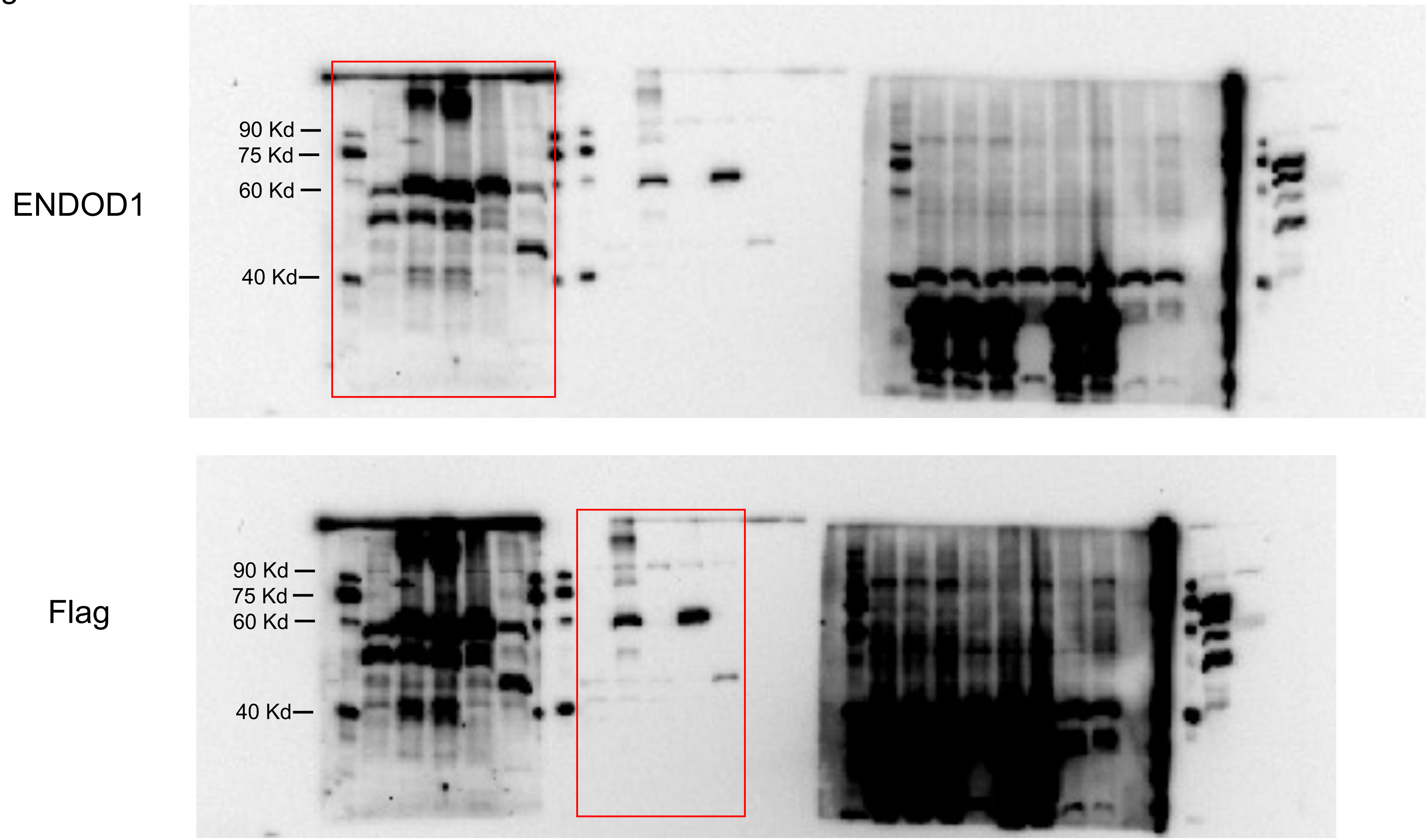

Fig S3a

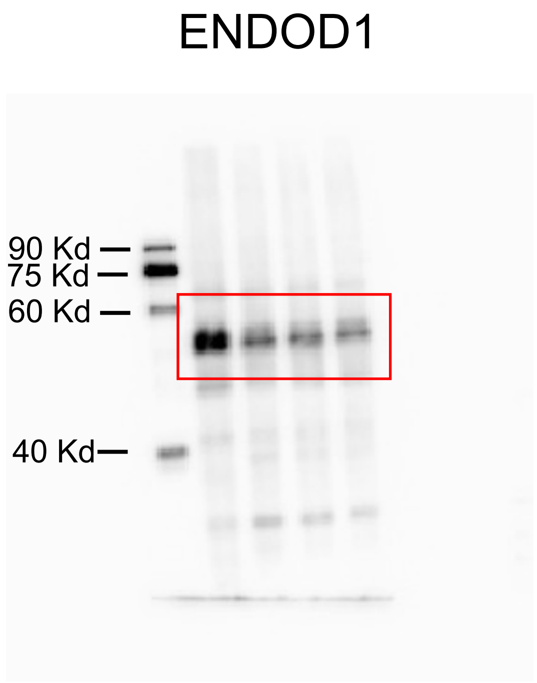


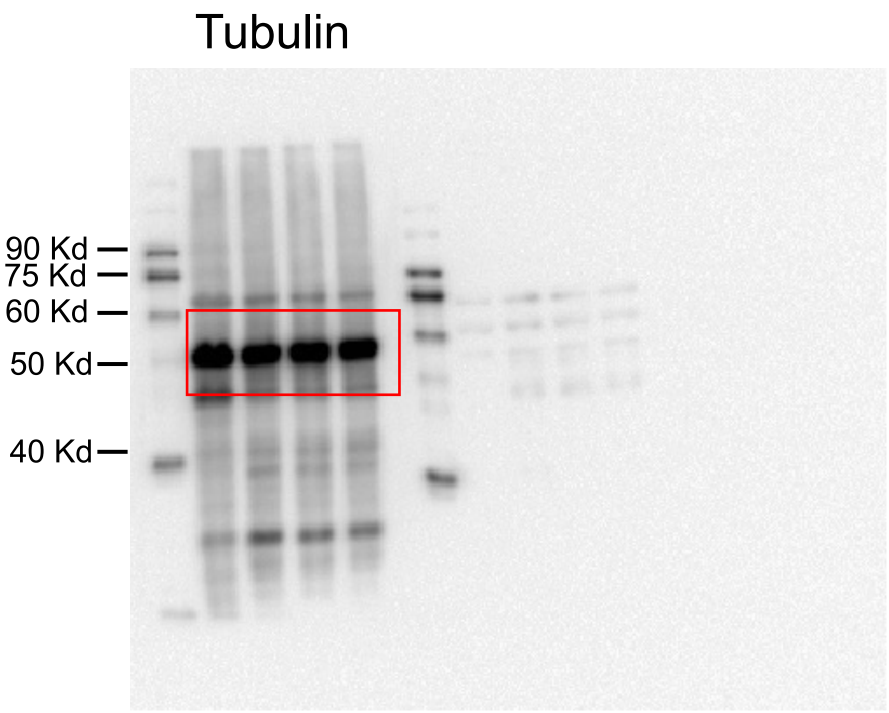

Fig S3b

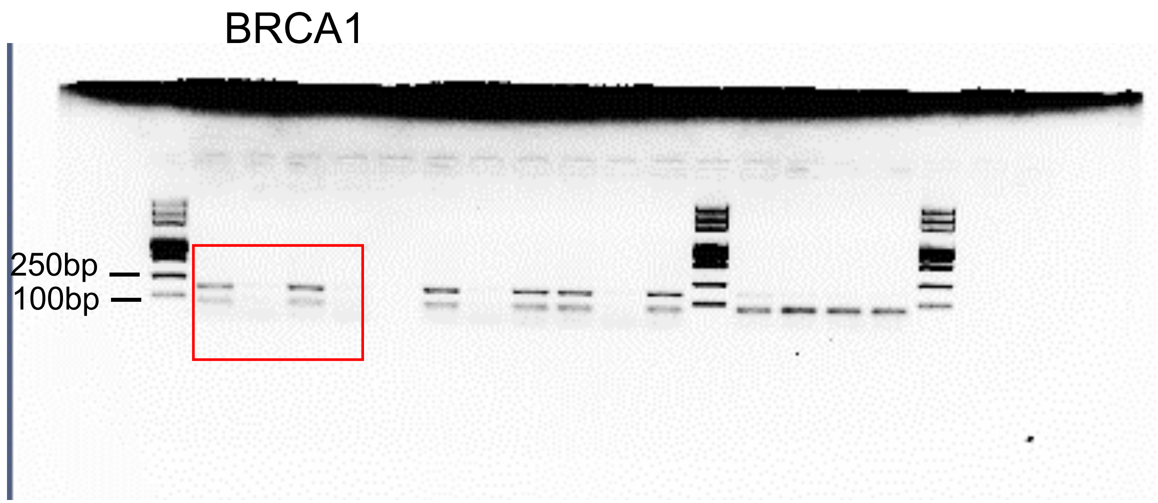


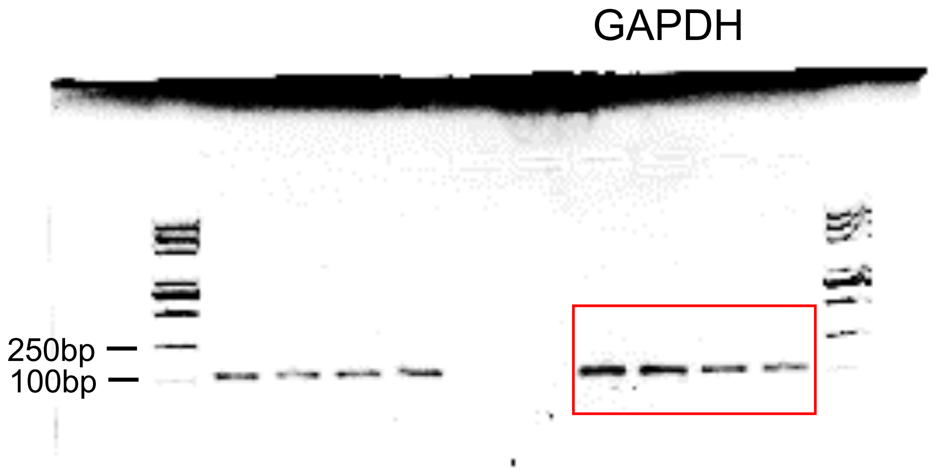

Fig S3e

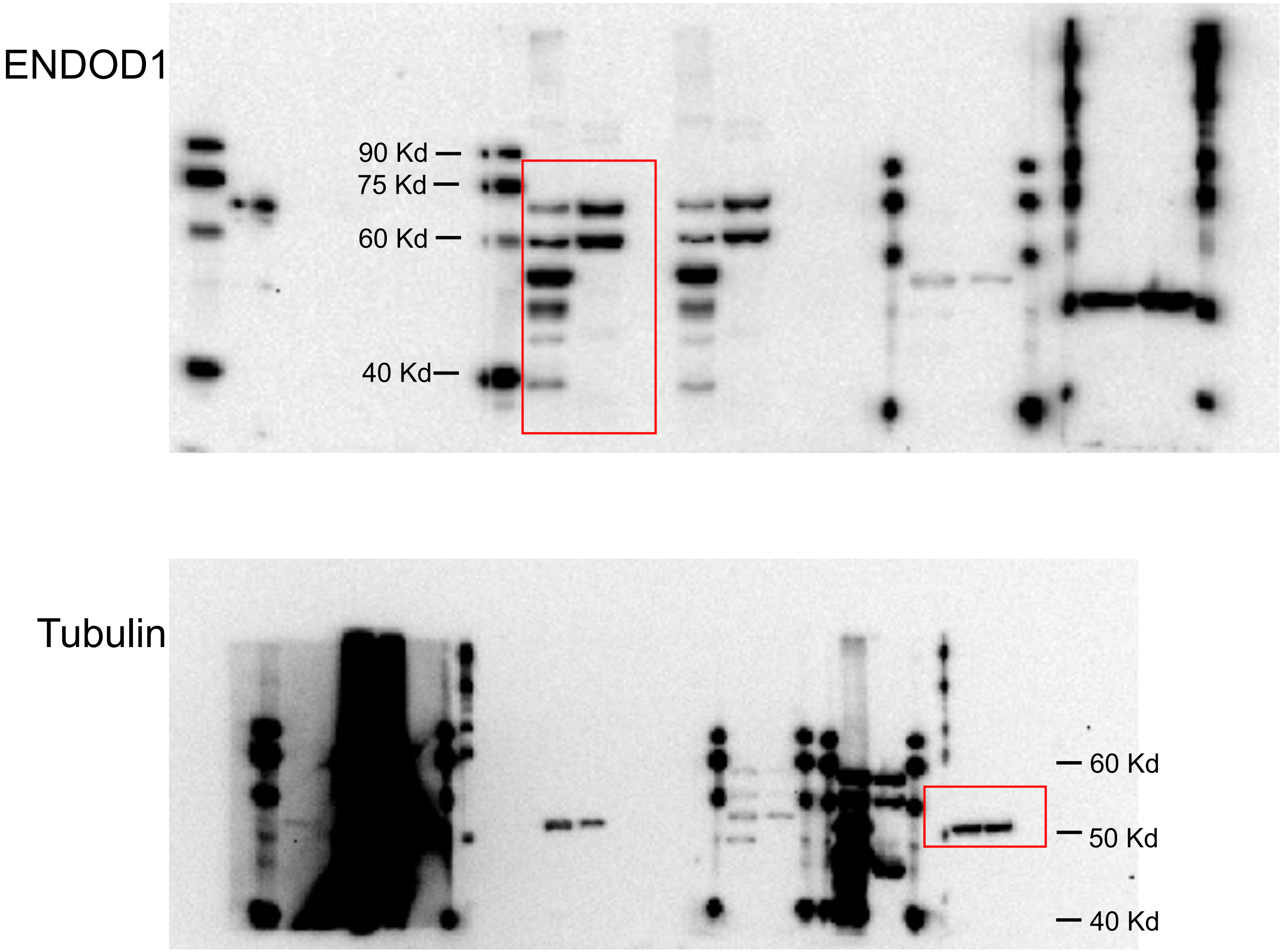

Fig S4b

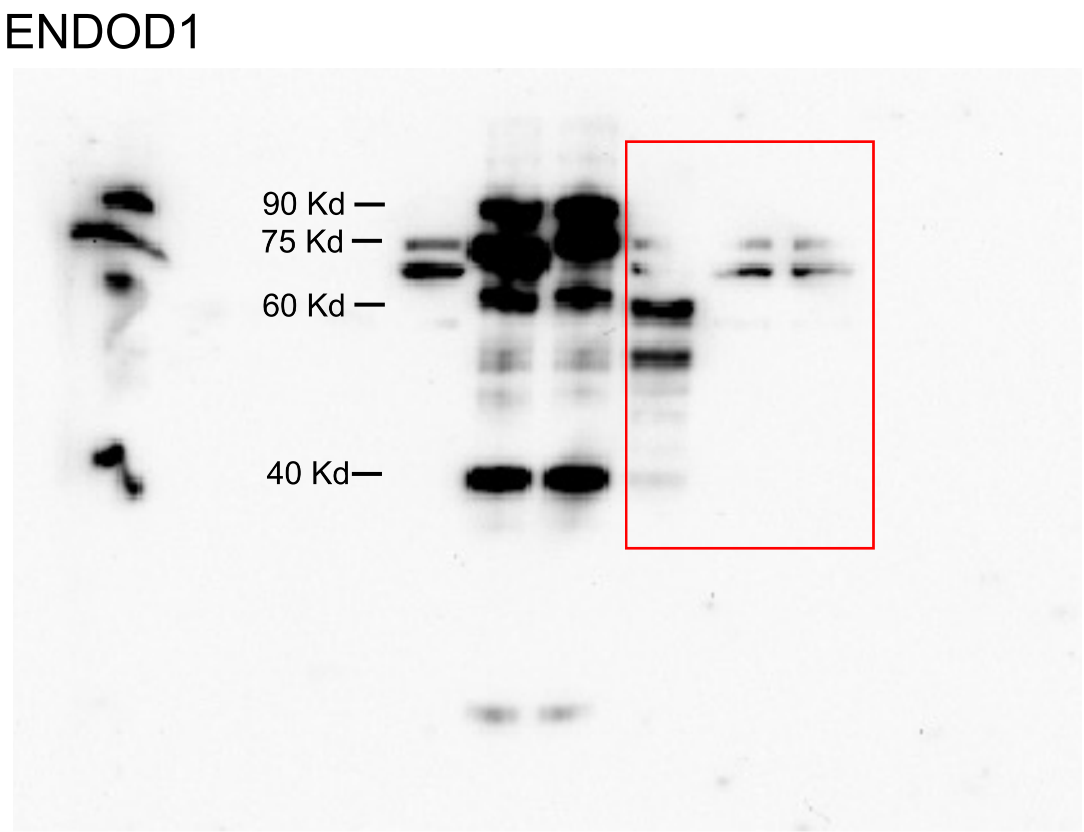


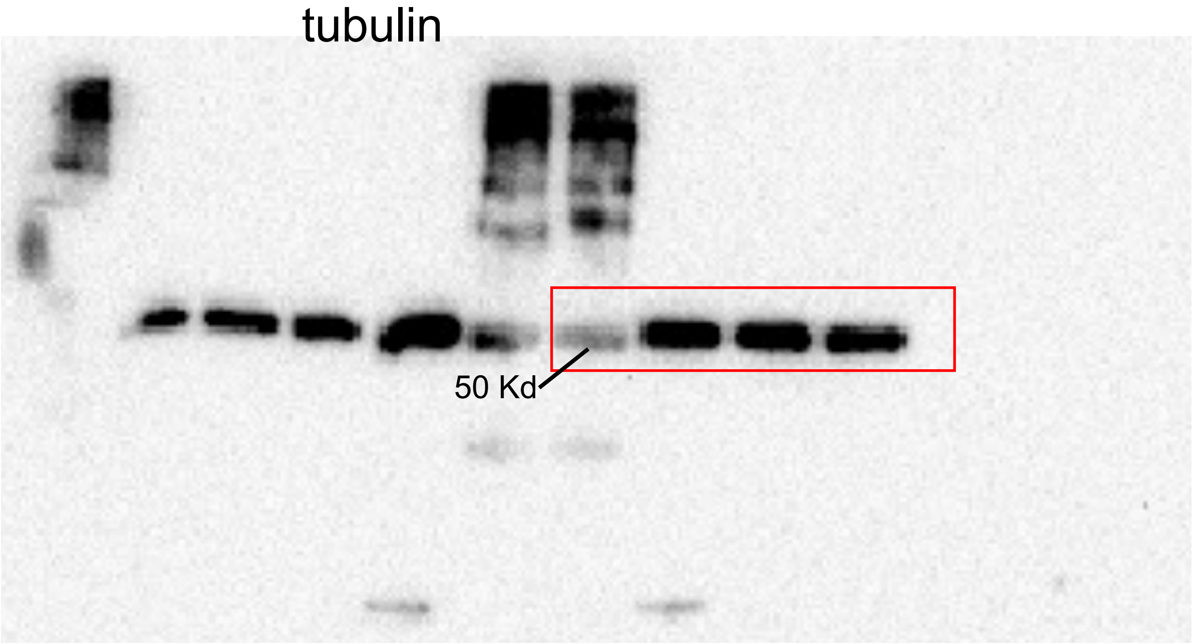

Fig S4f

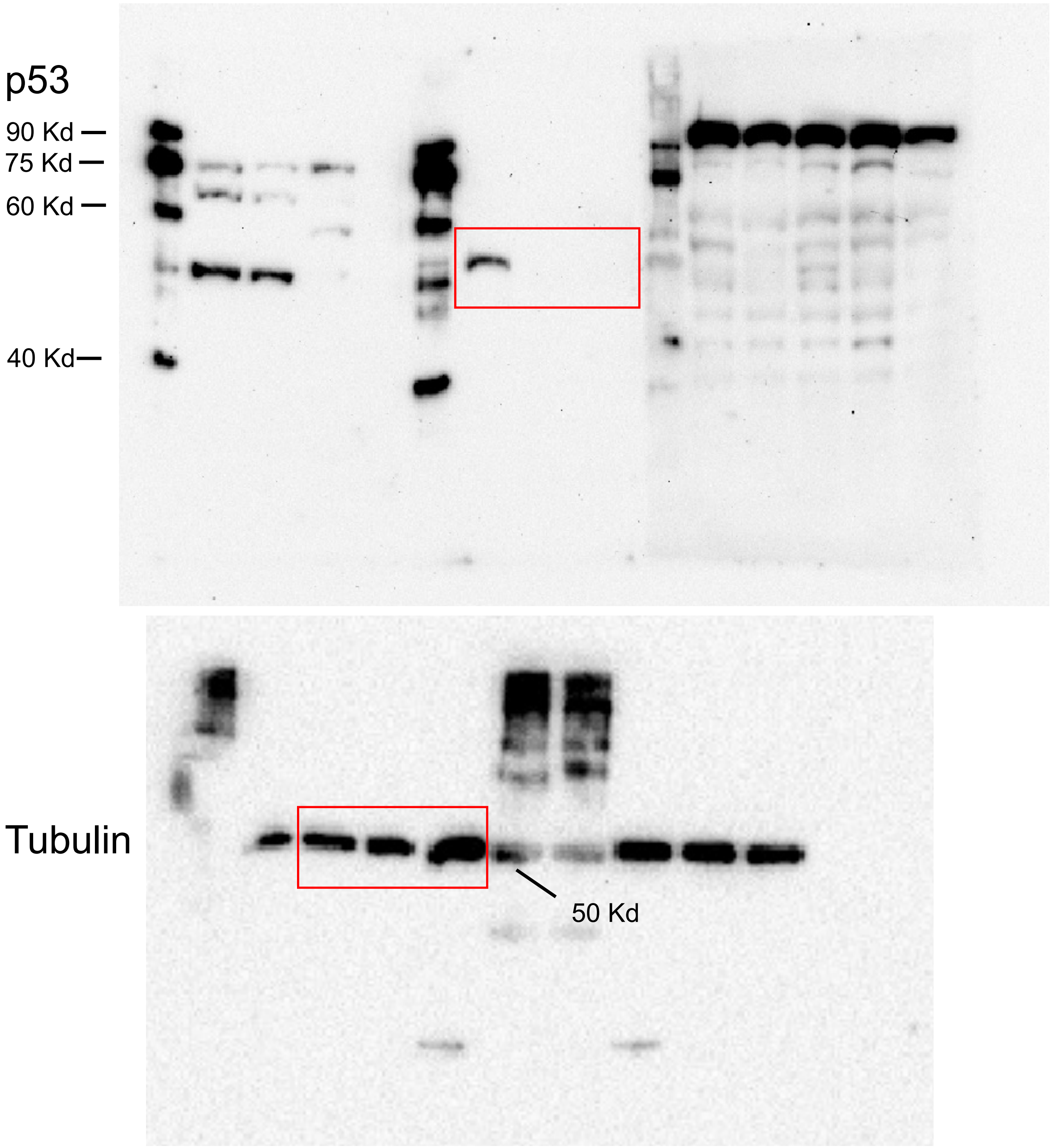

Fig S4f

ENDOD1

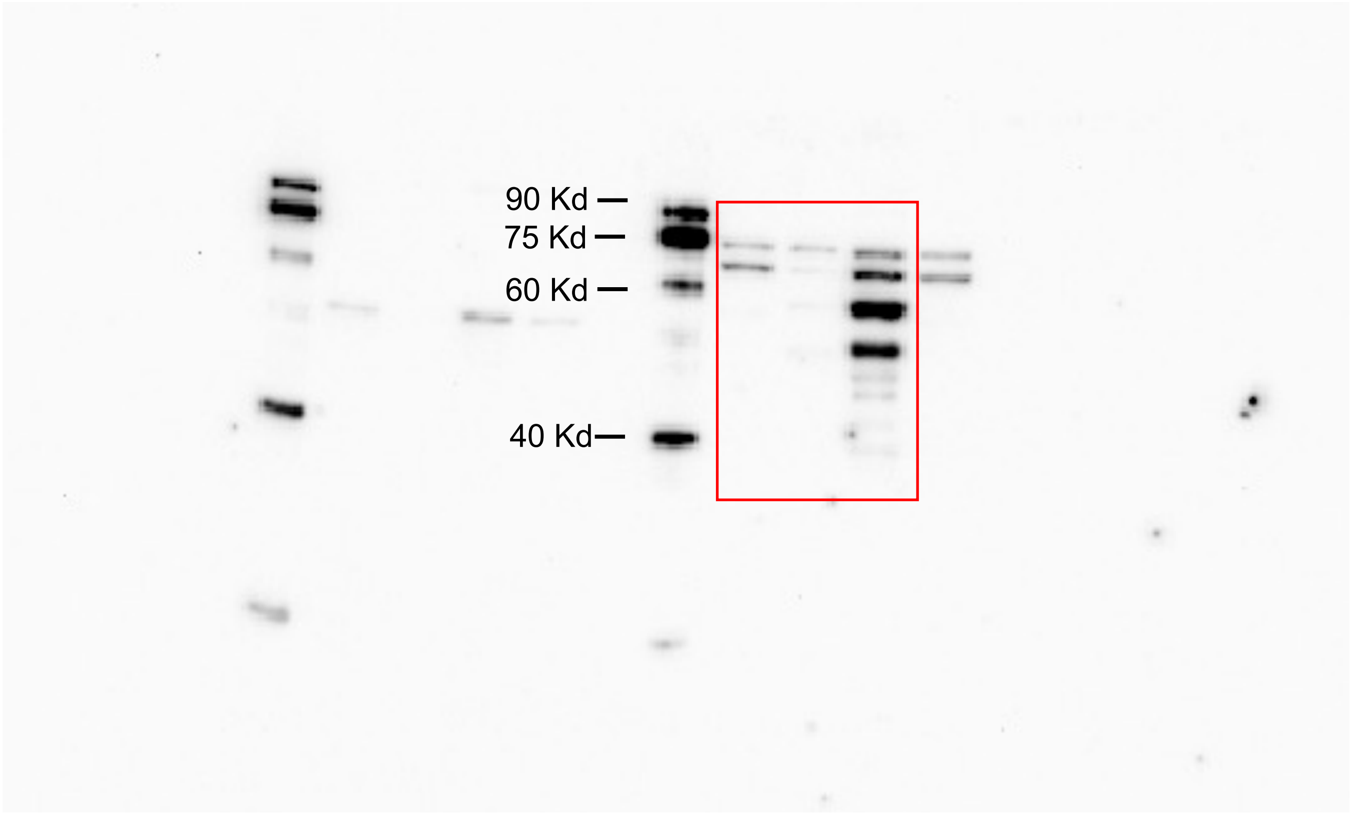

Fig S4h

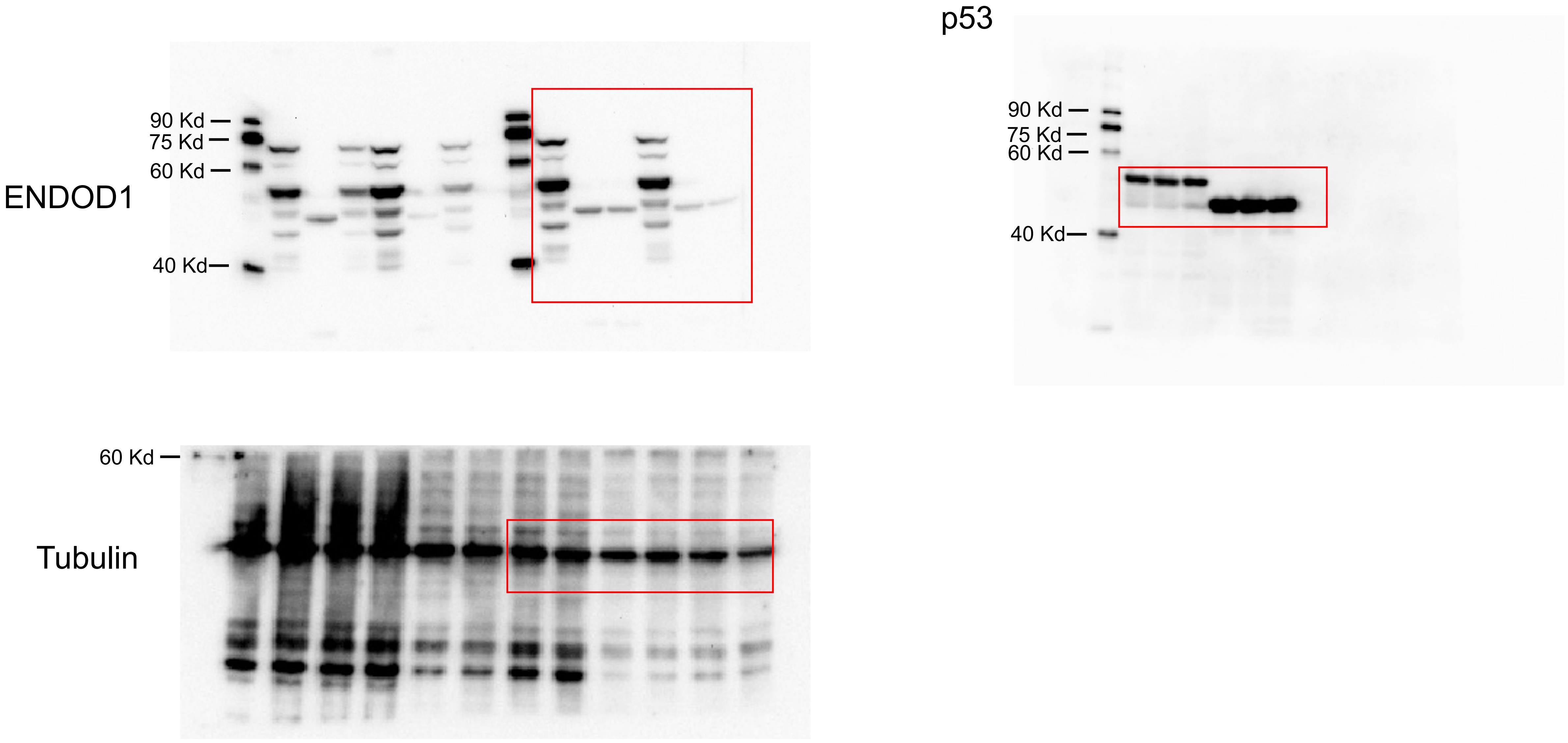
ENDOD1
90 Kd
75 Kd
60 Kd
40 Kd
Tubulin
60 Kd
p53
90 Kd
75 Kd
60 Kd
40 Kd

Fig S5e

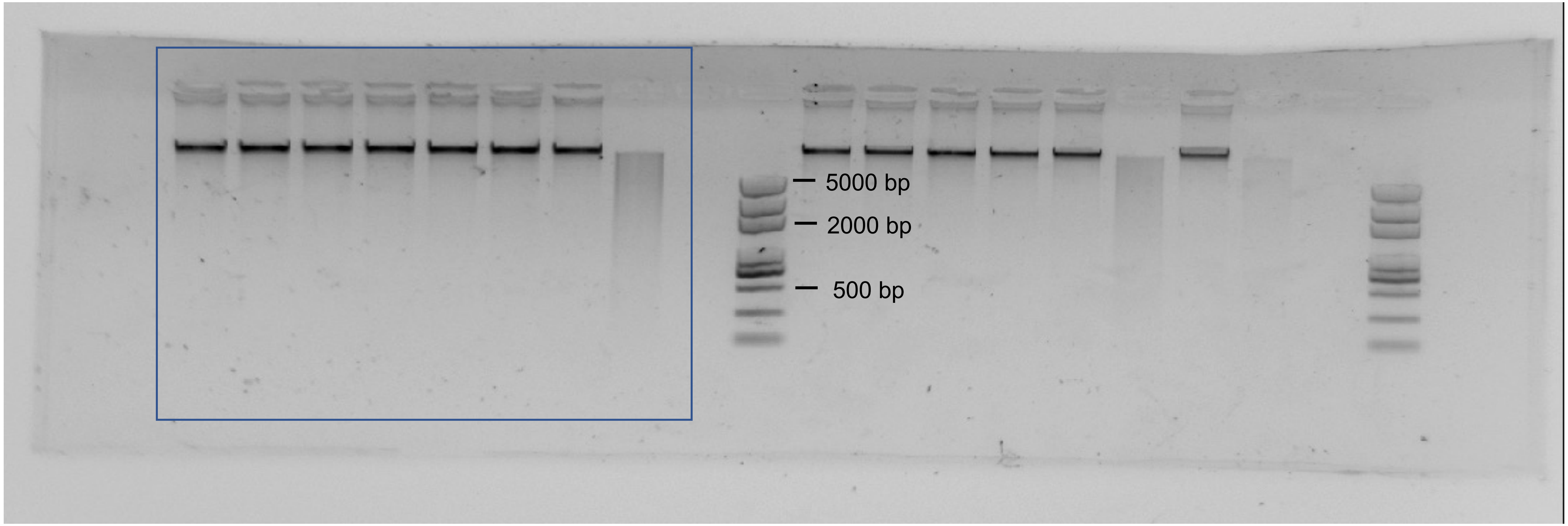

Fig S5h

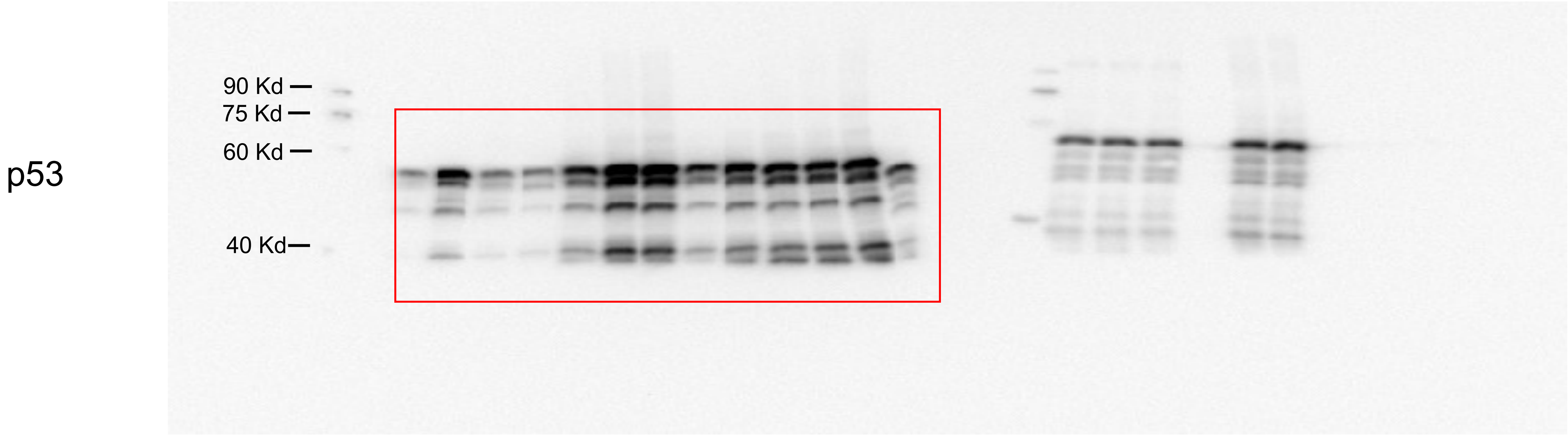


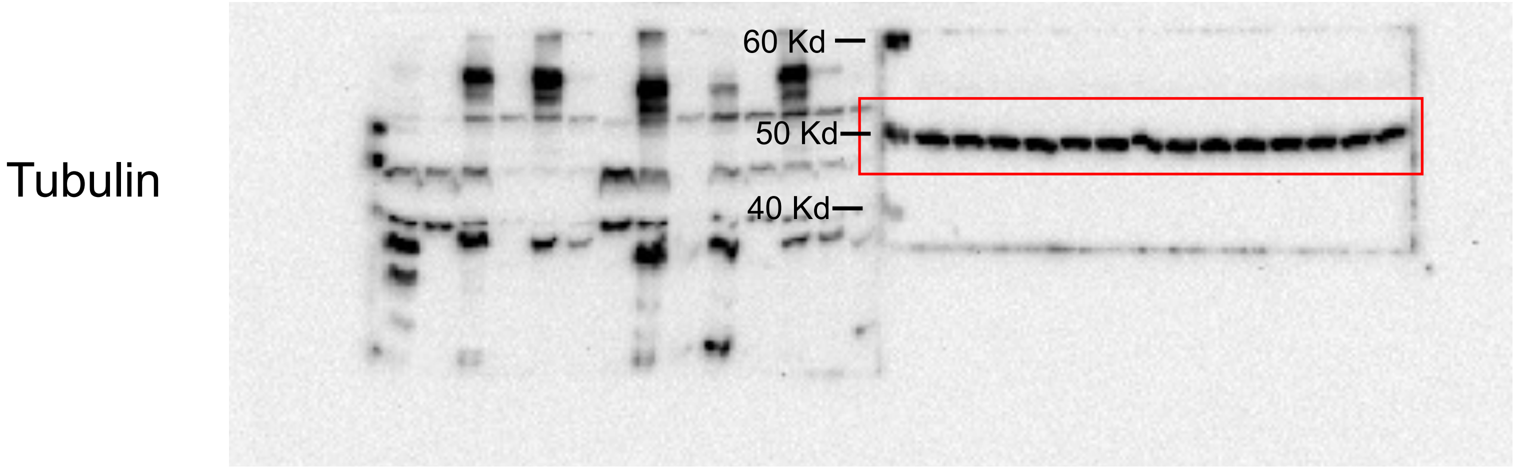

Fig S6h

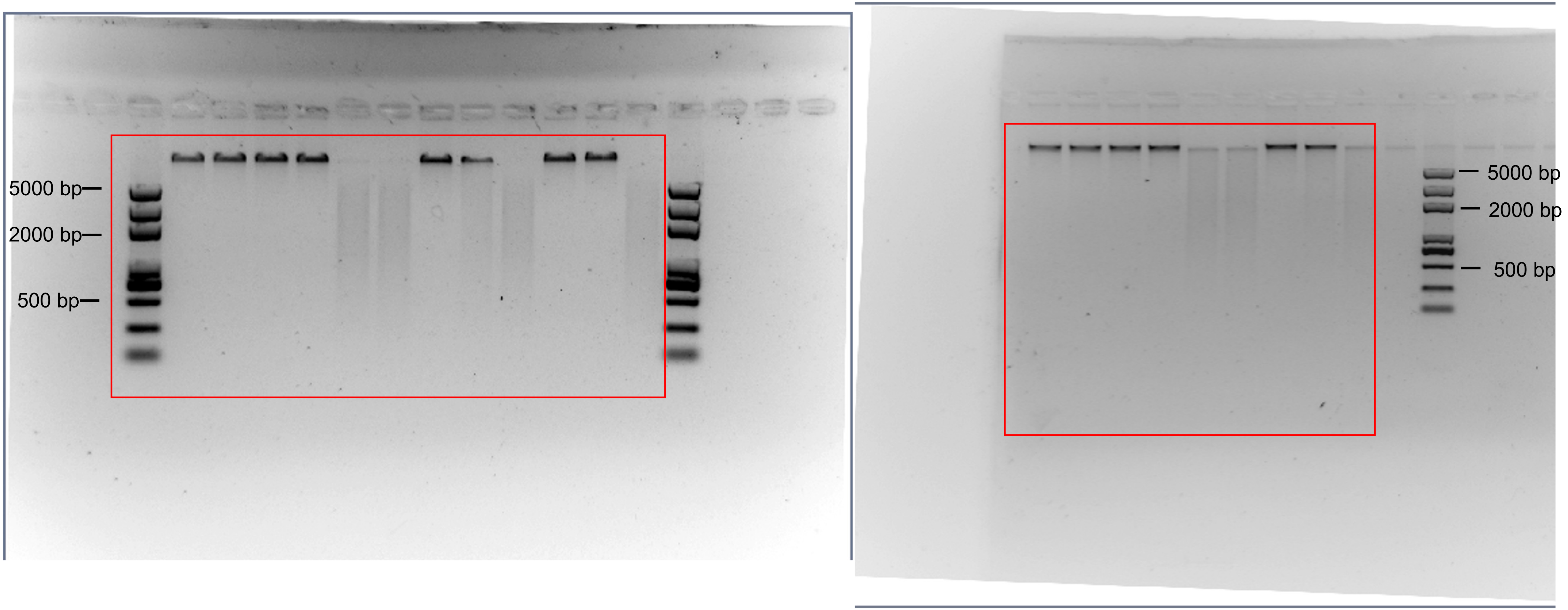

Fig S8a

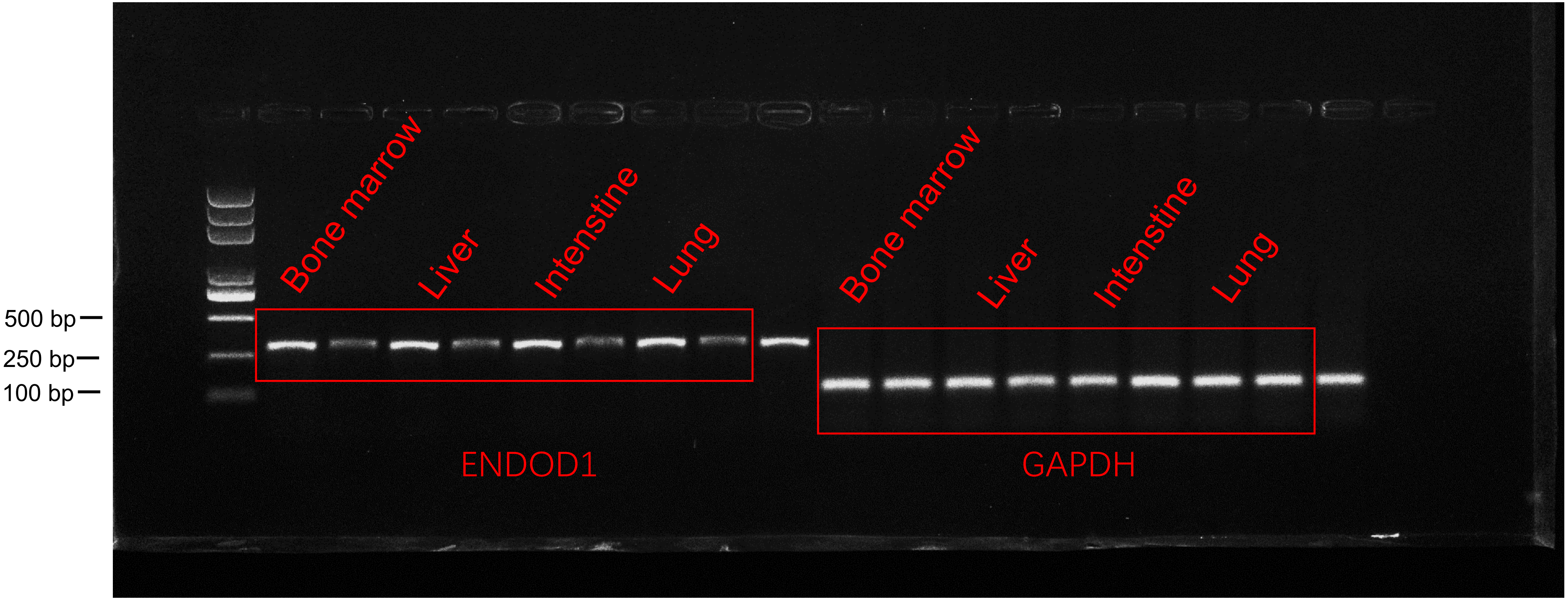
500 bp
250 bp
100 bp
Bone marrow
Liver
Intenstine
Lung
Bone marrow
Liver
Intenstine
Lung
ENDOD1
GAPDH

Fig S8a

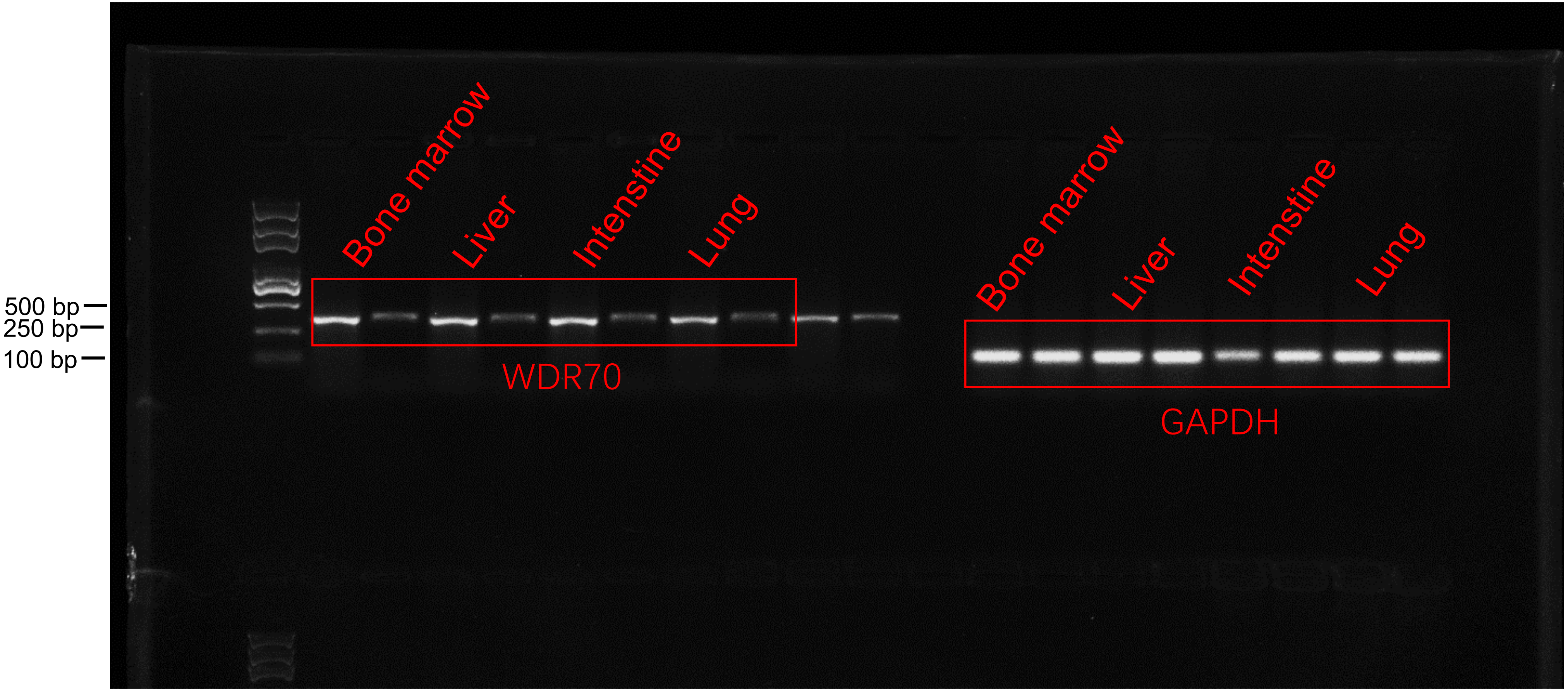
500 bp
250 bp
100 bp
Bone marrow
Liver
Intenstine
Lung
WDR70
Bone marrow
Liver
Intenstine
Lung
GAPDH

Fig S8a

heart

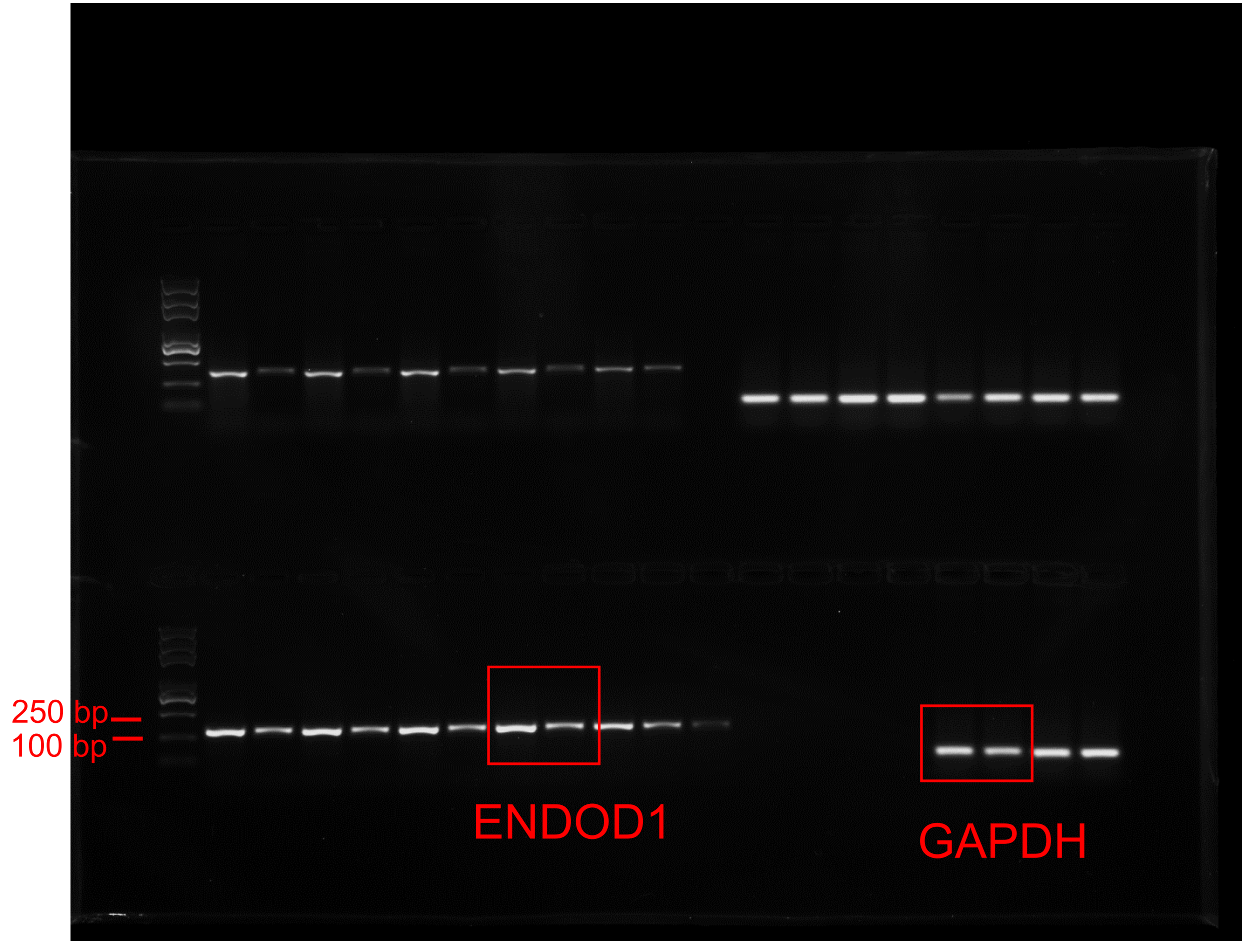


heart

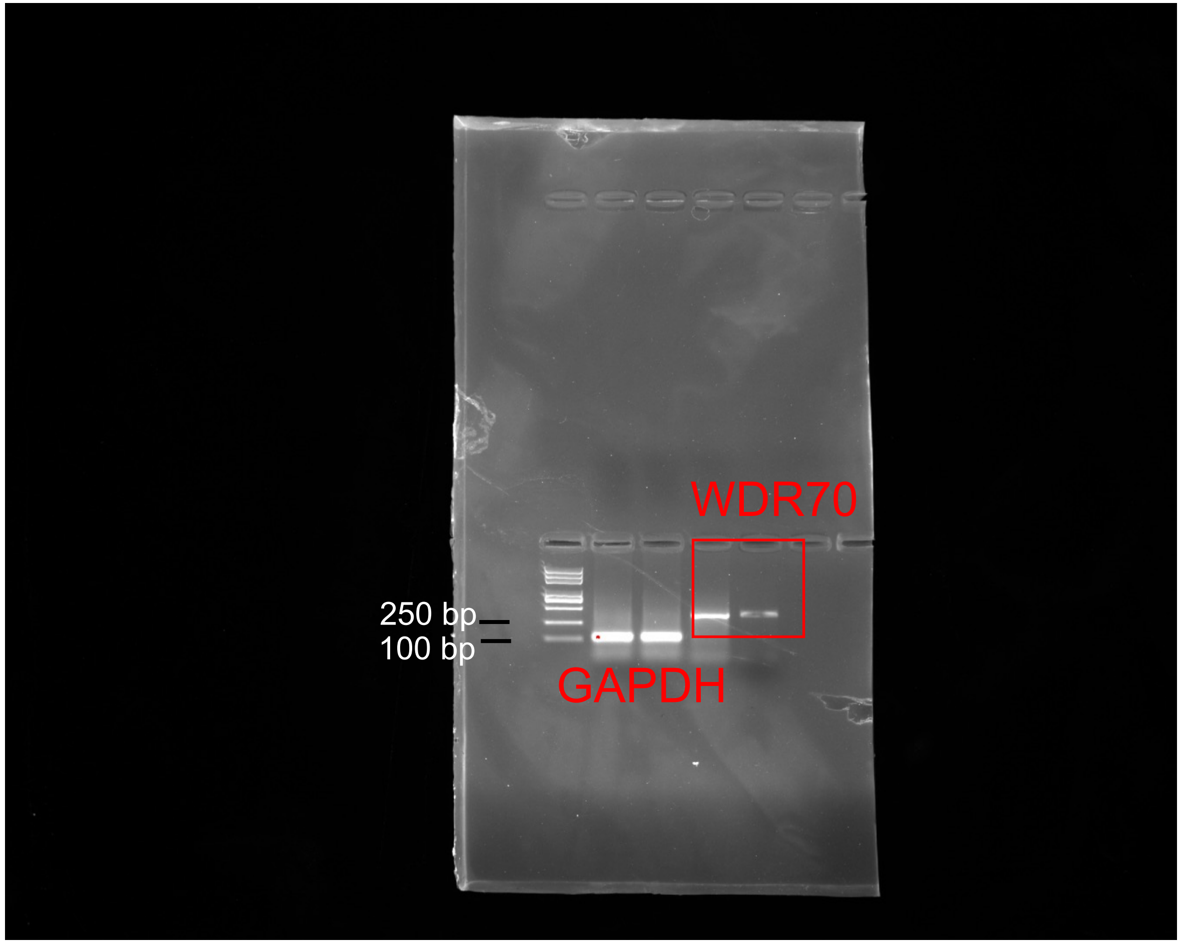

Fig S10 a

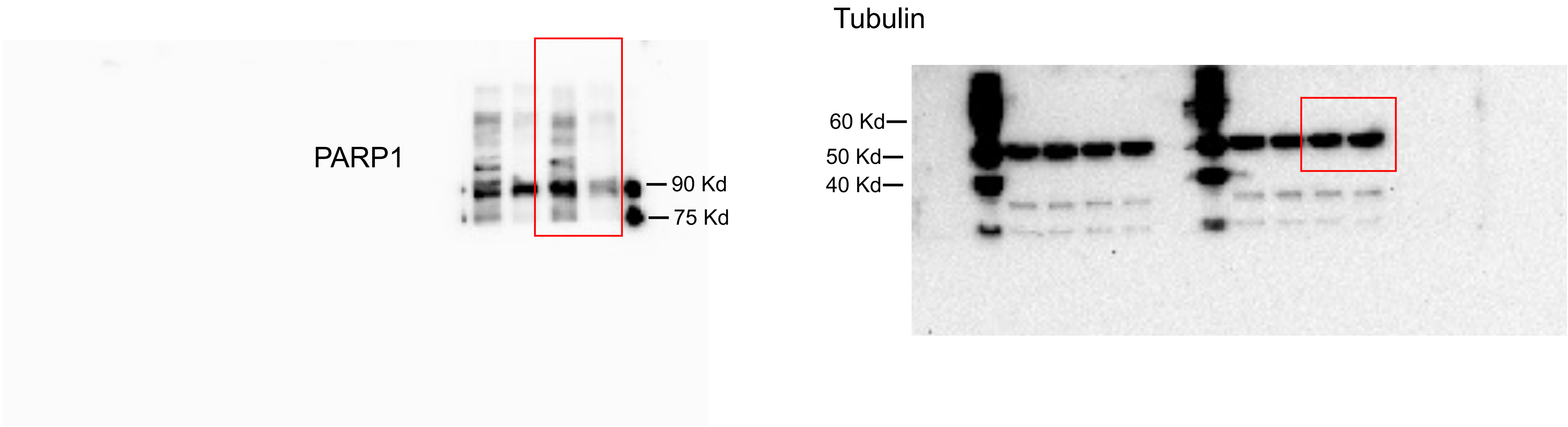

Fig S10a

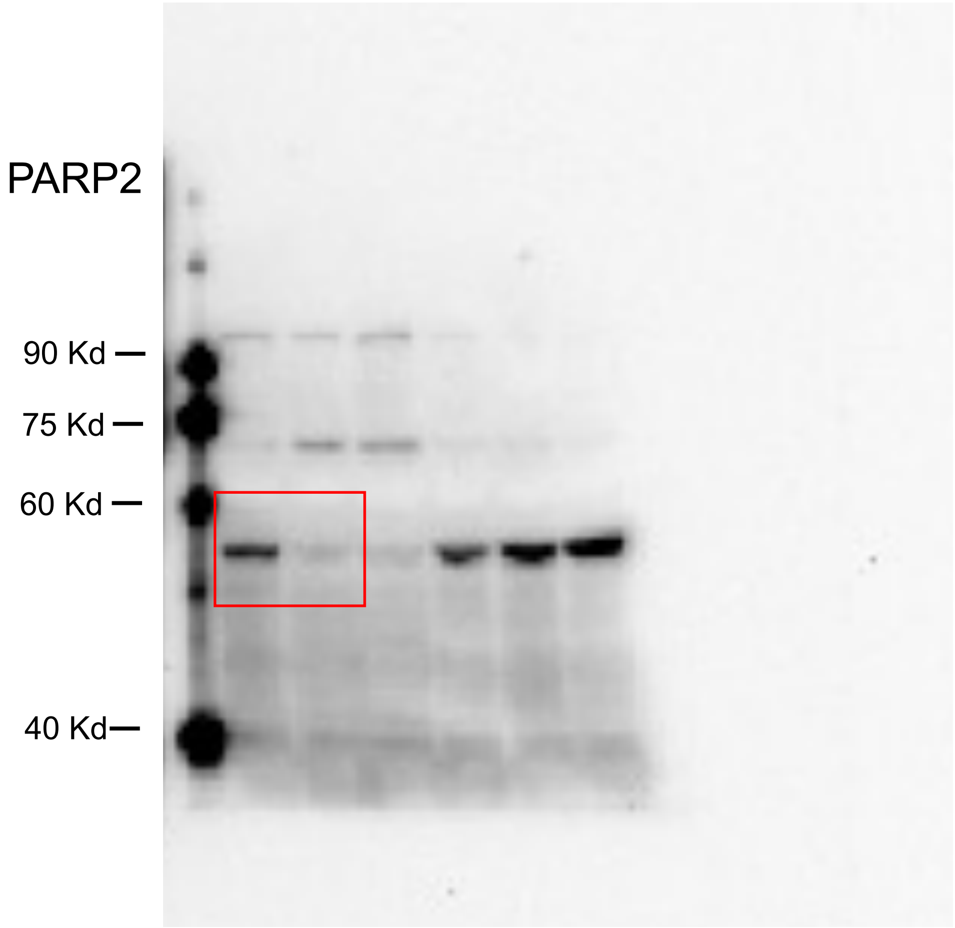


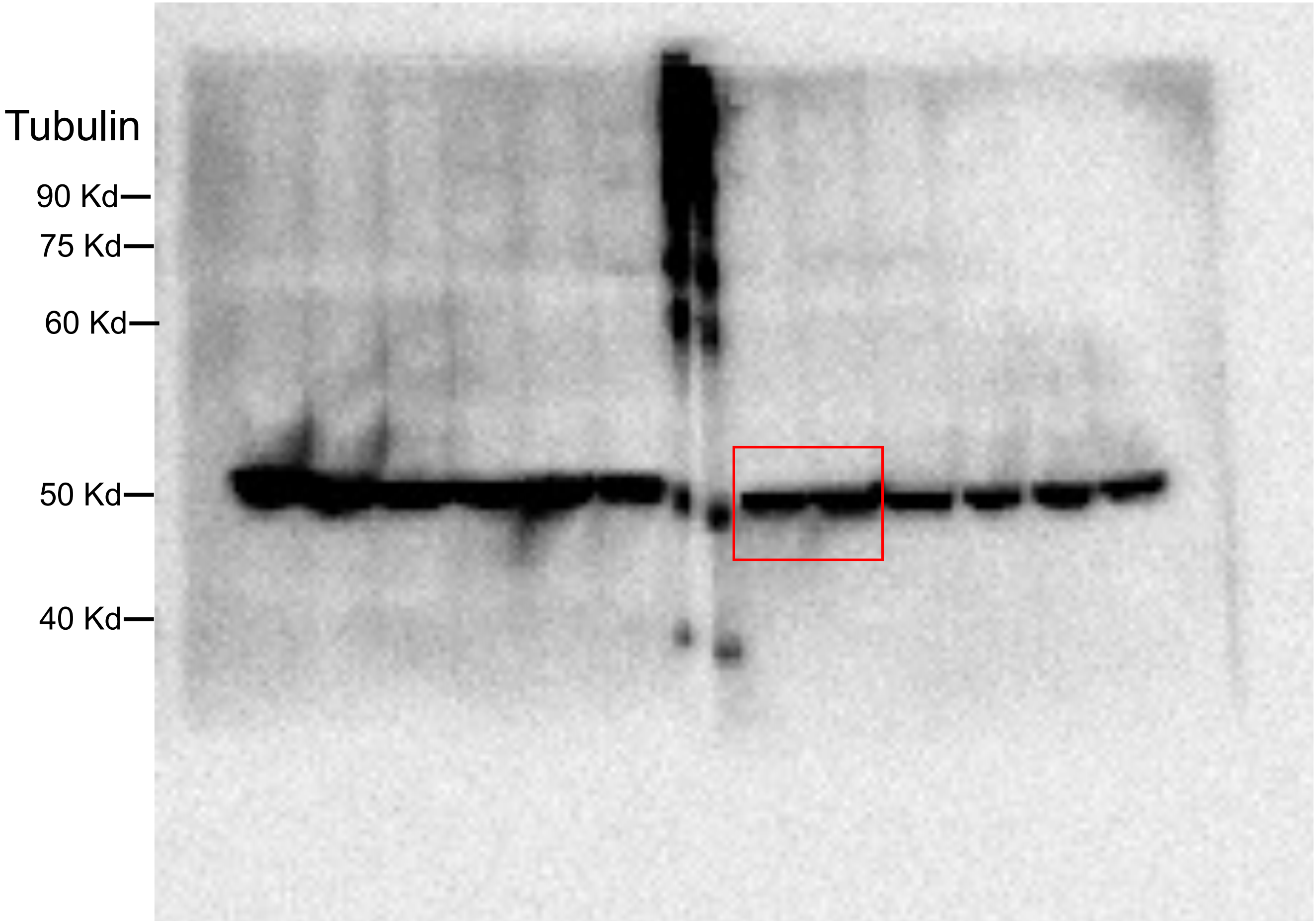

Fig S10a

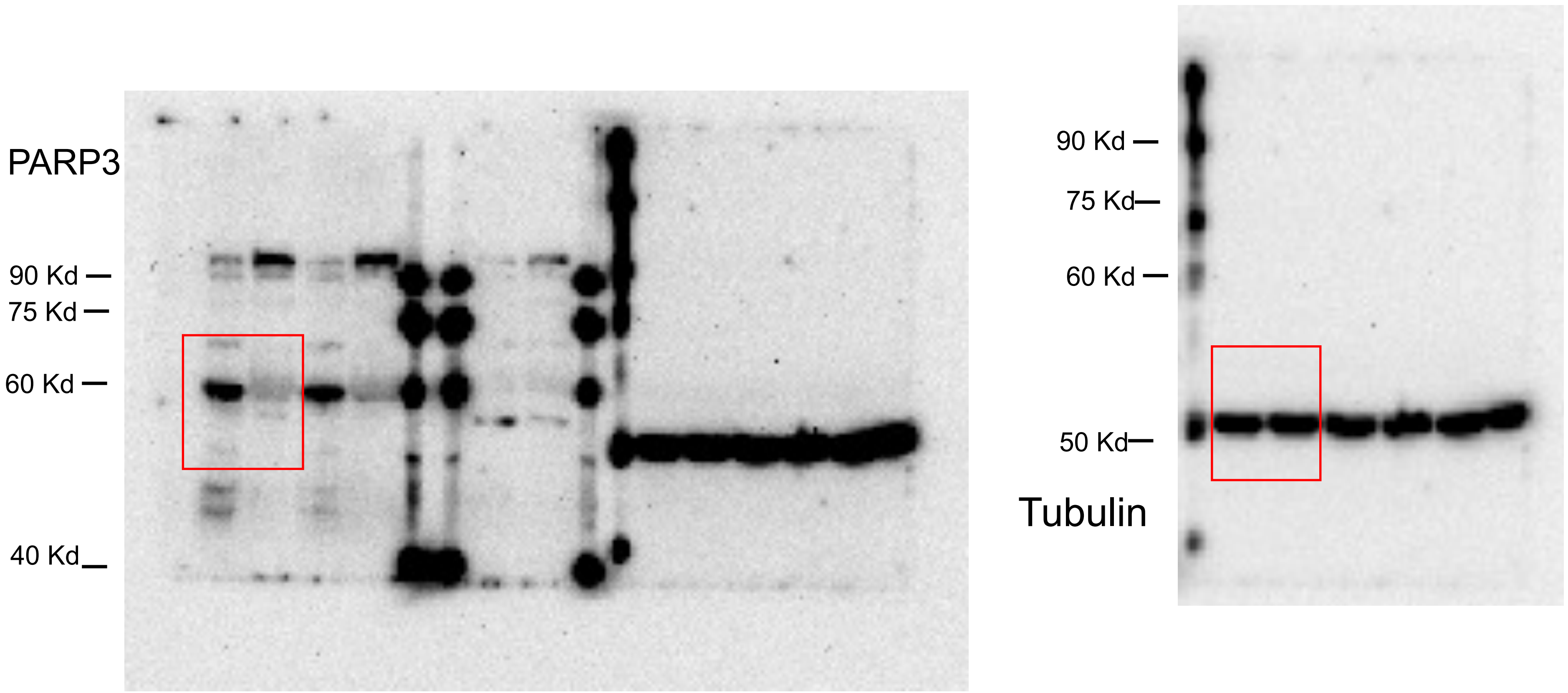

Fig S10 b

p21

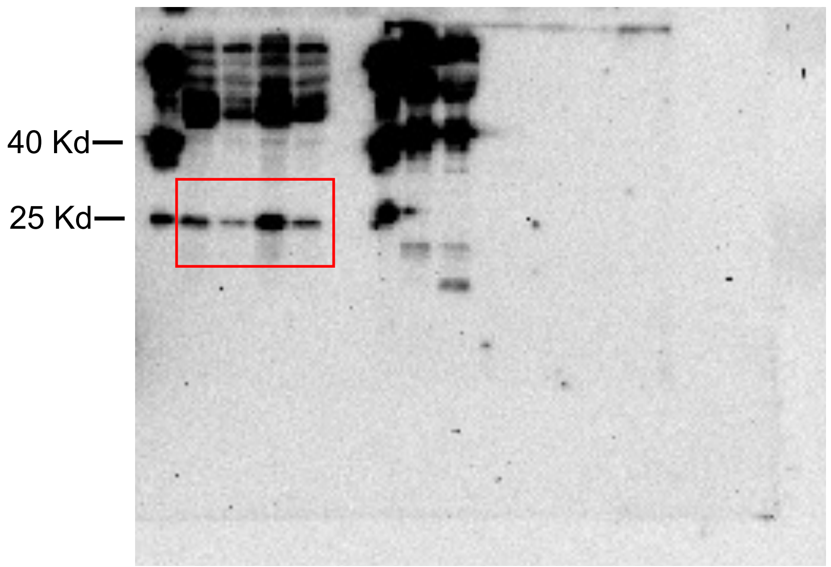


Tubulin

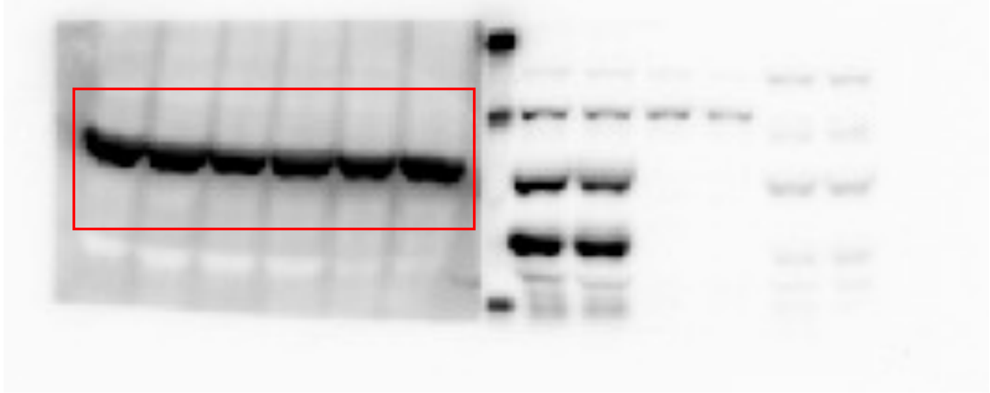

The size estimated by Precision Plus Protein Standards(Bio-Rad, 161-0374) of 50 Kd.

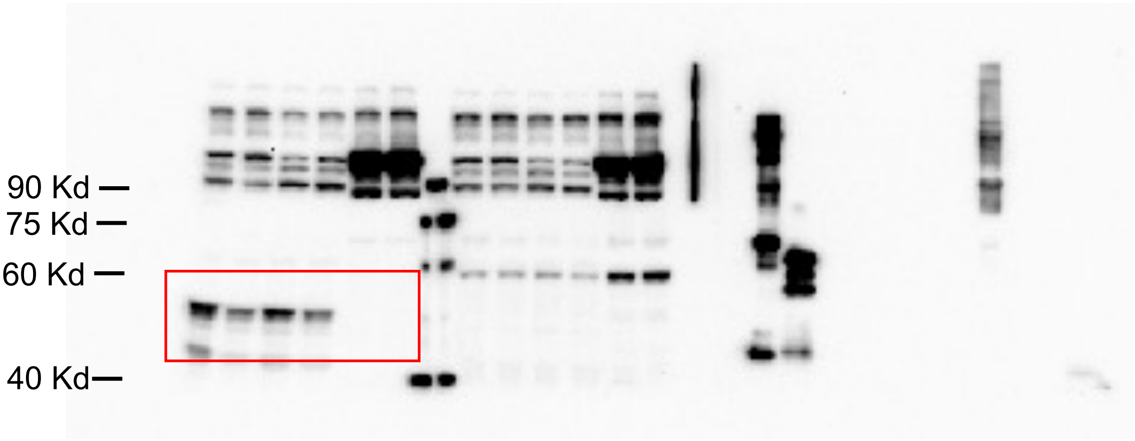


p53

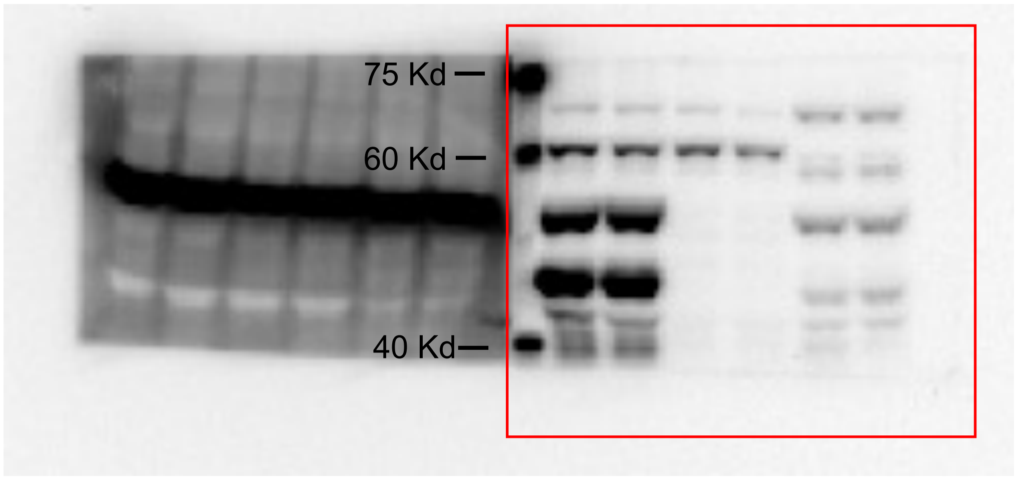


ENDOD1

Fig S10c

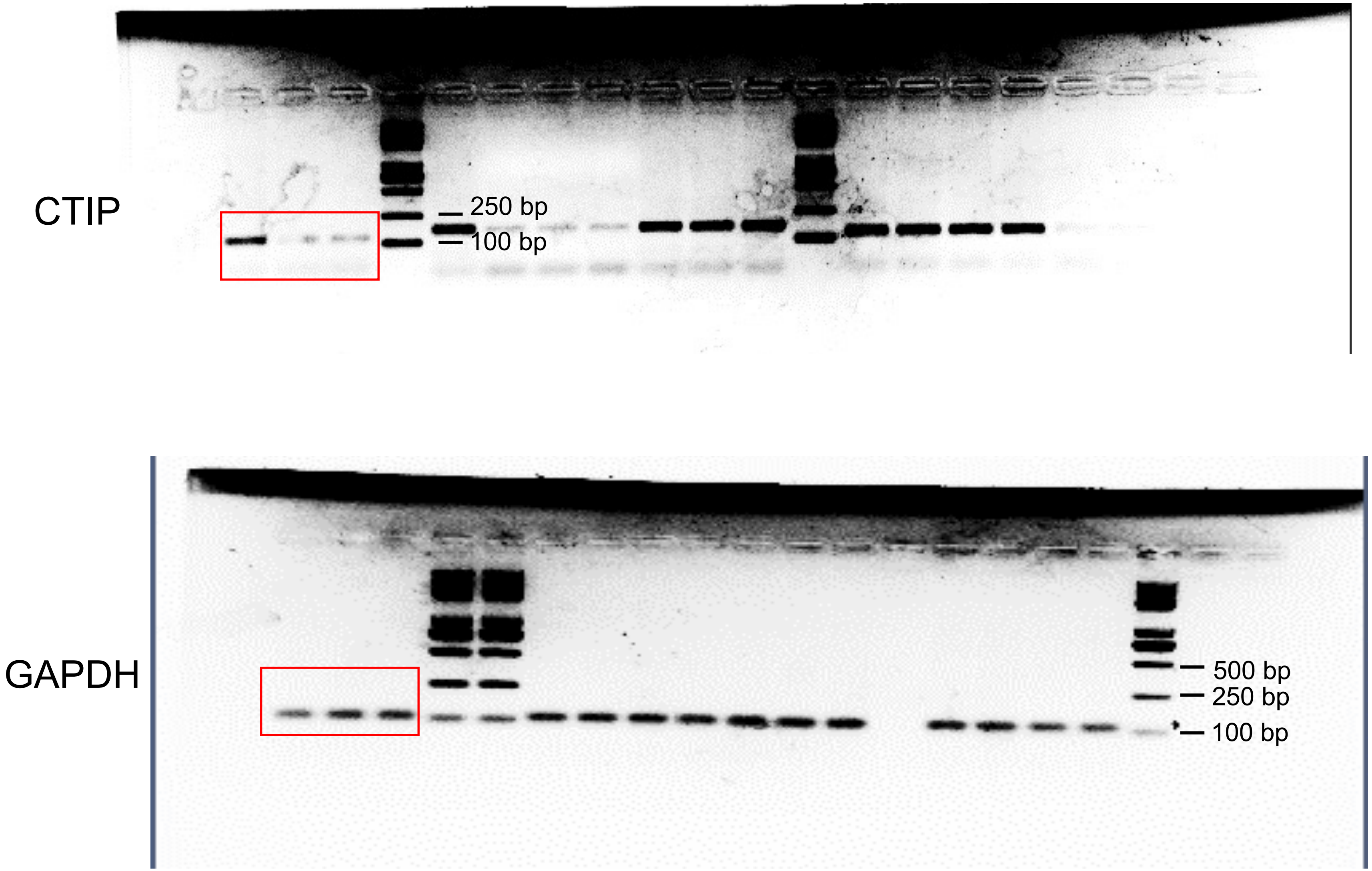

Fig S10c

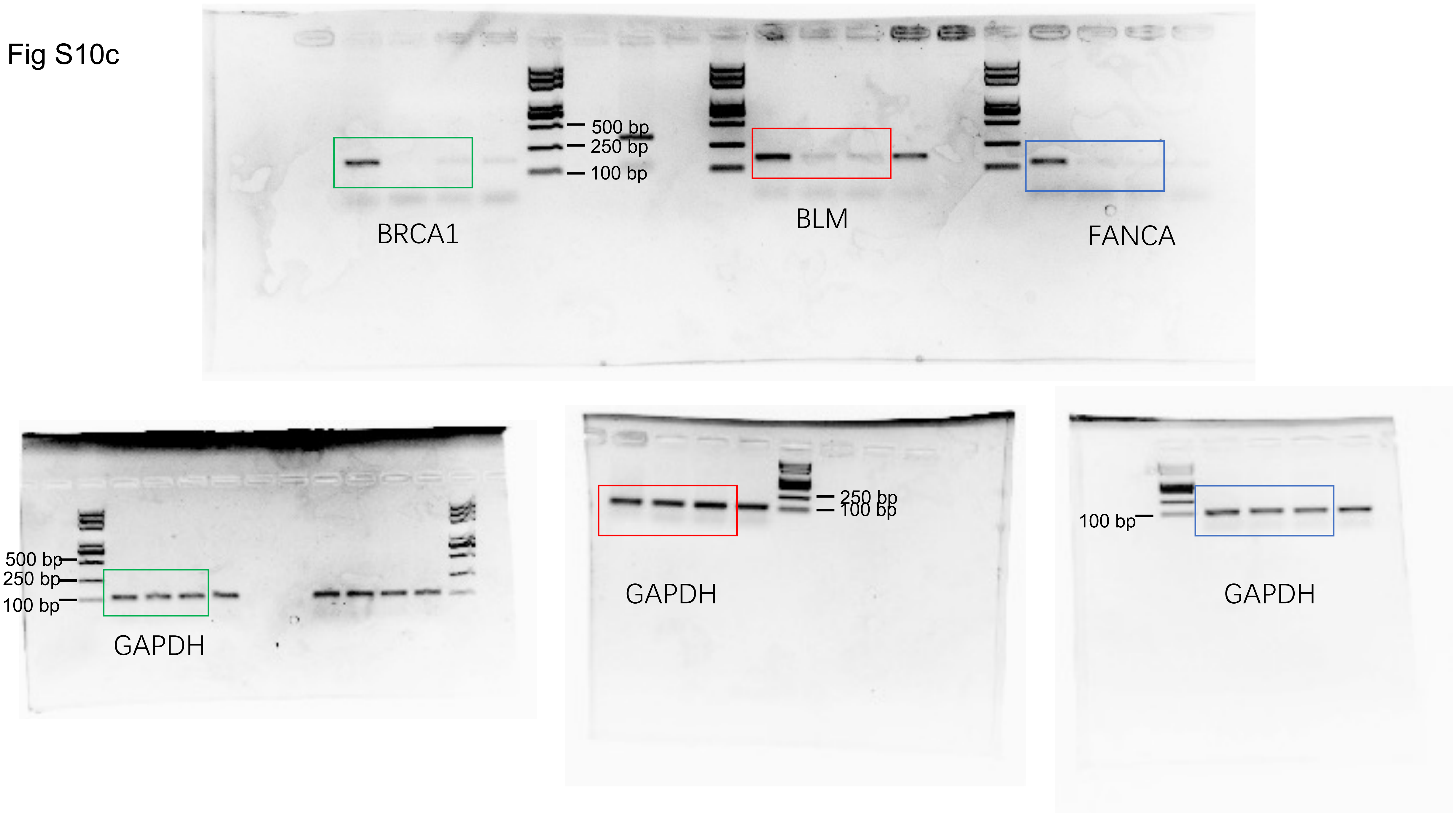
500 bp
250 bp
100 bp
BRCA1
BLM
FANCA
500 bp
250 bp
100 bp
GAPDH
250 bp
100 bp
GAPDH
100 bp
GAPDH

Fig S10c

BRCA2

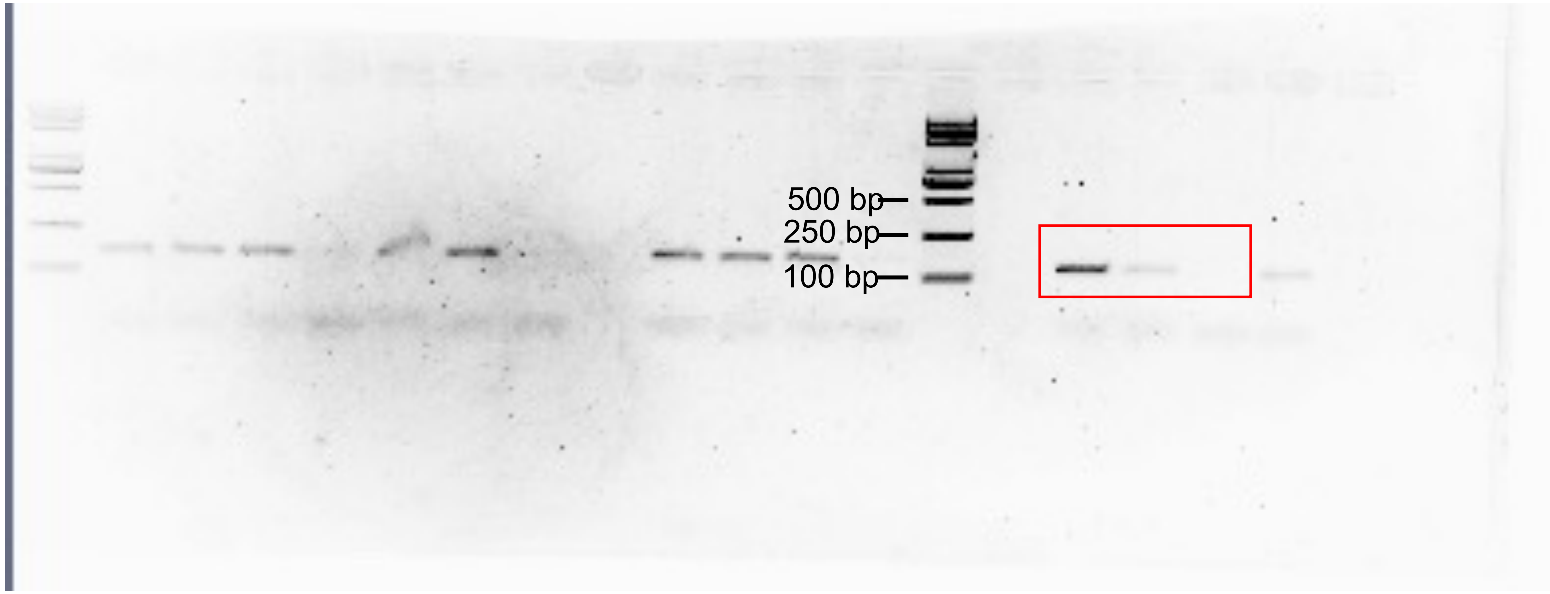


GAPDH

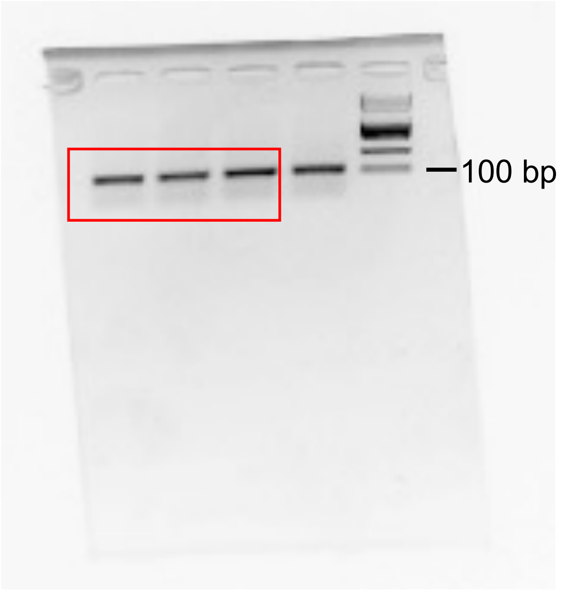

Fig S10c

CHK1

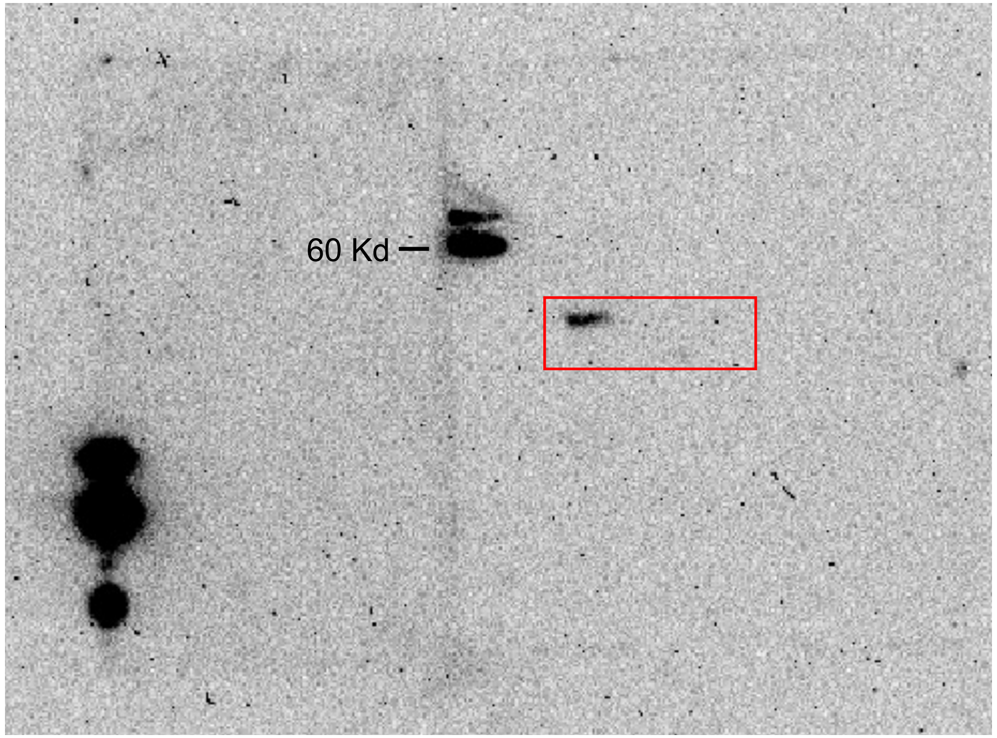


Tubulin

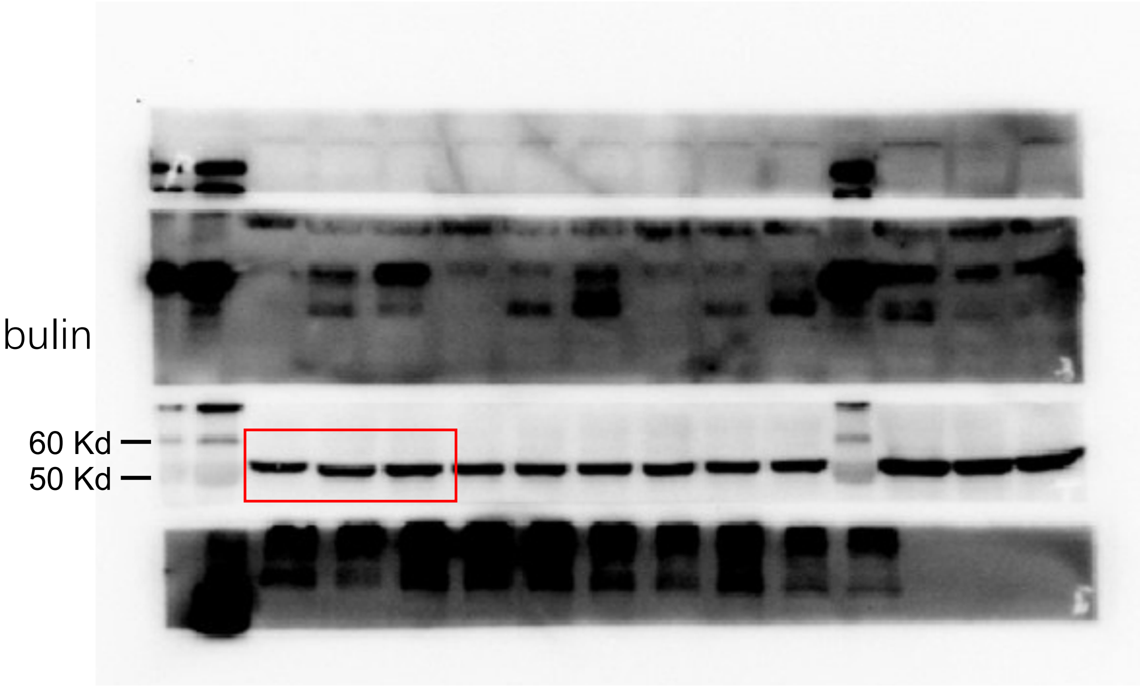

Fig S10c

BRCA2

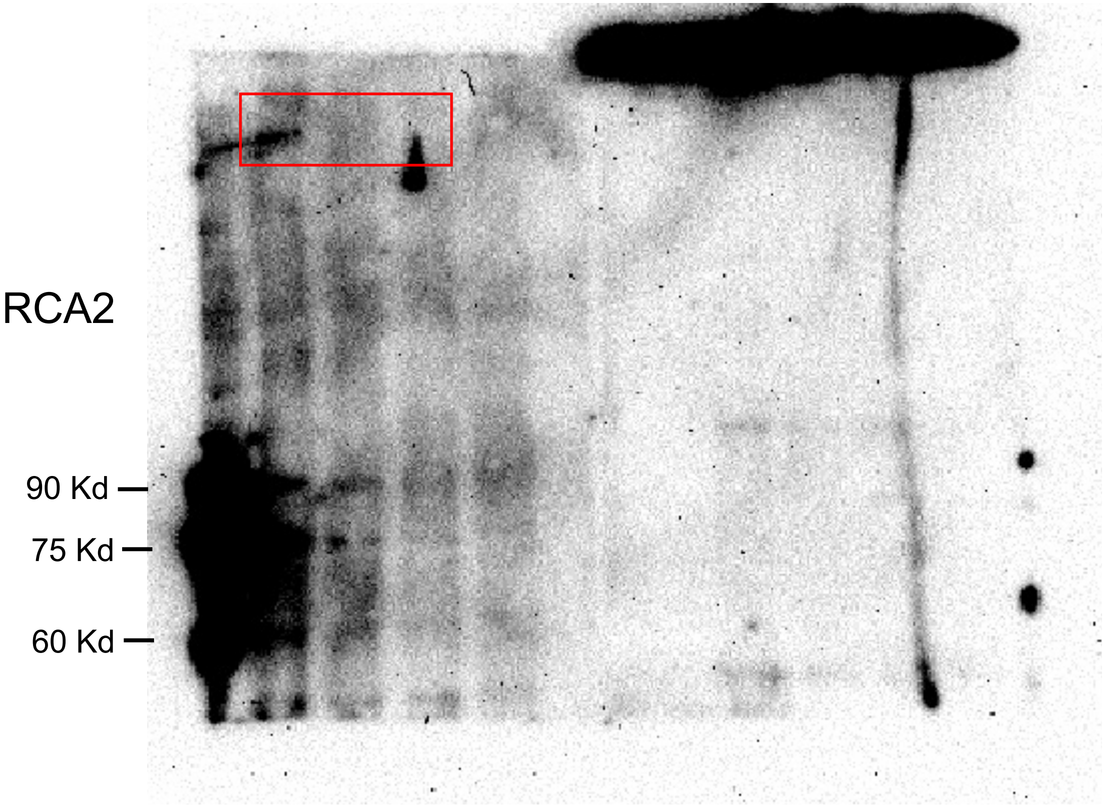


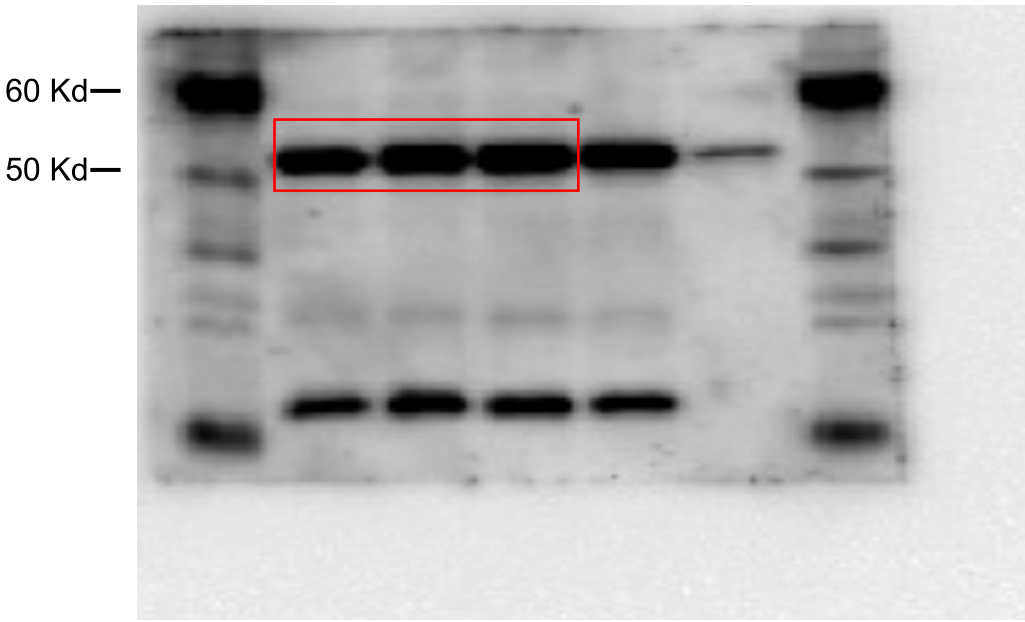


Tubulin

Fig S10c

ARID1A

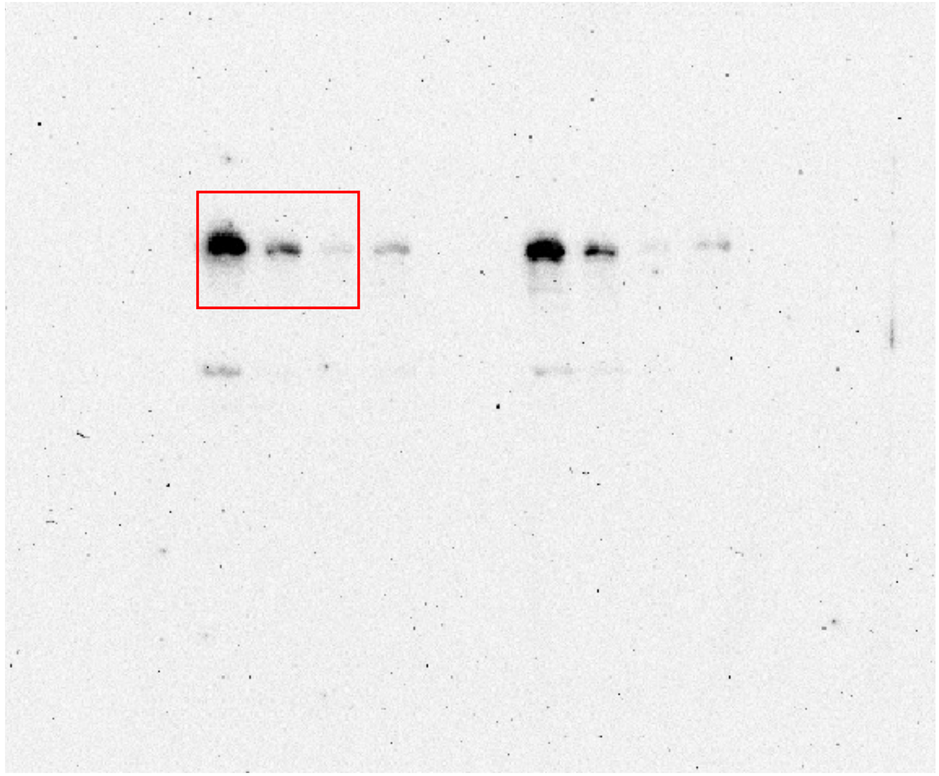

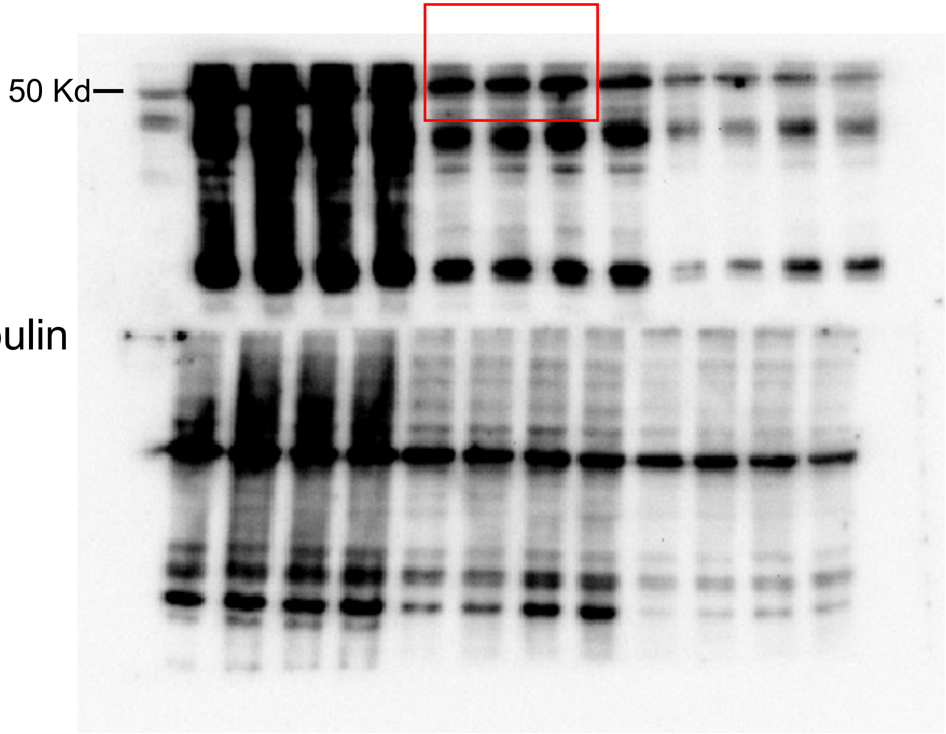

Fig S10c

MRE11

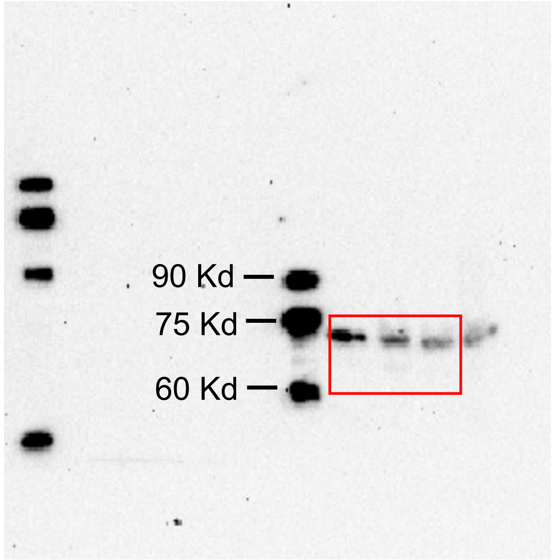


Tubulin

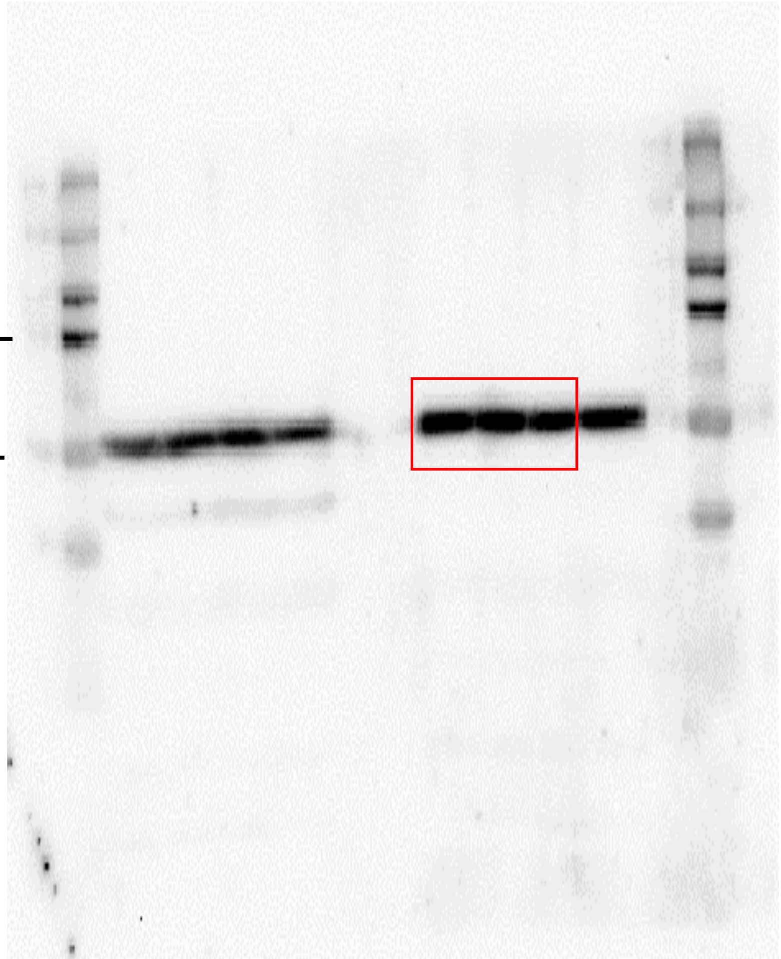

Fig S10d

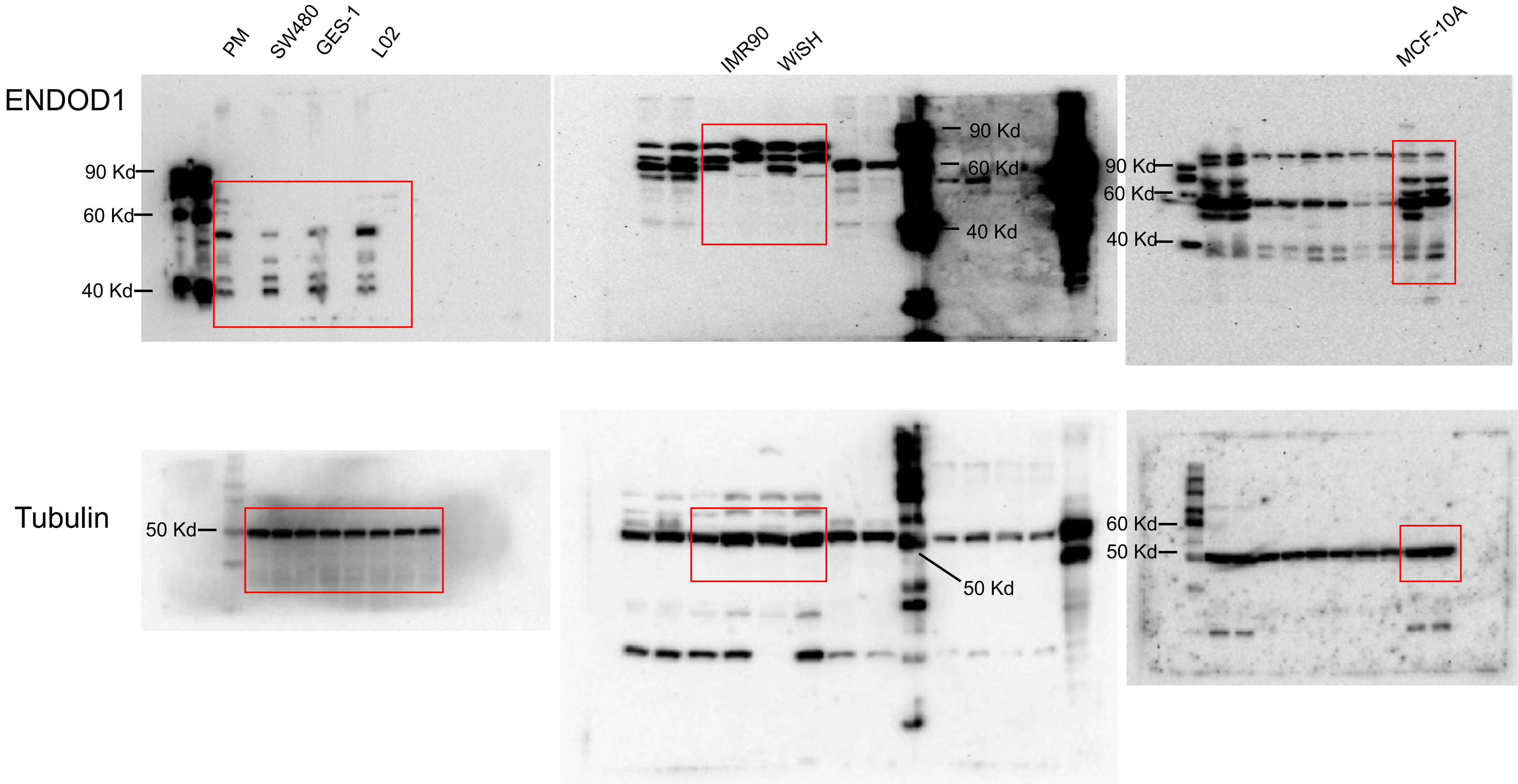

PM
SW480
GES-1
L02
IMR90
WiSH
MCF-10A
ENDOD1
90 Kd
60 Kd
40 Kd
90 Kd
60 Kd
40 Kd
90 Kd
60 Kd
40 Kd
Tubulin
50 Kd
50 Kd
60 Kd
50 Kd

Fig S10d

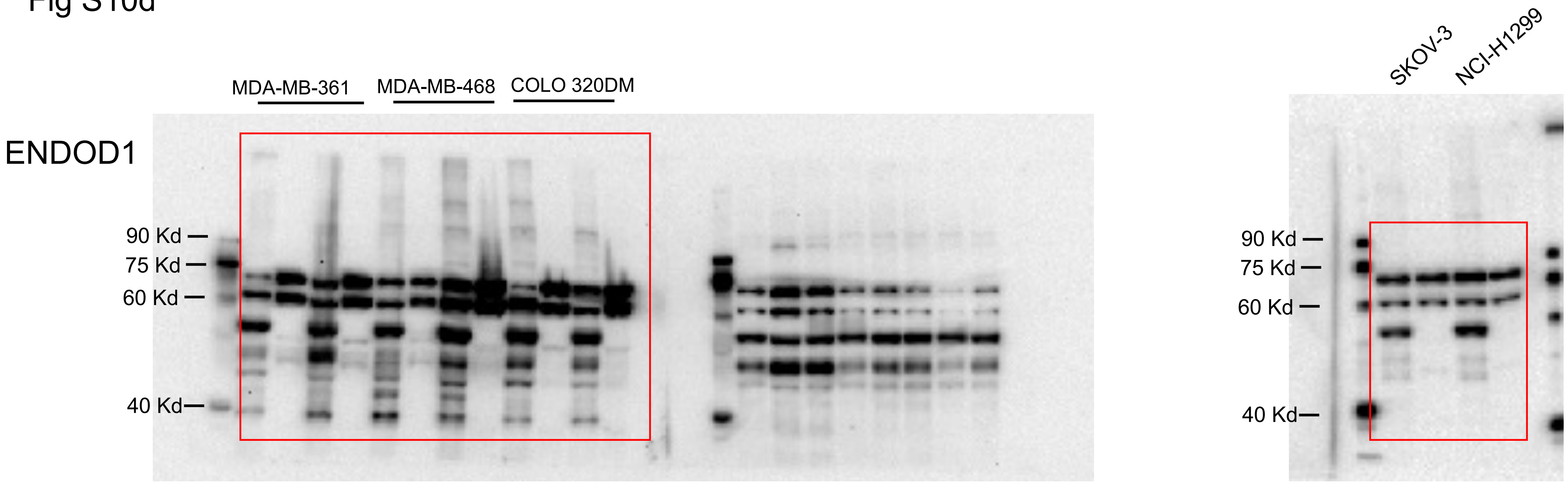
ENDOD1
MDA-MB-361
MDA-MB-468
COLO 320DM
90 Kd
75 Kd
60 Kd
40 Kd
SKOV-3
NCI-H1299
90 Kd
75 Kd
60 Kd
40 Kd

### Tubulin

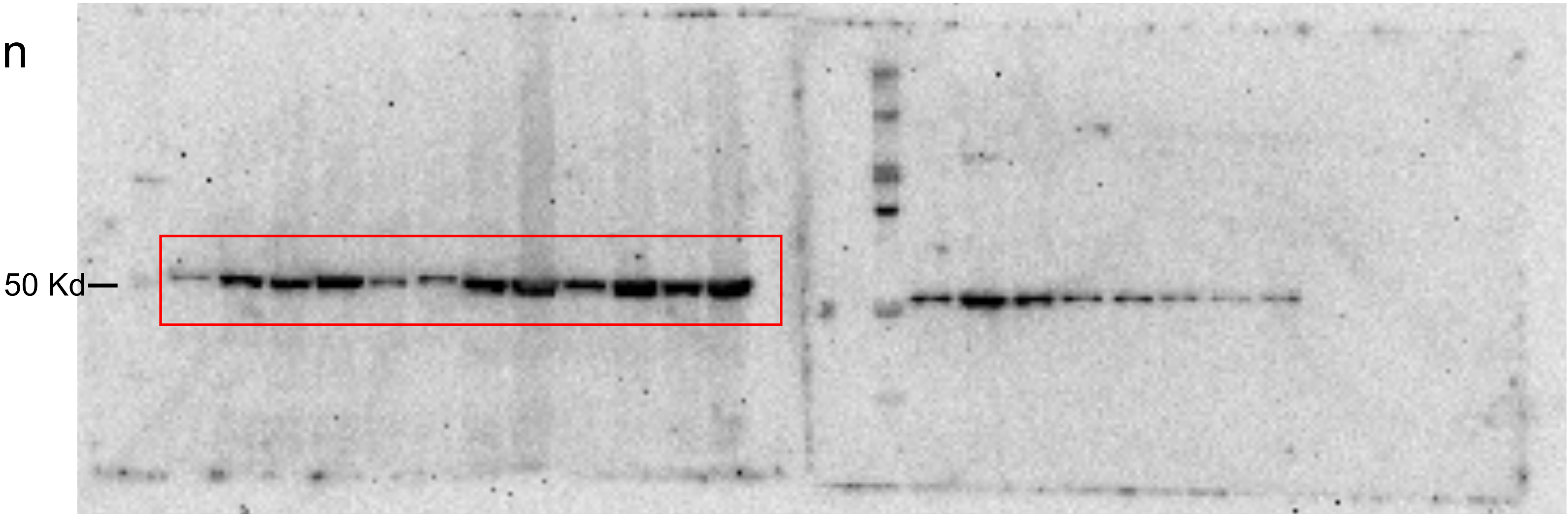


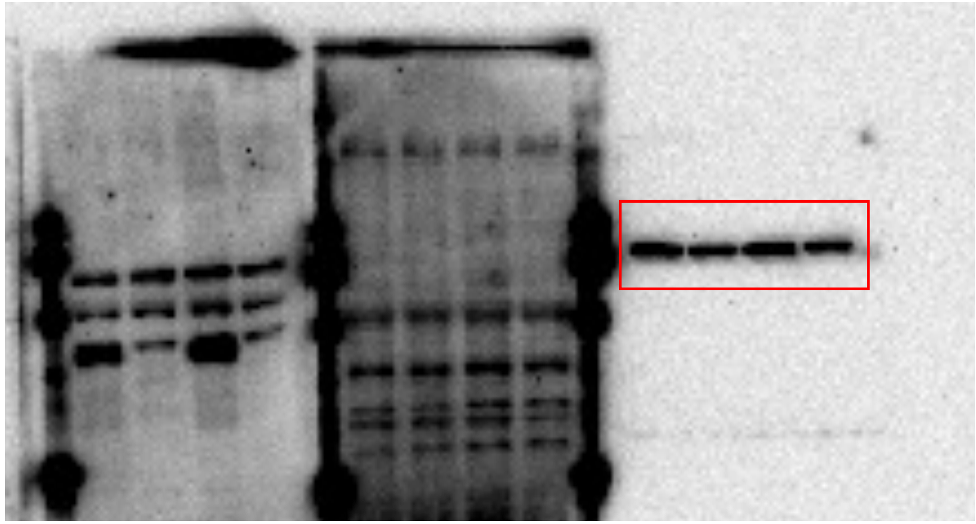

In right blot, protein size was estimated by prestained Precision Plus Protein Standards(Bio-Rad, 161-0374) of 50 Kd.

Fig S10d

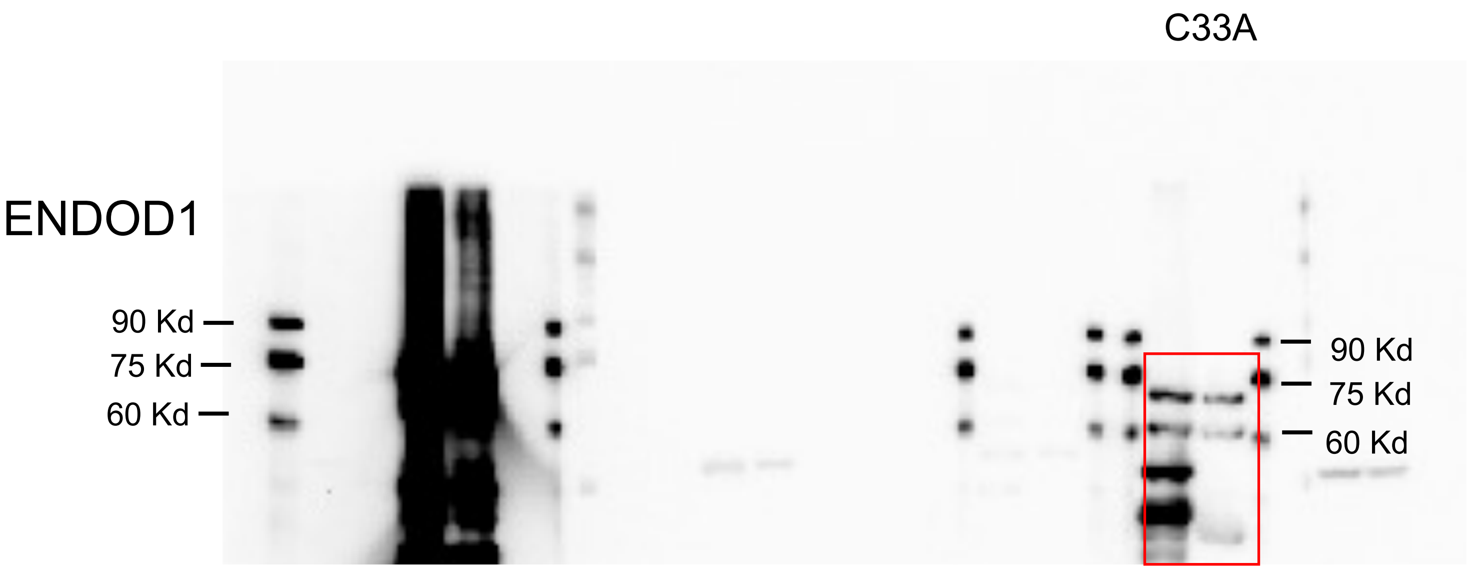
C33A
ENDOD1
90 Kd
75 Kd
60 Kd
90 Kd
75 Kd
60 Kd

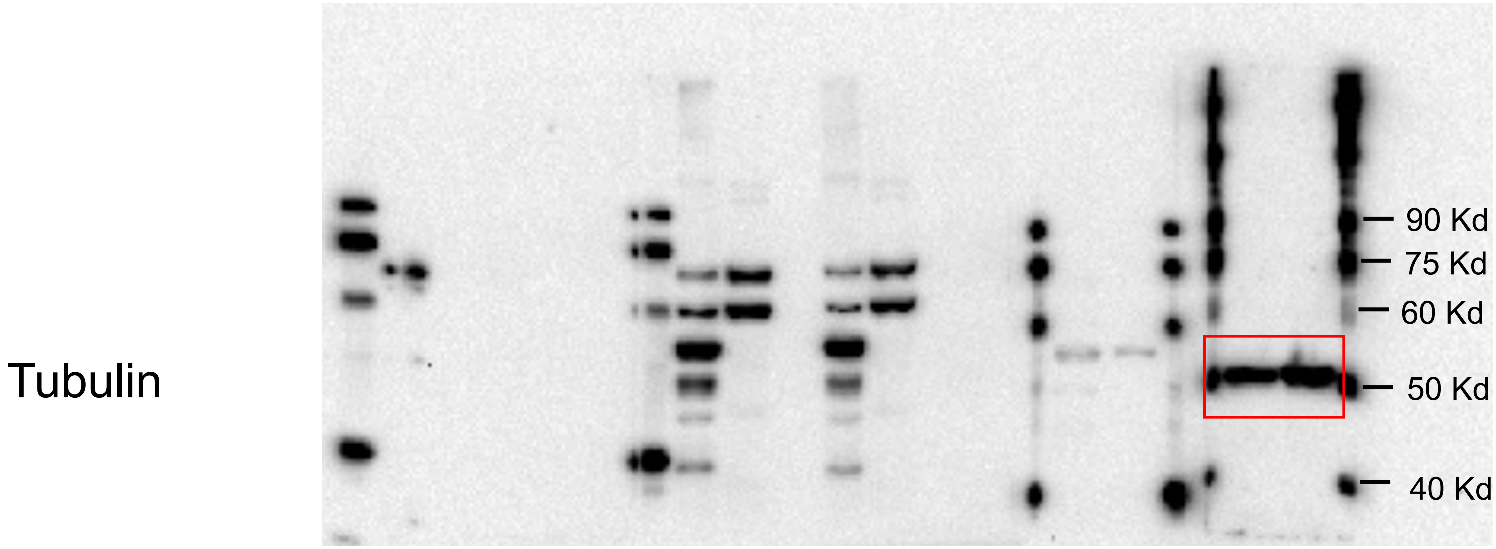
Tubulin
90 Kd
75 Kd
60 Kd
50 Kd
40 Kd

Fig S10d

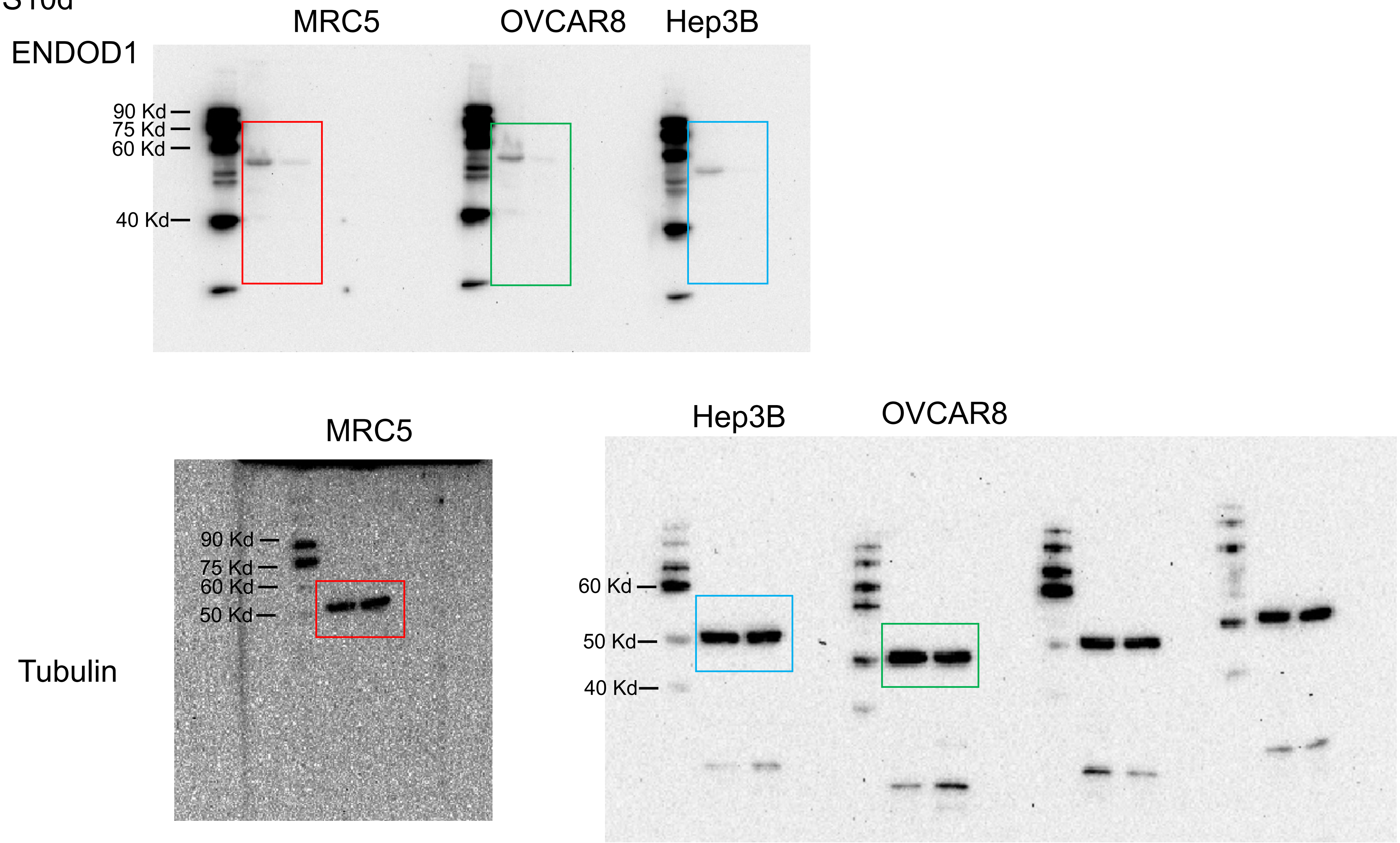
MRC5
OVCAR8
Hep3B
ENDOD1
90 Kd
75 Kd
60 Kd
40 Kd
MRC5
Hep3B
OVCAR8
90 Kd
75 Kd
60 Kd
50 Kd
Tubulin
60 Kd
50 Kd
40 Kd

Fig S10d

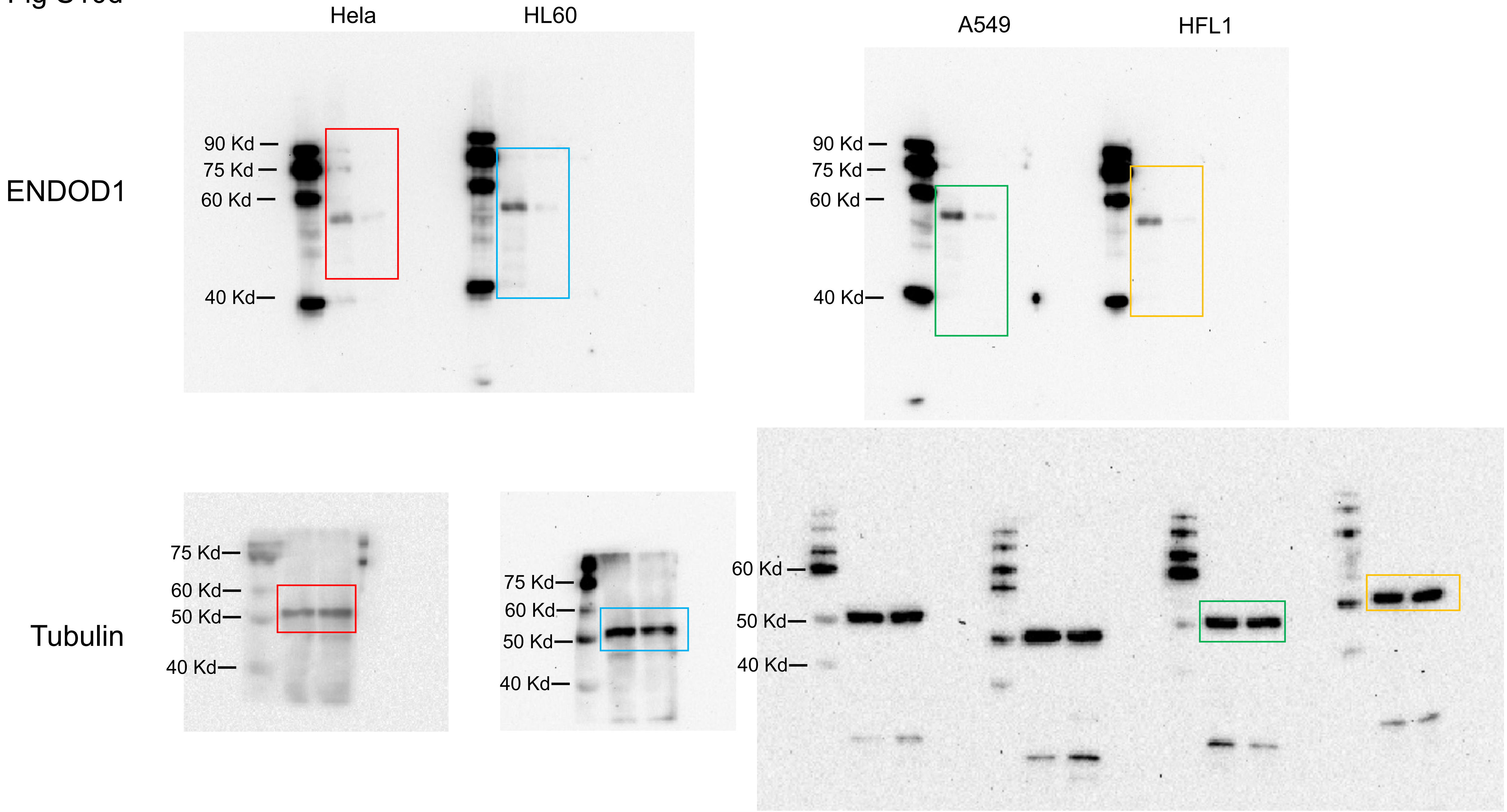

Hela
HL60
A549
HFL1
ENDOD1
90 Kd
75 Kd
60 Kd
40 Kd
90 Kd
75 Kd
60 Kd
40 Kd
Tubulin
75 Kd
60 Kd
50 Kd
40 Kd
75 Kd
60 Kd
50 Kd
40 Kd
60 Kd
50 Kd
40 Kd

**Supplementary Table 1.** Source and genetic background and human cell lines used in this study.

| Cell line | Source | *TP53* gene | p53 protein | HR status |
|---|---|---|---|---|
| RPE1 | Eye epithelial | Wt | Wt | Functional |
| L02 | Liver | Wt | Wt | Functional |
| MRC-5 | Lung fibroblast | Wt | Wt | Functional |
| GES-1 | Gastric | Wt | Wt | Functional |
| IMR-90 | Embryonic lung fibroblast | Wt | Wt | Functional |
| FHs 74 | Intestinal epithelial | Wt | Wt | Functional |
| HFL1 | Embryonic lung fibroblast | Wt | Wt | Functional |
| Wish | Umbilical | Wt | Wt | Functional |
| MCF 10A | Breast epithelial | Wt | Wt | Functional |
| HepG2 | Liver cancer | Wt | Wt | Functional |
| MHCC97L | Liver cancer | TP53 mu | NT | NRD |
| Hep3B | Liver cancer | null | c.(del) | NRD |
| SMMC7721 | Liver cancer | Wt | Wt | NRD |
| HepG2.2.15 | Liver cancer | Wt | Wt | HBV+[1] |
| T43 | Derived from L02 | Wt | Wt | HBV+[1] |
| SKOV3 | Ovarian cancer | null, c.267del | p.P90fs*33 | NRD |
| OVCAR-8 | Ovarian cancer | c.376-1G>A | p.0？ | BRCA1 hypermethylated |
| HO8910 | Ovarian cancer | Wt | Wt | BRCA1 5382 C |
| HO8910PM | Ovarian cancer | Wt | Wt | BRCA1 5382 C |
| C33A | Cervical cancer | homo, c.817C>T | p.R273C | NRD |
| HELA | Cervical cancer | Wt | HPV+[2] | NRD |
| NCI-H1299 | Lung cancer | null | c.(del) | NRD |
| NCI-H1975 | Lung cancer | homo, c.818G>A | p.R273H | NRD |
| A549 | Lung cancer | Wt | Wt | NRD |
| SW480 | Colon cancer | c.925C>T; c.818G>A | p.P309S, p.R273H | NRD |
| COLO-320DM | Colon cancer | homo， c.742C>T | p.R248W | NRD |
| MDA-MB-468 | Breast cancer | homo,c.818G>A | p.R273H | NRD |
| MDA-MB-231 | Breast cancer | homo,c.839G>A | p.R280K | NRD |
| HCC1937 | Breast cancer | homo,c.916C>T | p.R306 | BRCA1, 5382insC |
| MDA-MB-361 | Breast cancer | Wt | Wt | NRD |

| | | | | |
|---|---|---|---|---|
| MCF7 | Breast cancer | Wt | Wt | HRD and PARPi-sensitive |
| U2OS | Sarcoma | Wt | Wt | Functional |
| SH-SY5Y | Neuroblastoma | Wt | Wt | NRD |
| K-562 | Leukemia | homo, c.406 407insC | p.Q136fs*13 | NRD |
| HL-60 | Leukemia | null | c.(del) | NRD |
| Raji | Lymphoma | homo,c.638G>A | p.R213Q | NRD |

NRD: no reported defects.

[1]HBV-positive cells are defective in HR due to impaired DNA end resection[3].

[2]p53 protein in HPV-positive cells are reported to be unstable [2].

**Supplementary Table 2.** Gene functions and gene-specific siRNA used in this study.

| siRNA | Function of genes | Target sequences |
|---|---|---|
| *ENDOD1* | Single strand DNA break repair | GCAAGCGGATTGGCTACAA |
| *TP53* | Cell cycle control and DNA repair | GGAGTATTTGGATGACAG |
| *CHK1* | Check point control | 001: GCAACAGTATTTCGGTATA<br>002: GGAGTATTCTGACTGGAAA<br>003: GAAGCAGTCGCAGTGAAGA |
| *MRE11* | Nuclease for DNA end processing | 001: GCCTCGAGTTATTAAGAAA<br>002: GGATATTGTTCTAGCTAAT<br>003: GGAAATGATACGTTTGTAA |
| *WDR70* | Chromatin remodeling, HR | CTGCCAGAATGGAAGCATA |
| *BLM* | RecQ helicase responsible for Bloom syndrome | 001: GGAAGTGATTTCAGTATTA<br>002: GGAAGAAGCTGAATTACAT<br>003: GAGAAACTCACTTCAATAA |
| *WRN* | RecQ helicase responsible for Werner syndrome | 001: GTAGAAGTTTCTCGGTATA<br>002: GCACCTTCTTACTGAGATA<br>003: GACCAAACCTGTATTTAGA |
| *ARID1A* | ATR-dependent checkpoint control | 001:GAAGCAGGCACCACTAACT<br>002:GCCAGACTCCATATTACAA<br>003:GGCTCACAATGAAAGACAT |
| *ARID1B* | ATR-dependent checkpoint control | 001:CTCCGCAGGTAGAAAGAAA<br>002:CCATGGCGCTTTTATCGAA<br>003:GGCGAAAGATTACCTCCAA |
| *BRCA1* | HR | TCACAGTGTCCTTTATGTA |
| *BRCA2* | HR | 001:GAAGAACAATATCCTACTA<br>002:GCAAAATGTTCTTCAAAGA<br>003:GGTCAAGAATTTCTGTCTA |
| *CTIP* | Nuclease for DNA end processing | 001: GGAATACTCTACAGGAAGA<br>002: GGCCAAAGCACATGGAACA<br>003: CCATGGAGGATGTGAACTT |
| *EXO1* | Nuclease for DNA end processing | 001: GAACGAGTGATTAGTACTA<br>002: CGACAAGCCAATCTTCTTA<br>003: CCTCTTTGCCTGAGAATAA |
| *FANCA* | Intercrosss-link repair, HR | 001: GATCGTGGCTCTTCAGGAA<br>002: GGCCTATGCTAATCATTCT<br>003: GCTCTGCTTTGCAGGATCA |
| *PARP1* | DNA damage sensing, PARylation, SSB and DSB repair | GGAACCAACTCCTACTACA |
| *PARP2* | Ancillary for SSB repair | CCAACACTATAGAAACCTA |
| *PARP3* | | GCATCTACTTTGCCTCAGA |
| *XRCC1* | SSB repair | 001: GGCGGAAACTCATCCGATA |

| | | |
|---|---|---|
| | | 002: GGTTCAGTTTGTGATCACA<br>003: GGCAGACACTTACCGAAAA |
| *TDP1* | Removal of Top1 cleavage complex, SSB repair | 001: GGACCAGTTTAGAAGGATA<br>002: CAAGCACGATCTCTCTGAA<br>003: GAACATTCCTTATGTCAAA |
| *TDP2* | Removal of Top2 cleavage complex, SSB repair | |
| *APTX1* | SSB repair | 001: TGATTCTCCTTGCCTTAAA<br>002: GGACCTCCATTTCCAGTCT<br>003: AGTCAAGGCTTGAAGATTT |
| *DNA Ligase 3* | SSB repair | 001: GGGAAGCCATCTAAGATCA<br>002: GGTGACTTCTCCAGTGAAA<br>003: GCAGCAGGTACACCAAAGA |
| *PNKP* | SSB repair | |
| *POL$\beta$* | SSB repair | 001: GAACCATCATCAGCGAATT<br>002: TCGCAAACTTTGAGAAGAA<br>003: GTGGTGACATGGATGTTCT |
| *XPA* | Nucleotide excision repair | 001: GAAAGACTGTGATTTAGAA<br>002: GGAGACGATTGTTCATCAA<br>003: GACCTGTTATGGAATTTGA |
| *XPE* | Nucleotide excision repair | 001: GAAGCGGCATCTCCTGAAA<br>002: GGTGACTGTTAAATCTGAA<br>003: CCTCCAGGGTGTCTTATAA |
| m*Endod1* | | CATTGACTCTGACTATGAA |
| m*Wdr70* | | 001: GGCCAAGGTTATTGACAGA<br>002: GAAUGAACCAGAAUGGAAA<br>003: GGAUAUGAUUACGAUGUUA |

**Supplementary Table 3.** Primers used in this study.

| | | |
|---|---|---|
| Primer 1 | GACGACGATAAGGAATTCATGGAGGAGCCGCAGTCAG | TP53 to pLVX-Flag-IRES-ZsGreen1, forward |
| Primer 2 | GCTCTAGAACTAGTCTCGAGTCAGTCTGAGTCAGGCCC | TP53 to pLVX-Flag-IRES-ZsGreen1, reverse |
| Primer 3 | GAAACATTTTCAGACCAATCAAAACTACTTCCTGAAAAC | TP53 LW22/23QS, forward |
| Primer 4 | GTTTTCAGGAAGTAGTTTTGATTGGTCTGAAAATGTTTC | TP53 LW22/23QS, reverse |
| Primer 5 | CGTCTGGGCTTCTTGAATTCTGGGACAGCCAAG | TP53 H115N, forward |
| Primer 6 | CTTGGCTGTCCCAGAATTCAAGAAGCCCAGACG | TP53 H115N, reverse |
| Primer 7 | GACGGAGGTTGTGAGGCACTGCCCCCACCATGAG | TP53 R175H, forward |
| Primer 8 | CTCATGGTGGGGGCAGTGCCTCACAACCTCCGTC | TP53 R175H, reverse |
| Primer 9 | GCCCCTCCTCAGCATTTTATCCGAGTGGAAG | TP53 L194F, forward |
| Primer 10 | CTTCCACTCGGATAAAATGCTGAGGAGGGGC | TP53 L194F, reverse |
| Primer 11 | CATGGGCGGCATGAACCAGAGGCCCATCCTCACC | TP53 R248Q, forward |
| Primer 12 | GGTGAGGATGGGCCTCTGGTTCATGCCGCCCATG | TP53 R248Q, reverse |
| Primer 13 | CATGGGCGGCATGAACTGGAGGCCCATCCTCACC | TP53 R248W, forward |
| Primer 14 | GGTGAGGATGGGCCTCCAGTTCATGCCGCCCATG | TP53 R248W, reverse |
| Primer 15 | GAACAGCTTTGAGGTGTGTGTTTGTGCCTGTCCTG | TP53 R273C, forward |
| Primer 16 | CAGGACAGGCACAAACACACACCTCAAAGCTGTTC | TP53 R273C, reverse |
| Primer 17 | GAACAGCTTTGAGGTGCATGTTTGTGCCTGTCCTG | TP53 R273H, forward |
| Primer 18 | CAGGACAGGCACAAACATGCACCTCAAAGCTGTTC | TP53 R273H, reverse |
| Primer 19 | GTGCCTGTCCTGGGAAAGACCGGCGCACAGAG | TP53 R280K, forward |
| Primer 20 | CTCTGTGCGCCGGTCTTTCCCAGGACAGGCAC | TP53 R280K, reverse |
| Primer 21 | GAGATGTTCCGAGAGCCGAATGAGGCCTTGGAAC | TP53 L344P, forward |

| | | |
|---|---|---|
| Primer 22 | GTTCCAAGGCCTCATTCGGCTCTCGGAACATCTC | TP53 L344P, reverse |
| Primer 23 | GCTCTAGAACTAGTCTCGAGTCACCTGCTCCCCCCT GGCTC | TP53 del364-393, reverse |
| Primer 24 | GGATCTATTTCCGGTGAATTCATGGGCACCGCGCGCT GGC | ENDOD1 to pLVX-IRES-ZsGreen1, forward |
| Primer 25 | CGCTCTAGAACTAGTCTCGAGTTACTTATCGTCGTCAT CCTTGTAATCTAACTCCCCAGAATTGTCAAAAG | ENDOD1-Flag to pLVX-IRES-ZsGreen1, reverse |
| Primer 26 | GATGACGACGATAAGGAATTCATGGGCACCGCGCGC TGGC | ENDOD1 to pLVX-Flag-IRES-ZsGreen1, forward |
| Primer 27 | CGCTCTAGAACTAGTCTCGAGTTATAACTCCCCAGAA TTGTCAAAAG | ENDOD1 to pLVX-Flag-IRES-ZsGreen1, reverse |
| Primer 28 | GATGACGACGATAAGGAATTCATGCGGCTCGTGGGC GAGGAGG | ENDOD1(22-500) to pLVX-Flag-IRES-ZsGreen1, forward |
| Primer 29 | CGCTCTAGAACTAGTCTCGAGTTATTGAAAAAGCTTG ATGAATGGGG | ENDOD1(22-344) to pLVX-Flag-IRES-ZsGreen1, reverse |

**Supplementary Table 4.** Antibodies used in this study

| Antibodies | Source and dilutions | Dilution |
|---|---|---|
| Rabbit anti-phosphor-Serine 33, RPA32 | NOVUS, NB100-544 | 1:1000 |
| Mouse anti-α-Tubulin | Sigma, T6074 | 1:5000 |
| Mouse anti-phospho Serine 139, gH2AX | Millipore, 05-636 | 1:500 |
| Rabbit anti-53BP1 | Bethyl, A300-272A | 1:1000 |
| Rabbit anti-phosphor-Serine 1524, BRCA1 | Bethyl, A300-001A | 1:500 |
| Rabbit anti-BRCA1 | Huabio, HA500015 | 1:500 |
| Mouse anti-PAR | Abcam, ab14459 | 1:500 |
| Rabbit anti-MRE11 | CST, 4895S, | 1:500 |
| Rabbit anti-CTIP | NOVUS, NB100-79810 | 1:200 |
| Rabbit anti-BLM | Bethyl, A300-110A-2 | 1:200 |
| Rabbit anti-phospho-Serine 343, NBS1 | Abcam, ab109453 | 1:1000 |
| Rabbit anti-NBS1 | Huabio, ET1610-26 | 1:1000 |
| Rabbit anti-XRCC1 | Huabio, ET1704-01 | 1:500 |
| Mouse anti-BrdU | Roche, 11585860001 | 1:500 |
| Rabbit anti-FANCD2 | NOVUS, NBP2-57171 | 1:500 |
| Rabbit anti-Flag | Custom made | 1:1000 |
| Rabbit anti-p53 | HUABIO, ET1601-13 | 1:1000 |
| Rabbit anti-PARP1 | HUABIO, ER1802-67 | 1:500 |
| Rabbit anti-PARP2 | HUABIO, ET7108-05 | 1:500 |
| Rabbit anti-PARP3 | NOVUS, NBP2-49523 | 1:500 |
| Mouse anti-H3 | Millipore, 05-1341 | 1:20000 |
| Rabbit anti-ENDOD1 | ABcam, ab121293 | 1:1000 |
| Rabbit anti-ENDOD1 | ABclonal, A16502 | 1:1000 |
| HRP-conjugated anti-mouse IgG | DAKO, P0260 | 1:3000 |
| HRP-conjugated anti-rabbit IgG | DAKO, P0448 | 1:3000 |
| FITC-conjugated anti-mouse IgG | Sigma, F0257 | 1:300 |
| CY3- conjugated anti- rabbit IgG | Sigma, C2306 | 1:300 |
| B220 (Efluor 450) | Invitrogen, 48-0452-82 | 1 ul /$10^6$ cells |
| CD3e (PE-cy7) | Invitrogen, 25-0031-82 | 1 ul /$10^6$ cells |
| Ly-6G/Gr-1 (APC) | Invitrogen, 17-9668-82 | 1 ul /$10^6$ cells |
| F480 (FITC) | Biolegend, B257636 | 1 ul /$10^6$ cells |
| CD11b (APC) | Biolegend, B261578 | 1 ul /$10^6$ cells |
| NK1.1 (PE-cy7) | Invitrogen, 25-5941-82 | 1 ul /$10^6$ cells |
| Lineage (Efluor 450) | Invitrogen, 88-7772-72 | 1 ul /$10^6$ cells |
| Sca-1 (PE) | Invitrogen, 12-5981-83 | 1 ul /$10^6$ cells |
| c-Kit (APC) | invitrogen, 17-1171-82 | 1 ul /$10^6$ cells |
